# Giant polyketide synthase enzymes biosynthesize a giant marine polyether biotoxin

**DOI:** 10.1101/2024.01.29.577497

**Authors:** Timothy R. Fallon, Vikram V. Shende, Igor H. Wierzbicki, Robert P. Auber, David J. Gonzalez, Jennifer H. Wisecaver, Bradley S. Moore

## Abstract

*Prymnesium parvum* are harmful haptophyte algae that cause massive environmental fish-kills. Their polyketide polyether toxins, the *prymnesins*, are amongst the largest nonpolymeric compounds in nature, alongside structurally-related health-impacting “red-tide” polyether toxins whose biosynthetic origins have been an enigma for over 40 years. Here we report the ‘PKZILLAs’, massive *P. parvum* polyketide synthase (PKS) genes, whose existence and challenging genomic structure evaded prior detection. PKZILLA-1 and -2 encode giant protein products of 4.7 and 3.2 MDa with 140 and 99 enzyme domains, exceeding the largest known protein titin and all other known PKS systems. Their predicted polyene product matches the proposed pre-prymnesin precursor of the 90-carbon-backbone A-type prymnesins. This discovery establishes a model system for microalgal polyether biosynthesis and expands expectations of genetic and enzymatic size limits in biology.

## Introduction

Large-scale fish deaths caused by harmful algal blooms are global health, environmental, and food security problems (*1*). Anthropogenic causes continue to hasten the severity and frequency of toxic blooms in freshwater and marine ecosystems, including the massive fish kill along the Odra River in 2022 by the golden alga *Prymnesium parvum* (Haptophyta) that decimated half of the river’s fish population through Poland and Germany (*2*). Oceanic blooms of the red-tide algae *Karenia brevis* (Dinoflagellata), experienced annually off US southeastern coastlines, are similarly devastating to fish and marine mammals (*3*). Their respective poisons, prymnesin and brevetoxin, are just two of many notable examples of marine microalgal biotoxins that share giant, polycyclic polyether structures that are amongst the largest nonpolymeric carbon chain molecules in nature (*4*).

The massive and stereochemically-rich microalgal polyketide biotoxins prymnesin-1 (Fig. 1A) (*5*), palytoxin (*6*), and maitotoxin (*7*) contain 90, 115, and 142 contiguous carbon atoms, respectively, and pose significant human and environmental health risks. Their chemical structures imply a biosynthetic assembly line construction of two-carbon chain length iterations to a polyene intermediate that undergoes epoxidation followed by a nucleophilic reaction cascade to assemble their distinctive trans-fused (“ladder-frame”) polyether frameworks (*8, 9*). However, the biosynthesis of these massive microalgal toxins has remained an enigma despite a wealth of intimate knowledge of polyketide biochemistry from decades of research in bacteria and fungi (*10*) and recent transcriptomic studies identifying biosynthetic gene candidates in multimodular type I polyketide synthases (PKSs) from toxic microalgae (*11, 12*). The sheer size of microalgal polyether biotoxins present significant experimental challenges and are accompanied by a lack of methods to study their genetic origin. The model green alga *Chlamydomonas reinhardtii*, for instance, hosts a single large ∼80 kbp PKS, known as PKS1 (Fig. 1B), and while genetic knockout experiments established that PKS1 participates in formation of the zygospore cell wall, its polyketide product remains unknown (*13*). Furthermore, unlike bacteria and fungi that organize their PKS encoding genes into polycistrons and biosynthetic gene clusters (BGCs), other eukaryotes typically employ monocistronic mRNAs and infrequently functionally co-localize most genes, thus greatly obfuscating gene discovery efforts (*14*).

**Fig. 1.**
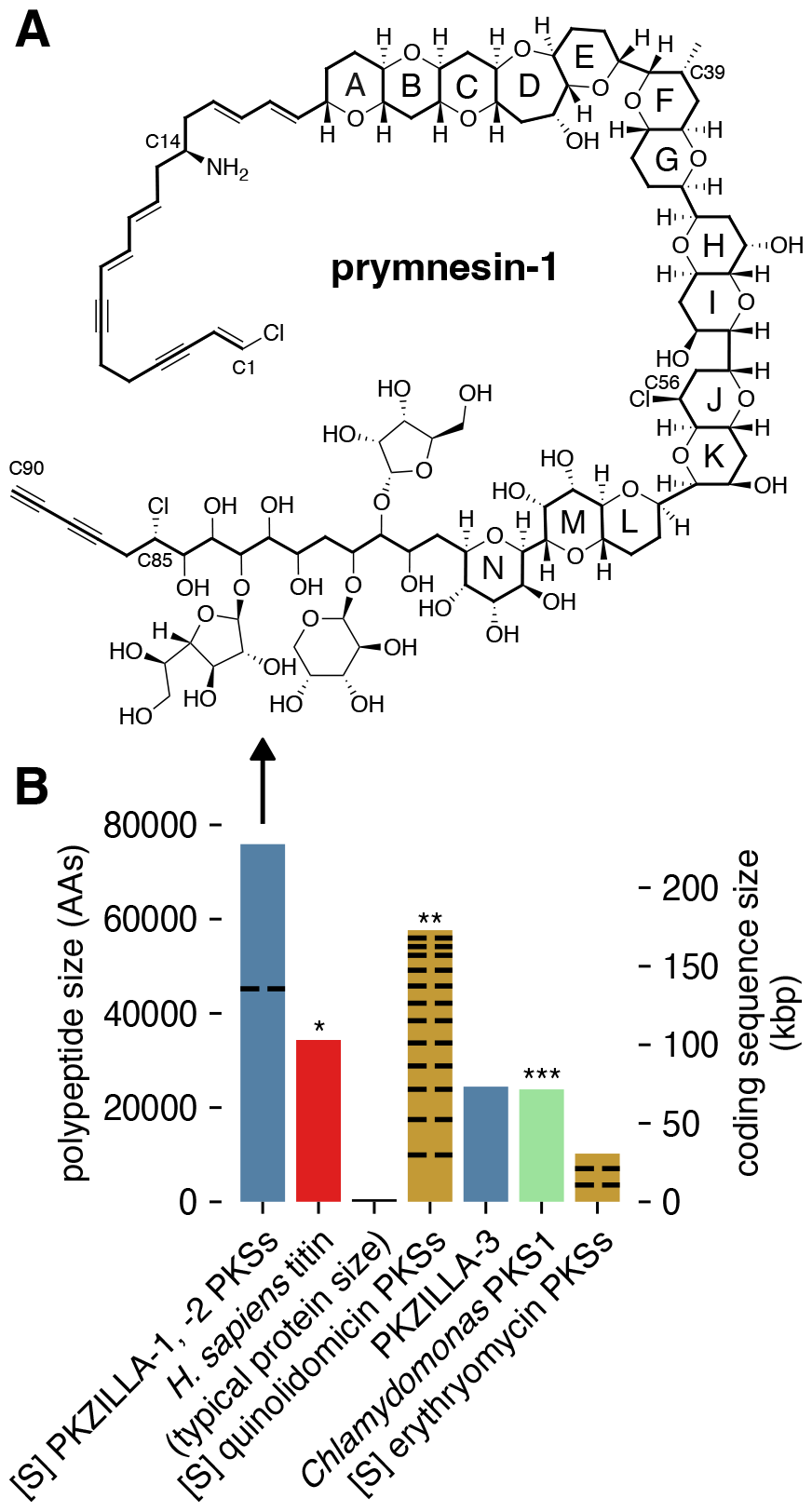
Prymnesin, its source PKZILLA polyketide synthases (PKSs), and other large proteins and PKS systems. **(A)** Molecular structure of prymnesin-1 (*16*). **(B)** Comparison of polypeptide and coding nucleotide sizes from representative PKSs or computationally summed PKS systems. Blue=PKZILLAs from *P. parvum* 12B1 (this work). [S]=Computationally summed lengths for independent PKS proteins that participate in the same biosynthetic system. Black dashed lines=Divisions of PKS systems into independent proteins. Red/*=largest known protein (non-PKS) (*15*). Gold=Representative bacterial PKS systems, including the quinolidomicin (**=previous largest known PKS system) (*17*) and erythromycin (*18*) PKSs. Green/***=previous largest genetically studied microalgal PKS (*13*).

Herein we report the application of a customized gene annotation strategy which enabled the discovery of two massive PKS genes, PKZILLA-1 and -2, from *P. parvum* strain 12B1 that we propose are responsible for the complete backbone assembly of its notorious ladder-frame polyether toxin, prymnesin-1. Not only are the two giant PKZILLA “gigaproteins” organized consistently with the long-anticipated polyene intermediate structure of a microalgal ladder-frame polyether, but PKZILLA-1 is larger than titin (*15*), the presently largest known protein in life (Fig. 1B).

## Results

### Genomic and transcriptomic evidence for the PKZILLA gigasynthases

We selected the A-type prymnesin (*19*) producing *P. parvum* strain 12B1 as a model system to resolve microalgal polyether biosynthesis, as its 116 Mbp genome and our recently published near-chromosome-level genomic assembly (*20*) makes strain 12B1 relatively tractable amongst microalgae and other toxic *P. parvum* strains. By contrast, polyether toxin-producing marine dinoflagellates have genomes ∼100X larger at 25+ Gbp with extreme tandem gene repeat structures (*21, 22*) that have prevented even draft genome assemblies. We first cataloged PKS genes potentially involved in prymnesin biosynthesis within our automated gene annotation (*20*), identifying 44 PKS genes encoding relatively small proteins with 1-3 *trans*-acyltransferases (*trans*-AT) PKS modules. However, confirmatory tblastn queries using PKS domains unveiled three seemingly contiguous and strikingly large PKS “hotspot” loci that stood apart at 137, 93, and 74 kbp on pseudo-chromosomes 17, 7, and 10, respectively. These hotspots showed a high concentration of visible coding regions that were only partially captured by 25 fragmented PKS gene models, thus we hypothesized they represented massive and mis-annotated single-genes. Upon manual revision, we successfully constructed single-gene models from each hotspot, as described below, and dubbed the resulting genes PKZILLA-1, -2, and -3 (Fig. 1, Fig. 2, table S1). At final count, we annotated 22 PKS genes distributed across 16 of 34 pseudo-chromosomes that dramatically ranged in size from 3 to 137 kbp (table S1).

**Fig. 2.**
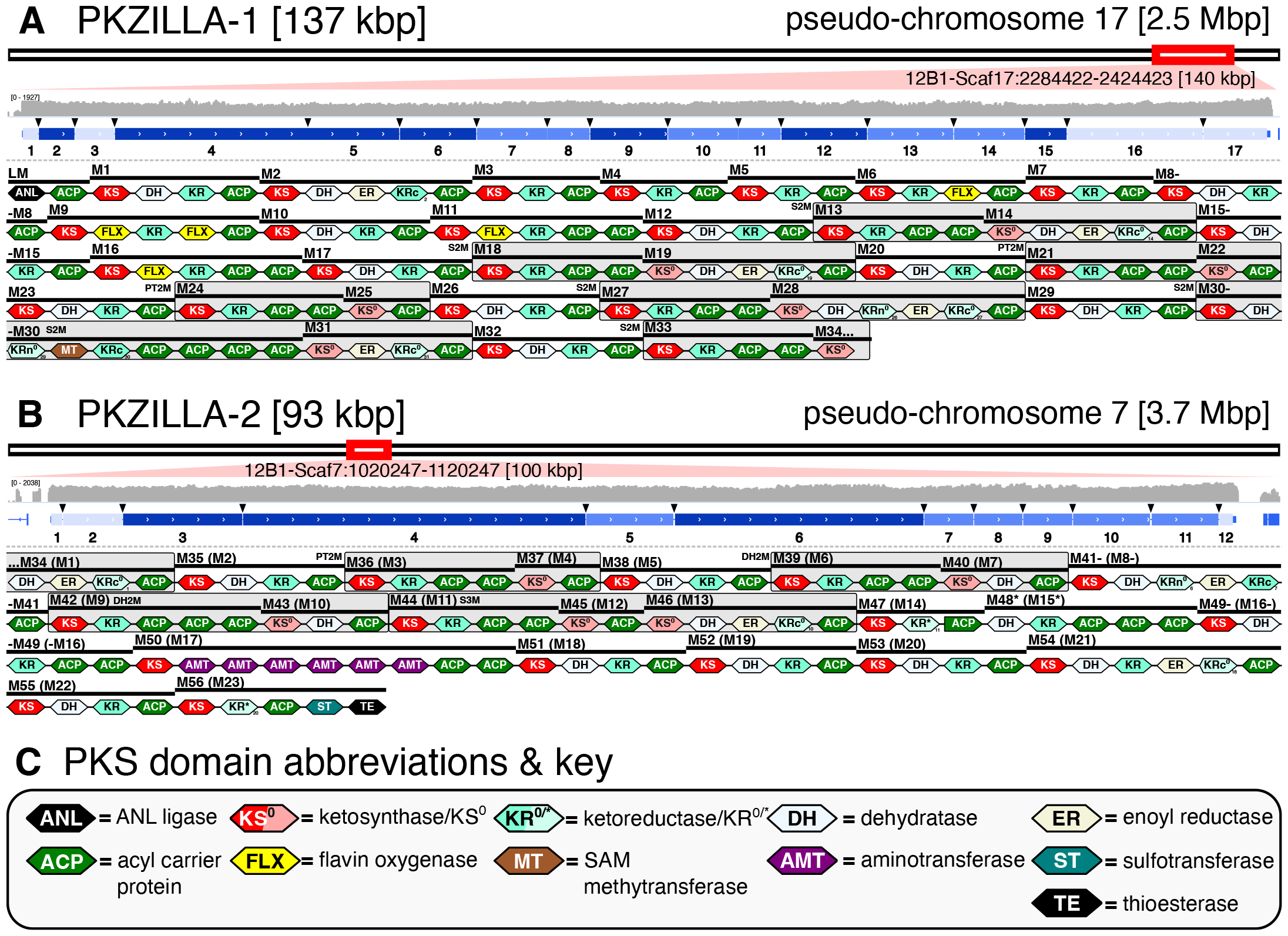
Genomic, transcriptomic, and proteomic evidence for the PKZILLAs. Genomic PKS hotspot loci with gene models and PKS domain and module annotations for **(A)** PKZILLA-1 and **(B)** PKZILLA-2. **(A, B)** Red boxes denote chromosomal locations and relative sizes of the PKZILLA genes. The contiguous log-scale forward-stranded read coverage from the stranded rRNA-depletion RNA-Seq (in gray) is shown across the PKZILLA gene models (in blue). Introns are highlighted with black arrows, while exons are numbered 1-17 for PKZILLA-1 and 1-12 for PKZILLA-2. See fig. S2 for an alternative view and fig. S3, S4 for a detailed view of each intron. The numbered protein-coding exons are colored light blue, medium blue, or dark blue based on whether supporting proteomic peptides from that exon were not detected, detected by protein-multimatch peptide matches alone, or detected by protein-unique plus exon-unique peptide matches, respectively (See *Proteomic evidence for the PKZILLAs* in Results; fig. S9). Domain and module annotations (starting with the loading module (LM) and module 1 (M1) of PKZILLA-1 and ending with M56 of PKZILLA-2) are shown below the gene models, see key in **(C)**. The bi/tri-modules are boxed in gray and categorized as S2/3M=saturating bi/tri-module, PT2M=pass-through bimodule, and DH2M=dehydrating bimodule. See fig. S6, S7 for non-length-normalized domains.

Constructing the PKZILLA gene models from their candidate hotspots required several manual gene annotation interventions. Initial realignments of our Oxford Nanopore Technologies (ONT) long genomic DNA reads (*20*) to the PKZILLA hotspots revealed an assembly collapse of a tandem repetitive region within the coding N-terminus of PKZILLA-1 that was fixed by targeted reassembly. After that revision, we found no further PKZILLA assembly concerns (fig. S1). To test for and classify transcriptional activity at these hotspots, we next analyzed coverage from a *P. parvum* 12B1 oligo-dT mRNA enrichment/poly-A tail pulldown Illumina RNA sequencing (RNA-seq) dataset (*23*) to localize the putative PKZILLA mRNA 3’-ends. This dataset indicated one transcriptional termination site (TTS) per hotspot (fig. S2), however as established for poly-A pulldown RNA-Seq datasets, the coverage was negligible beyond ∼10 kbp to the 5’-end and thus uninformative to 5’ transcriptional activity (fig. S2). To evaluate if the full PKZILLA hotspots were transcriptionally active and consistent with single genes, we applied mRNA length-unbiased rRNA-depletion Illumina RNA-seq by generating and sequencing two dUTP-stranded libraries from exponentially growing *P. parvum* cultures from the day and night phases. The low relative expression of the PKZILLAs required four independent sequencing runs to accumulate sufficient coverage. Ultimately, we calculated that the PKZILLA transcript expression levels were uniformly low with transcripts per million (TPM) values of 1, 2, and 0.5 for PKZILLA-1, -2, and -3, respectively, in both day and night phases (table S1). These rRNA datasets further showed contiguous and sense-stranded transcriptional activity across the three PKZILLA gene models indicating one transcriptional start site (TSS) per PKZILLA hotspot (Fig. 2, fig. S2). Critically, the rRNA depletion data identified the presence and location of the 34 PKZILLA introns, all of which showed canonical eukaryotic GT-AG splice sites (table S2, fig. S3, S4, S5). The translated PKZILLA polypeptide sequences show near-contiguous sequence similarity to known PKS domains, with limited evidence for internal breaks (fig. S6, S7). Thus, we concluded that PKZILLA-1, -2, and -3 are single genes that each encode a single major transcriptional and translational product (Fig. 2). Remarkably, the calculated size of the PKZILLA-1 transcript at 136,071 nucleotides and the associated protein at 45,212 amino acids would make it about 25% larger than the presently largest known protein, the mammalian muscle protein titin (*15*) (Fig. 1B, table S5).

### Proteomic evidence for the PKZILLAs

To validate the PKZILLA proteins predicted by our gene models, we analyzed lyophilized *P. parvum* 12B1 biomass using an optimized bottom-up proteomics method. We identified and confidently validated 43 and 38 proteomic peptides from PKZILLA-1 and -2, respectively, yet none for PKZILLA-3. Only 9 and 6 peptides from PKZILLA-1 and -2, respectively, were single-copy (*single-match*) within a single predicted PKZILLA polypeptide (*protein-unique*). Instead, most of the detected peptides were *multimatch peptides* present in multiple copies, either *protein-unique* to a given PKZILLA, or present in both PKZILLA-1 and -2 polypeptides (*protein-multimatch*) (fig. S8, S9). This high proportion of multimatch peptides highlights the internally repetitive nature of the “giga-modular” PKZILLAs, both within and across proteins. These peptides were only present in the PKZILLA gene models and were not found anywhere else in 6-frame translations of the 12B1 genome.

We next established which regions of the PKZILLA polypeptides were supported by proteomics. A complication is that most of the *P. parvum* proteomic data were multimatch peptides, which are rare in proteomic analyses of typical non-large, non-repetitive proteins. Since they cannot be unambiguously assigned to a single polypeptide region, multimatch peptides are often ignored in downstream analyses, in favor of simpler protein-unique single-match peptides (*24*). We judged that overlooking the multimatch peptides, while a simple solution, needlessly limited our analysis and discarded valuable data. We adapted to this challenge by sub-classifying each protein-unique yet multimatch peptide that also only arose from the translation product of a single exon as *exon-unique* (fig. S9), thus localizing proteomic support to the *exon* rather than the residue level. Of the 43 PKZILLA-1 proteomic peptides, 14 met both the protein-unique and exon-unique criteria, and thus established unambiguous proteomic support for translation of 7 out of 17 PKZILLA-1 exons (41%), bounded upstream and downstream by exons 2 and 15, respectively (Fig. 2). When considering the remaining 29 tryptic peptides despite their exon-multimatch or protein-multimatch ambiguity (fig. S9), we established increased proteomic support for 76% of the PKZILLA-1 exons (Fig. 2). Applying the same criteria to PKZILLA-2, we measured proteomic support of 16 exon-unique peptides from 3 of the 12 exons (25%), bounded by exons 3 and 6, which increased to 75% of PKZILLA-2 exons after considering all PKZILLA matching peptides (Fig. 2). Overall, these results confidently validate the translation of the PKZILLA-1/-2 transcripts into proteins and are consistent with a single translational product per gene.

### Annotation of PKZILLA domain & modular structures and their compatibility with prymnesin

With proteomics-validated PKZILLA gene models in hand, we next tested their possible role in prymnesin biosynthesis by annotating the PKS domains and evaluating their modular-arrangement against the chemical structure of a proposed pre-prymnesin biosynthetic precursor (PPBP; Fig. 3A). We identified 140 and 99 protein domains for PKZILLA-1 and -2, respectively (table S3, S4, S5), using InterProScan (*25*). We also cataloged 30 candidate domains of unknown function (cDUFs), however none of these cDUFs showed strong evidence of being unannotated enzyme domains (table S6, S7). The first two domains of PKZILLA-1, an <u>A</u>cyl-CoA synthetase/<u>N</u>RPS adenylation domain/<u>L</u>uciferase (ANL) superfamily ligase adjacent to an acyl-carrier-protein (ACP) domain (Fig. 2), comprise an unconventional, yet precedented (*26*), loading module (LM) to initiate polyketide chain elongation. PKZILLA-2 lacked any recognizable N-terminal loading domains; however, it does possess a C-terminal thioesterase (TE) domain, consistent with polyketide chain termination. In the end, we organized the combined 239 domains into 56 *trans*-AT PKS modules, including module-34 (M34) split across the C-terminus of PKZILLA-1 and the N-terminus of PKZILLA-2 (Fig. 2, table S5, S8, S9).

**Fig. 3.**
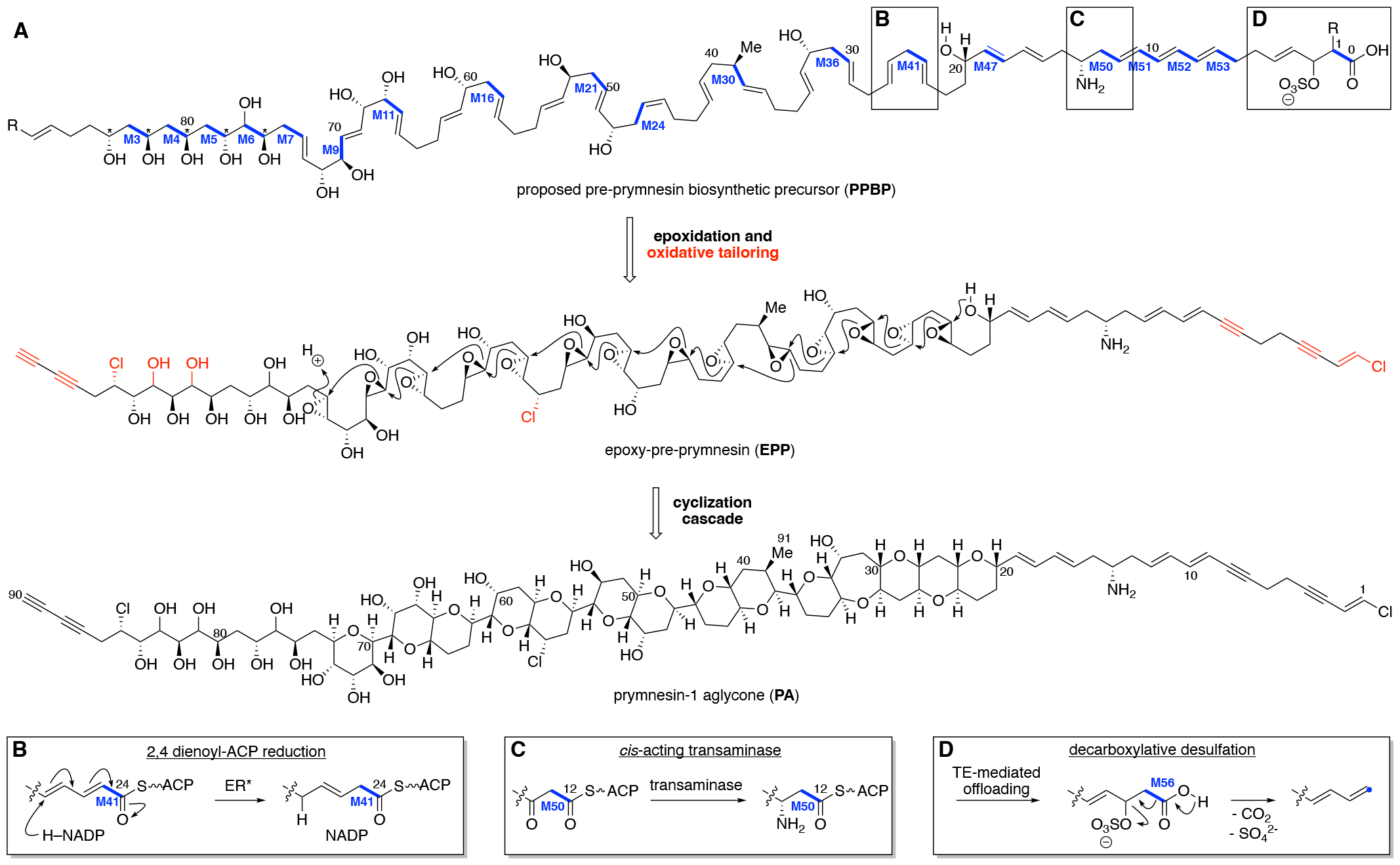
Alignment of PKZILLA PKS modules with the proposed prymnesin biosynthetic precursor. **(A)** Structure of the pre-prymnesin biosynthetic precursor (PPBP) inferred from retrobiosynthetic analysis of a structurally compatible poly-epoxide cyclization cascade from the prymnesin-1 aglycone (PA) (*16*), and its end-to-end alignment with the assembly line modules of the PKZILLA-1/-2 gigasynthase. Select hypothesized reactions are called out in subpanels **(B, C, D)**. See fig S20 for further detail.

Prymnesin is devoid of standard polyketide termini and had an unknown direction of chain elongation. We resolved its biosynthetic directionality by first correlating the diagnostic polyol segment at C84–C76 with the five adjacent modules M3–M7 that contain a ketoreductase (KR) as the terminal reductive domain. Based on this observation, we could infer the directionality of biosynthesis, with the PKZILLA-1 LM initiating the polyketide biosynthetic pathway with a three-carbon carboxylate of yet unknown origin followed by chain extension with seven malonate molecules via M1–M7 with interceding KR reduction to produce a hexol intermediate (fig. S20). While five of the “β”-hydroxyls originate from malonate extender units, the out-of-sequence “α”-hydroxy group at C77 is instead likely installed by the α-hydroxylase flavoprotein (FLX) domain (*27, 28*) contained in M6. PKZILLA-1 harbors two additional FLX-domain containing modules, M9 and M11, both of which align with the installation of the further α-hydroxylations at C71 and C67, respectively. The absence of domains corresponding to additional C–H oxygenations (C81 and C83) and halogenation (C85) is suggestive that these functionalities are installed by intermodular- or *trans*-acting enzymes during chain elongation or post polyketide assembly via oxidative enzymes (*29*).

The PPBP region spanning prymnesin’s polyether rings (C74–C20) coincides with a break from canonical *trans*-AT modular architecture with 11 non-elongating ketosynthase (KS^0^) containing modules interspersed amongst 16 canonical *trans*-AT PKS modules (Fig. 2, 3A, table S8, S9). Two of the KS^0^s are found as part of “dehydrating bimodules” (Fig. 2; M39/M40, M42/M43) first characterized in bacterial *trans*-AT pathways (*30*), wherein the first module (KS-KR(-ACP_n_)) catalyzes chain elongation and ketone reduction, and the second module, consisting of a (KS^0^-DH-ACP), performs the corresponding dehydration to yield an α,β-alkene thioester intermediate (fig. S20). Three other KS^0^s are integrated into simple “pass-through bimodules” (Fig 2.; M21/M22, M24/M25, M36/M37), also found in bacterial *trans*-AT BGCs (*31*), whose minimal KS^0^-ACP architecture preserves the β-hydroxy group generated by the upstream module. The remaining KS^0^s reside in unprecedented “saturating” bi- and trimodules (Fig. 2; M13/M14, M18/M19, M27/M28, M30/M31, M33/M34, and M44/M45/M46). In these systems, the second and third modules contain the full complement of reductive domains which convert the β-hydroxy group to a saturated methylene at positions C64, C56, C44, C40, C36, and C22 (Fig. 3A, fig. S20). Notably, saturating bimodule M30/M31 also contains the only methyltransferase (MT) domain in the entire assembly line and is positioned to install prymnesin’s lone methyl group at C39 (Fig. 3, fig. S20). We further confirmed the module-to-precursor alignment throughout this C74–C20 region by applying *trans*-AT ketoreductase precedent (*32*) to bioinformatically predict the stereochemical outcome of reduction for each KR domain (table S11). These bioinformatic predictions matched with 6 of the 7 known configurations from the most recent structure revision of prymnesin-1 (*16*), with the C32 hydroxyl as the exception (table S12, fig. S20). By extrapolating these predictions, we propose β-hydroxy stereochemical assignments for the yet unassigned C84–C76 region of prymnesin (Fig. 3A, table S12).

Finally, the polyene segment (C19–C1) contains several distinguishing structural features, all of which align with the final ten modules of PKZILLA-2. The M49 dehydratase is positioned to catalyze a precedented (*33*) vinylogous dehydration to reconfigure the C19–C16 diene out of conjugation relative to the ACP-tethered thioester, and the six consecutively arranged pyridoxal 5’-phosphate (PLP) dependent aminotransferase (AMT) domains in M50 are located at the precise position to incorporate the sole primary amine at C14 (Fig. 3D) as in mycosubtilin biosynthesis (*34*). Much like the initial modules in PKZILLA-1, the final modules in PKZILLA-2, M51–M55, possess traditional *trans*-AT domain architecture, and generate the C12–C7 triene and a transient C4–C3 alkene that must undergo further desaturation to give prymnesin’s observed alkyne. The final module, M56, is precedented by the terminal PKS module from curacin biosynthesis (*35*) wherein an unusual sulfotransferase (ST) domain sulfates the β-hydroxy, and TE-mediated offloading initiates simultaneous decarboxylation and sulfate elimination to give a terminal alkene (Fig. 3D). As prymnesin terminates in a vinyl chloride, an additional halogenase must act pre- or post-chain offloading to install the third and final chloride at C1 (*36*).

While much of the prymnesin assembly line conforms to *trans*-AT PKS biochemistry, there are a few unique module-to-precursor alignments that may signal new enzymology. In the case of M40 and its dienoyl intermediate, we propose that the adjacent M41 module elongates the growing polyketide chain to a 2,4-dienoyl-ACP intermediate before reduction (Fig. 3B, S20) by a phylogenetically distinct enoyl reductase “ER2” (table S9, fig. S14, S19) to generate a β,γ-alkene out of conjugation with the thioester carbonyl. This type of reduction has precedence in fatty acid biosynthesis on acyl-CoA intermediates (*37*). Furthermore, while we anticipate the KRs from M47 and M56 are active per our model (Fig. 3, S20), these domains are phylogenetically distinct (fig. S12, S17) and show unprecedented active site residues (table S9) that may suggest a novel mechanism or substrate. Similarly, module M48 is unique in missing an explicit KS domain (Fig. 2, S7, table S9), and instead harbors the 186-residue sequence-unique cDUF9 in the analogous upstream position (table S7, S9) that may help recruit a restorative *cis*- or *trans*-acting KS enzyme (Fig. 3A). Finally, based on the currently revised structure of prymnesin-1 (*16*), our biosynthetic model requires a (*Z)*-ene at C45–C46 to accommodate the stereochemical outcome of the prymnesin aglycone, despite the M26-DH/KR pair predicting an (*E)*-ene product (table S11). This may suggest further refinement of prymnesin-1’s configuration is warranted (*16*). Taken together, the sequence of the assembly line operation strongly supports a causal role for PKZILLA-1/2 as the “gigasynthase” responsible for synthesis of the prymnesin backbone, and suggests future focus areas to identify the remaining biosynthetic enzymes for the polyether cascade.

## Discussion

The characterization of brevetoxin B as the causative agent of the toxic red-tide dinoflagellate *Karenia brevis* over four decades ago (*38*), established microalgae as exquisite producers of polycyclic polyether toxins. Over 150 members with their five- to nine-membered cyclic ethers, including maitotoxin as the largest with 32 fused rings, have since been discovered (*39*) and have helped shape the field of marine natural products due to their toxic and drug-like properties (*40*). The discovery and initial characterization of the prymnesin PKZILLA gigasynthases now sheds light on the long-standing question about how microalgae biosynthesize their giant polyketide biotoxins.

The domain composition of the PKZILLA-1 and -2 gigasynthases are impeccably aligned with the cooperative assembly of the prymnesin carbon scaffold, and support decades-old hypotheses that ladder-frame polyethers are constructed from all (*E*)- linear polyene intermediates (*8, 9*). Our discovery identified several unexpected features of the prymnesin assembly line. The sizes of the PKZILLA enzymes are stunning, with the 140-domain protein PKZILLA-1 being larger than the presently largest recognized protein in life. Second, only two modular PKS proteins are required for the construction of the 90-carbon long prymnesin molecule. In contrast, the longest known bacterial polyketide, quinolidomicin at 68 carbons (*17*), is assembled by 13 PKS gene products (Fig. 1B). The remarkable size of the PKZILLASs expands our imagination on the capabilities of enzymes in the construction of complex molecules. And finally, the unprecedented abundance of non-elongating KS^0^s are featured in modules associated with the construction of the polycyclic interior of prymnesin, which may contribute to the timing and mechanism of polyether assembly.

No comparable PKS system has yet been identified from a toxic dinoflagellate, however, numerous studies have established that dinoflagellates encode large numbers of modular and single-domain type I PKSs (*41*), including a promising, yet reportedly incomplete 35-knt 7-module PKS transcript candidate from the ciguatoxin-producing *Gambierdiscus polynesiensis* (*12*). If dinoflagellates similarly encode giant PKSs reminiscent of the PKZILLAs, then common transcriptomic practices involving poly-A pulldown RNA sequencing may bias against giant transcripts and instead require length-unbiased rRNA-depletion RNA-seq alongside customized assembly and annotation as performed in this study. Notably, the PKZILLAs went unreported from recent *P. parvum* transcriptomic (*11*) and genomic (*42*) analyses, highlighting the challenges of assembling and annotating giant PKS genes with their highly repetitive sequences.

Dinoflagellate polyketides also share a distinctive biosynthetic feature involving the irregular incorporation of intact and C1-deleted acetate building blocks as illuminated by isotope labeling studies (*43*). The prymnesin biosynthetic model, on the other hand, supports the intact incorporation of 43 contiguous malonate units, which is standard in most bacterial systems. The α-hydroxylating FLX domains and “pass-through” modules found within the PKZILLA modules provide a tantalizing hypothesis for this yet to be described dinoflagellate PKS biochemistry: Assembly line oxidation to the α-ketone followed by transacylation by a KS^0^ may lead to excision of single carbon atoms by decarbonylation as precedented in the biosynthesis of marine polyketides enterocin (*44*) and barbamide (*45*).

Though PKZILLA-1 and -2 are responsible for the construction of the majority of the prymnesin molecule, additional enzymes (acyltransferases, desaturases, hydroxylases, chlorinases, epoxidases, glycosyltransferases) are needed to complete the full biosynthetic pathway and install prymnesin’s remaining functional groups and sugar moieties. In contrast to the only other microalgal toxin with a fully resolved genetic basis, the small alkaloid domoic acid with its clustered causal genes (*46*), the distribution of the PKZILLA-1 and -2 genes across separate pseudo-chromosomes indicates prymnesin biosynthesis is not encoded within a single biosynthetic gene cluster, and suggests that its tailoring enzymes may also not be clustered. The discovery of the PKZILLAs and their role in prymnesin biosynthesis lays the foundation for the development and implementation of alternative linked ‘omics approaches to fully uncover the complete suite of prymnesin biosynthetic enzymes. Moreover, the PKZILLAs now offer the opportunity to dissect the enzymology of ladder-frame polyether biosynthesis and will serve as a model to capture and dissect giant genes, transcripts, and proteins in specialized metabolism.

## Acknowledgments

The authors acknowledge helpful discussions & feedback from members of the Moore laboratory, and from Prof. April Lukowski (UCSD). This publication includes data generated at the UC San Diego IGM Genomics Center utilizing an Illumina NovaSeq 6000 that was purchased with funding from a National Institutes of Health SIG grant (#S10 OD026929).

## Author contributions

Conceptualization: TRF, BSM

Data curation: TRF, VVS, IHW, RPA

Formal Analysis: TRF, VVS, IHW

Funding acquisition TRF, VVS, DJG, JHW, BSM

Investigation: TRF, VVS, IHW, RPA

Methodology: TRF, VVS, IHW, RPA

Project administration: DJG, JHW, BSM

Resources: TRF, BSM

Software: TRF Supervision: TRF, BSM

Validation: TRF, VVS, BSM

Visualization: TRF, VVS, BSM

Writing – original draft: TRF, VVS, BSM

Writing – review & editing: TRF, VVS, IHW, RPA, DJG, JHW, BSM

## Data and materials availability

Raw rRNA depletion RNA-Seq has been deposited to the NCBI SRA archive (BioProject PRJNA936443). The mass spectrometry proteomics data have been deposited to the ProteomeXchange Consortium via the PRIDE (*47*) partner repository with the dataset identifier PXD044632 and doi:10.6019/PXD044632. [[During the review process, the data can be accessed with the following credentials upon login to the PRIDE website (https://www.ebi.ac.uk/pride/archive/login): Username, Password: ERk9Y0kO .]] Other extended datasets and analysis code (scripts) are available on zenodo.org and/or github.com and are both cited in-line throughout the manuscript and listed in table S14.

## Supplementary Materials

Materials and Methods

Figs. S1 to S20

Tables S1 to S14

References (1-94)

## References and Notes

1. G. M. Hallegraeff, D. M. Anderson, K. Davidson, F. Gianella, P. Hansen, Fish-Killing Marine Algal Blooms: Causative Organisms, Ichthyotoxic Mechanisms, Impacts and Mitigation. (UNESCO, Paris, France, 2023; 10.25607/OBP-1964)IOC Manuals and Guides.

2. J. Sobieraj, D. Metelski, Insights into Toxic Prymnesium parvum Blooms as a Cause of the Ecological Disaster on the Odra River. Toxins 15, 403 (2023).

3. D. M. Anderson, E. Fensin, C. J. Gobler, A. E. Hoeglund, K. A. Hubbard, D. M. Kulis, J. H. Landsberg, K. A. Lefebvre, P. Provoost, M. L. Richlen, J. L. Smith, A. R. Solow, V. L. Trainer, Marine harmful algal blooms (HABs) in the United States: History, current status and future trends. Harmful Algae 102, 101975 (2021).

4. K. C. Nicolaou, M. O. Frederick, R. J. Aversa, The Continuing Saga of the Marine Polyether Biotoxins. Angew. Chem. Int. Ed. 47, 7182–7225 (2008).

5. T. Igarashi, M. Satake, T. Yasumoto, Structures and Partial Stereochemical Assignments for Prymnesin-1 and Prymnesin-2: Potent Hemolytic and Ichthyotoxic Glycosides Isolated from the Red Tide Alga Prymnesium parvum. J. Am. Chem. Soc. 121, 8499–8511 (1999).

6. R. E. Moore, G. Bartolini, Structure of palytoxin. J. Am. Chem. Soc. 103, 2491–2494 (1981).

7. M. Murata, H. Naoki, T. Iwashita, S. Matsunaga, M. Sasaki, A. Yokoyama, T. Yasumoto, Structure of maitotoxin. J. Am. Chem. Soc. 115, 2060–2062 (1993).

8. K. Nakanishi, The chemistry of brevetoxins: A review. Toxicon 23, 473–479 (1985).

9. I. Vilotijevic, T. F. Jamison, Epoxide-Opening Cascades Promoted by Water. Science 317, 1189–1192 (2007).

10. A. Nivina, K. P. Yuet, J. Hsu, C. Khosla, Evolution and Diversity of Assembly-Line Polyketide Synthases. Chem. Rev. 119, 12524–12547 (2019).

11. K. Anestis, G. S. Kohli, S. Wohlrab, E. Varga, T. O. Larsen, P. J. Hansen, U. John, Polyketide synthase genes and molecular trade-offs in the ichthyotoxic species Prymnesium parvum. Sci. Total Environ. 795, 148878 (2021).

12. F. M. V. Dolah, J. S. Morey, S. Milne, A. Ung, P. E. Anderson, M. Chinain, Transcriptomic analysis of polyketide synthases in a highly ciguatoxic dinoflagellate, Gambierdiscus polynesiensis and low toxicity Gambierdiscus pacificus, from French Polynesia. PLOS ONE 15, e0231400 (2020).

13. N. Heimerl, E. Hommel, M. Westermann, D. Meichsner, M. Lohr, C. Hertweck, A. R. Grossman, M. Mittag, S. Sasso, A giant type I polyketide synthase participates in zygospore maturation in Chlamydomonas reinhardtii. Plant J. 95, 268–281 (2018).

14. M. H. Medema, T. de Rond, B. S. Moore, Mining genomes to illuminate the specialized chemistry of life. Nat. Rev. Genet. 22, 553–571 (2021).

15. M.-L. Bang, T. Centner, F. Fornoff, A. J. Geach, M. Gotthardt, M. McNabb, C. C. Witt, D. Labeit, C. C. Gregorio, H. Granzier, S. Labeit, The Complete Gene Sequence of Titin, Expression of an Unusual ≈700-kDa Titin Isoform, and Its Interaction With Obscurin Identify a Novel Z-Line to I-Band Linking System. Circ. Res. 89, 1065–1072 (2001).

16. M. Sasaki, N. Takeda, H. Fuwa, R. Watanabe, M. Satake, Y. Oshima, Synthesis of the JK/LM-ring model of prymnesins, potent hemolytic and ichthyotoxic polycyclic ethers isolated from the red tide alga Prymnesium parvum: confirmation of the relative configuration of the K/L-ring juncture. Tetrahedron Lett. 47, 5687–5691 (2006).

17. T. Hashimoto, J. Hashimoto, I. Kozone, K. Amagai, T. Kawahara, S. Takahashi, H. Ikeda, K. Shin-ya, Biosynthesis of Quinolidomicin, the Largest Known Macrolide of Terrestrial Origin: Identification and Heterologous Expression of a Biosynthetic Gene Cluster over 200 kb. Org. Lett. 20, 7996–7999 (2018).

18. S. Donadio, M. Staver, J. McAlpine, S. Swanson, L. Katz, Modular organization of genes required for complex polyketide biosynthesis. Science 252, 675–679 (1991).

19. S. B. Binzer, D. K. Svenssen, N. Daugbjerg, C. Alves-de-Souza, E. Pinto, P. J. Hansen, T. O. Larsen, E. Varga, A-, B- and C-type prymnesins are clade specific compounds and chemotaxonomic markers in Prymnesium parvum. Harmful Algae 81, 10–17 (2019).

20. J. H. Wisecaver, R. P. Auber, A. L. Pendleton, N. F. Watervoort, T. R. Fallon, O. L. Riedling, S. R. Manning, B. S. Moore, W. W. Driscoll, Extreme genome diversity and cryptic speciation in a harmful algal-bloom-forming eukaryote. Curr. Biol. 33, 2246-2259.e8 (2023).

21. J. H. Wisecaver, J. D. Hackett, Dinoflagellate Genome Evolution. Annu. Rev. Microbiol. 65, 369–387 (2011).

22. H.-H. Hong, H.-G. Lee, J. Jo, H. M. Kim, S.-M. Kim, J. Y. Park, C. B. Jeon, H.-S. Kang, M. G. Park, C. Park, K. Y. Kim, H.-H. Hong, H.-G. Lee, J. Jo, H. M. Kim, S.-M. Kim, J. Y. Park, C. B. Jeon, H.-S. Kang, M. G. Park, C. Park, K. Y. Kim, The exceptionally large genome of the harmful red tide dinoflagellate Cochlodinium polykrikoides Margalef (Dinophyceae): determination by flow cytometry. Algae 31, 373–378 (2016).

23. National Center for Biotechnology Information (NCBI), Sequence Read Archive (SRA), Prymnesium parvum strain 12B Illumina reads - SRA PRJNA201451, SRR1685644. https://www.ncbi.nlm.nih.gov/sra/?term=SRR1685644.

24. Y. Dou, Y. Liu, X. Yi, L. K. Olsen, H. Zhu, Q. Gao, H. Zhou, B. Zhang, SEPepQuant enhances the detection of possible isoform regulations in shotgun proteomics. Nat. Commun. 14, 5809 (2023).

25. P. Jones, D. Binns, H.-Y. Chang, M. Fraser, W. Li, C. McAnulla, H. McWilliam, J. Maslen, A. Mitchell, G. Nuka, S. Pesseat, A. F. Quinn, A. Sangrador-Vegas, M. Scheremetjew, S.-Y. Yong, R. Lopez, S. Hunter, InterProScan 5: genome-scale protein function classification. Bioinformatics 30, 1236–1240 (2014).

26. F. Hemmerling, K. E. Lebe, J. Wunderlich, F. Hahn, An Unusual Fatty Acyl:Adenylate Ligase (FAAL)– Acyl Carrier Protein (ACP) Didomain in Ambruticin Biosynthesis. ChemBioChem 19, 1006–1011 (2018).

27. F. Hemmerling, R. A. Meoded, A. E. Fraley, H. A. Minas, C. L. Dieterich, M. Rust, R. Ueoka, K. Jensen, E. J. N. Helfrich, C. Bergande, M. Biedermann, N. Magnus, B. Piechulla, J. Piel, Modular Halogenation, α-Hydroxylation, and Acylation by a Remarkably Versatile Polyketide Synthase. Angew. Chem. Int. Ed. 61, e202116614 (2022).

28. A. J. Winter, R. N. Khanizeman, A. M. C. Barker-Mountford, A. J. Devine, L. Wang, Z. Song, J. A. Davies, P. R. Race, C. Williams, T. J. Simpson, C. L. Willis, M. P. Crump, Structure and Function of the α-Hydroxylation Bimodule of the Mupirocin Polyketide Synthase. Angew. Chem. Int. Ed. 62, e202312514 (2023).

29. E. J. N. Helfrich, J. Piel, Biosynthesis of polyketides by trans-AT polyketide synthases. Nat. Prod. Rep. 33, 231–316 (2016).

30. D. T. Wagner, J. Zeng, C. B. Bailey, D. C. Gay, F. Yuan, H. R. Manion, A. T. Keatinge-Clay, Structural and Functional Trends in Dehydrating Bimodules from trans-Acyltransferase Polyketide Synthases. Structure 25, 1045-1055.e2 (2017).

31. J. Masschelein, P. K. Sydor, C. Hobson, R. Howe, C. Jones, D. M. Roberts, Z. Ling Yap, J. Parkhill, E. Mahenthiralingam, G. L. Challis, A dual transacylation mechanism for polyketide synthase chain release in enacyloxin antibiotic biosynthesis. Nat. Chem. 11, 906–912 (2019).

32. E. J. N. Helfrich, R. Ueoka, A. Dolev, M. Rust, R. A. Meoded, A. Bhushan, G. Califano, R. Costa, M. Gugger, C. Steinbeck, P. Moreno, J. Piel, Automated structure prediction of trans-acyltransferase polyketide synthase products. Nat. Chem. Biol. 15, 813–821 (2019).

33. F. Taft, M. Brünjes, T. Knobloch, H. G. Floss, A. Kirschning, Timing of the Δ10,12-Δ11,13 Double Bond Migration During Ansamitocin Biosynthesis in Actinosynnema pretiosum. J. Am. Chem. Soc. 131, 3812–3813 (2009).

34. G. Hibi, T. Shiraishi, T. Umemura, K. Nemoto, Y. Ogura, M. Nishiyama, T. Kuzuyama, Discovery of type II polyketide synthase-like enzymes for the biosynthesis of cispentacin. Nat. Commun. 14, 8065 (2023).

35. L. Gu, B. Wang, A. Kulkarni, J. J. Gehret, K. R. Lloyd, L. Gerwick, W. H. Gerwick, P. Wipf, K. Håkansson, J. L. Smith, D. H. Sherman, Polyketide Decarboxylative Chain Termination Preceded by O-Sulfonation in Curacin A Biosynthesis. J. Am. Chem. Soc. 131, 16033–16035 (2009).

36. Y. Jiang, A. Kim, C. Olive, J. C. Lewis, Selective C-H Halogenation of Alkenes and Alkynes Using Flavin-Dependent Halogenases. ChemRxiv [Preprint] (2023). 10.26434/chemrxiv-2023-23r4l.

37. T. Hua, D. Wu, W. Ding, J. Wang, N. Shaw, Z.-J. Liu, Studies of Human 2,4-Dienoyl CoA Reductase Shed New Light on Peroxisomal β-Oxidation of Unsaturated Fatty Acids. J. Biol. Chem. 287, 28956–28965 (2012).

38. Yong-Yeng Lin, Martin Risk, Sammy M. Ray, Donna Van Engen, Jon Clardy, Jerzy Golik, John C. James, Koji Nakanishi, Isolation and structure of brevetoxin B from the “red tide” dinoflagellate Ptychodiscus brevis (Gymnodinium breve). J. Am. Chem. Soc. 103, 6773–6775 (1981).

39. Z.-P. Jiang, S.-H. Sun, Y. Yu, A. Mándi, J.-Y. Luo, M.-H. Yang, T. Kurtán, W.-H. Chen, L. Shen, J. Wu, Discovery of benthol A and its challenging stereochemical assignment: opening up a new window for skeletal diversity of super-carbon-chain compounds. Chem. Sci. 12, 10197–10206 (2021).

40. J. Cortes, P. Schöffski, B. A. Littlefield, Multiple modes of action of eribulin mesylate: Emerging data and clinical implications. Cancer Treat. Rev. 70, 190–198 (2018).

41. F. M. V. Dolah, G. S. Kohli, J. S. Morey, S. A. Murray, Both modular and single-domain Type I polyketide synthases are expressed in the brevetoxin-producing dinoflagellate, Karenia brevis (Dinophyceae). J. Phycol. 53, 1325–1339 (2017).

42. J. Jian, Z. Wu, A. Silva-Núñez, X. Li, X. Zheng, B. Luo, Y. Liu, X. Fang, C. T. Workman, T. O. Larsen, P. J. Hansen, E. C. Sonnenschein, Long-read genome sequencing provides novel insights into the harmful algal bloom species Prymnesium parvum. Sci. Total Environ., 168042 (2023).

43. R. M. Van Wagoner, M. Satake, J. L. C. Wright, Polyketide biosynthesis in dinoflagellates: what makes it different? Nat. Prod. Rep., 37 (2014).

44. R. Teufel, A. Miyanaga, Q. Michaudel, F. Stull, G. Louie, J. P. Noel, P. S. Baran, B. Palfey, B. S. Moore, Flavin-mediated dual oxidation controls an enzymatic Favorskii-type rearrangement. Nature 503, 552–556 (2013).

45. S. Guo, Y. Sang, C. Zheng, X.-S. Xue, Z. Tang, W. Liu, Enzymatic α-Ketothioester Decarbonylation Occurs in the Assembly Line of Barbamide for Skeleton Editing. J. Am. Chem. Soc. 145, 5017–5028 (2023).

46. J. K. Brunson, S. M. K. McKinnie, J. R. Chekan, J. P. McCrow, Z. D. Miles, E. M. Bertrand, V. A. Bielinski, H. Luhavaya, M. Oborník, G. J. Smith, D. A. Hutchins, A. E. Allen, B. S. Moore, Biosynthesis of the neurotoxin domoic acid in a bloom-forming diatom. Science 361, 1356–1358 (2018).

47. Y. Perez-Riverol, J. Bai, C. Bandla, D. García-Seisdedos, S. Hewapathirana, S. Kamatchinathan, D. J. Kundu, A. Prakash, A. Frericks-Zipper, M. Eisenacher, M. Walzer, S. Wang, A. Brazma, J. A. Vizcaíno, The PRIDE database resources in 2022: a hub for mass spectrometry-based proteomics evidences. Nucleic Acids Res. 50, D543–D552 (2022).

