## Supplementary Materials for "Giant polyketide synthase enzymes biosynthesize a giant marine polyether biotoxin"

Timothy R. Fallon[1]*, Vikram V. Shende[1], Igor H. Wierzbicki[2], Robert P. Auber[3,4], David J. Gonzalez[2,5], Jennifer H. Wisecaver[3,4], Bradley S. Moore[1,5]*
[1]Center for Marine Biotechnology and Biomedicine, Scripps Institution of Oceanography and University of California, San Diego; 9500 Gilman Dr #0204, La Jolla, CA 92093, USA.
[2]Department of Pharmacology, University of California, San Diego; 9500 Gilman Dr, La Jolla, CA 92093, USA.
[3]Department of Biochemistry, Purdue University; 175 S University St, West Lafayette, IN 47907, USA.
[4]Purdue Center for Plant Biology, Purdue University; 175 S University St, West Lafayette, IN 47907, USA.
[5]Skaggs School of Pharmacy and Pharmaceutical Sciences, University of California, San Diego; 9500 Gilman Dr, La Jolla, CA 92093, USA.

### Materials and Methods

**Cell strains:**

The haploid *Prymnesium parvum* strain 12B1 (*48*) was a gift of Dr. William Driscoll (Pennsylvania State University, Harrisburg). Strain 12B1 is available from the UTEX Culture Collection of Algae at UT-Austin under UTEX accession UTEX LB 3227.

**Media preparation:**

*For proteomic mass spectrometry:*

L1-Si@25 (25% salinity) algal growth media (*49*) was prepared from 12L of 18.2 Mohm·cm water (ultrapure water) dispensed from a laboratory water purifier [Millipore, Milli-Q Advantage A10, CAT# Z00Q0V0WW], 4L seawater [Scripps Institution of Oceanography seawater system], concentrated stocks for L1 media [National Center for Marine Algae, CAT# MKL150L] and 1.5 mM $NaHCO_3$ to adjust for the lack of carbon from seawater, all within a polycarbonate 20L carboy [Thermo Fisher Scientific, CAT# 2251-0050]. The carboy was then capped with a 3-port closure [Thermo Fisher Scientific, CAT# 2162-0831] plus pre-installed interior 6" and 30" silicone tubing. Two of the three ports were temporarily blocked for autoclaving, while the third port was interfaced with a small piece of silicone tubing and a Whatman HEPA-VENT disk filter [Cytiva, CAT# 6723-5000]. The vented carboy was autoclaved using the "Cycle 3 LIQUID 60 MIN" in a Beta Star Medium Sterilizer Series autoclave.

*For rRNA depletion RNA-Seq:*

A similar method was used to above, excepting media preparation was scaled down to 8L in a single 9L bottle [Corning, CAT# 1596-9L] & it was autoclaved in a Hirayama HV-50 autoclave at 121˚C for 60 minutes, with the 0% exhaust setting.

**Creation of v1.1 P. parvum 12B1 reference genome by targeted reassembly:**

Previously published (*20*) Oxford Nanopore Technology (ONT) MinION genomic DNA reads [# of reads: 447907, # of nucleotides: 4005418850, filename:12B1_above_3kb_nanopore_sorted.bam, checksum: seqkit.v0.1_DLS_k0_9c68687a7acb147c412be4b83753e237, SRA:SRR18033808] were aligned against an unpublished intermediate version v0.3 of the 12B1 reference genome assembly [12B1_scaffolds_v0.3.fasta ; checksum: seqkit.v0.1_DLS_k0_6561035ed292bc5039428c769267f70b] using the NGMLR aligner (*50*). A suspicious region [v1.0:12B1-Scaf17:2286312-2287047] of the PKZILLA-1 N-terminus that showed increased coverage, a higher proportion of visible single nucleotide variation (SNV), and a high proportion of a large indel was noted through graphical inspection with Integrated Genomics Viewer v2.14 (*51*). ONT gDNA reads intersecting this suspicious PKZILLA-1 N-terminal region, were extracted using "bedtools intersect" (*52*). The extracted reads [# of reads: 41, # of nucleotides: 475352, filename:, checksum: seqkit.v0.1_DLS_k0_d202591ca9f50940752c50f50395048e], were then assembled using Flye (v2.9.2-b1786), using a parameter scan from 100-10000 for the "--min-overlap" parameter, and inclusion of the "--keep-haplotypes --meta --scaffold" parameters. The source code of Flye had been patched to allow for values of <1000 for the "--min-overlap" parameter. The resulting 29 targeted assemblies all consisted of only a single contig, of which there were only 5 unique sequences across the 29 assemblies. Next, the PKZILLA-1 gene model was partially lifted over to each of the 5 unique targeted reassembly contigs using the flo pipeline (https://github.com/wurmlab/flo) (*53*), which uses the ucsc-kent tools (http://hgdownload.cse.ucsc.edu/admin/exe/), blat (*54*), GNU parallel (*55*), and genometools (*56*) internally. The translated peptide and nucleotide representations of the lifted-over PKZILLA-1 N-terminal 1$^{st}$ exon, were extracted using a custom Nextflow (*57*) workflow (https://github.com/photocyte/PPYR_OGS/blob/master/utility_scripts/extract_gff_features.nf). These peptide and nucleotide representations of the lifted over PKZILLA-1 features were identical across the 5 unique contigs, so a random representative selection was made for downstream analyses. These representative peptide & nucleotide representations of the lifted 1st exon were next aligned against the analogous extractions of the reference 1st exon using kalign2 (*58*), and these multiple sequence alignments were manually inspected with mview (*59*). This analysis revealed a 609 bp tandem repetitive region that was missing from the reference assembly, but also revealed that the targeted re-assembly was otherwise identical to the reference assembly (excepting a single upstream homopolymer indel, that is not unusual with unpolished Nanopore assemblies & was ignored in further downstream analyses). The v1.1 version of the 12B1 assembly [filename: 12B1_scaffolds_v1.1.fasta ; checksum: seqkit.v0.1_DLS_k0_99243f7852f0aa7f150e6169739e61a0] was then constructed by adding the 609 bp region with the 'replace' subcommand of seqkit, targeting the replacement to its homologous loci on scaffold 12B1-Scaf17 of the v1.0 genome, as determined by kalign2. This 12B1 genome assembly v1.1 is available on Zenodo (doi:10.5281/zenodo.10023322).

**_P. parvum_ growth and harvesting for rRNA depletion RNA-Seq:**

*P. parvum* strain 12B1 was grown within 125 mL borosilicate glass Erlenmeyer flasks in 50 mL of L1-Si@25 media. Mouths of the flasks were sealed against contamination using 10 cm x 10 cm squares of air-permeable and sterilizable cellulose-polyester wrapping paper [Cardinal Health, CAT# 4008] that was sealed around the neck of the flask using rubber bands. Before use, the flasks including mouth seal & rubber bands were sterilized and cleaned by autoclaving with 75 mL of ultrapure water. This water was removed before addition of the L1-Si@25 media & seeding of the *P. parvum* culture. Post-autoclave manipulations of the flasks took place in a Type II Biosafety Cabinet. Ongoing culturing of the flasks took place in a 22˚C incubator [Fisher Scientific, CAT# 146E] retrofit with full spectrum LED grow lights [GE Lighting, BR30, CAT# 93101230], corresponding to ~2200 lux from above under a 14:10 light:dark cycle. Routine cell concentration measurements were recorded with a Muse flow cytometer [Luminex, CAT# 0500-3115]. The day phase cells were harvested 3 hours after the light to dark transition and a concentration of ~378e3 cells/mL. The night phase cells were harvested 3 hours after the dark to light transition and a concentration of 559e3 cells/mL. Harvesting occurred by transferring the flask content to a 50 mL polypropylene (PP) centrifuge tube (CFT) and centrifuging with a swinging bucket centrifuge [Beckman, CAT# X-15R], with parameters: 3000×*g*, 10 minutes, 18˚C, accel=MAX, decel=MAX. The supernatant was removed by decanting, and the cell pellet resuspended in the remaining adherent liquid with a P1000 pipette. The cell suspension was transferred to a 2 mL PP microcentrifuge tube (MCT), and 1000 µL plus 400 µL of 4˚C TRIzol Reagent [Invitrogen, CAT# 1559026] were added in quick succession. Post-TRIzol addition, samples were stored at -80˚C for less than a week.

**RNA extraction and rRNA depletion RNA-Seq:**

TRIzol-stored cell samples (see "P. parvum growth and harvesting for rRNA depletion RNA-Seq" methods) were removed from the -80˚C and 280 µL of 24:1 chloroform:isoamyl alcohol [Acros Organics, CAT#327155000] was added. Samples were manually mixed by inversion for ~1 minute, and then centrifuged in a fixed angle centrifuge at 24,000×*g* for 10 minutes. The upper aqueous layer was transferred to a new 2 mL PP MCT and 1 mL of 100% ethanol (denatured) was added. Total RNA was purified from the resulting mixture using chaotropic silica column nucleic acid purification with a commercial kit [Zymo Research, RNA Clean & Concentrator-25, CAT# R1017], following the manufacturer's instructions (*60*). The day and night phase samples yielded 632 ng and 1177 ng of RNA, respectively. These samples were stored at -80˚C for less than a week. The ppMCTs containing purified RNA were then transported to the UCSD Institute for Genomic Medicine (IGM) genomics core facility on dry ice and stored at -80˚C until use. The samples were prepared into deoxyuridine triphosphate (dUTP)-stranded sequencing libraries by the IGM, using a commercially available kit [Illumina, TruSeq Stranded Total RNA with Ribo-Zero Plant, CAT# 20020610], following the manufacturer's instructions. Libraries were sequenced at IGM on a NovaSeq 6000 sequencer [Illumina] using the paired-end mode. Libraries were sequenced over 4 separate NovaSeq runs to increase coverage. Libraries were split across an unknown number of physical flowcell lanes. Libraries were multiplexed within lanes with an unknown type and number of other samples. Raw RNA-Seq reads are available on NCBI SRA with Bioproject: PRJNA936443, with SRA accessions SRR23517916 (night) and SRR23517917 (day).

**RNA-Seq alignment, reference guided transcriptome assembly, and quantification:**
rRNA depletion RNA-Seq reads were trimmed using trimmomatic v0.39 (*61*) with parameters "ILLUMINACLIP:adapters.fa:2:30:10 SLIDINGWINDOW:4:5 LEADING:5 TRAILING:5 MINLEN:25" and aligned to an unpublished intermediate version v0.3 of the 12B1 reference assembly [12B1_scaffolds_v0.3.fasta ; checksum: seqkit.v0.1_DLS_k0_6561035ed292bc5039428c769267f70b] with hisat2 v2.2.1 (*62*) using parameters "--rna-strandness R --fr --no-mixed --no-discordant --max-intronlen 3000". Reads from the day and night phase samples were merged, and the merged reads were then assembled into transcripts using stringtie v2.2.1 with parameters "--rf" (*63*). Poly-A pulldown (SRR1685644)(*23*) and rRNA depletion RNA-Seq reads shown in figures were re-aligned against the 12B1 v1.1 assembly using the same programs (hisat2) and parameters, excepting the changed parameter "--max-intronlen 5000". RNA-Seq was independently quantified against the 12B1 v1.1 gene-model annotation predicted transcripts (doi:10.5281/zenodo.10023330) using the rRNA depletion day phase (SRR23517917) and night phase (SRR23517916) samples with Kallisto v0.50.0 with parameters "--rf-stranded". Kallisto results are available on Zenodo (doi:10.5281/zenodo.10023426).

**Definition of PKS hotspots within the 12B1 genome**
The *P. parvum* 12B1 PKS hotspots were originally defined off the unpublished intermediate versions v0.3 of the 12B1 reference assembly and gene annotation. First, PKS domain polypeptide sequences from *P. parvum* 12B1 were selected by analyzing the TSV format InterProScan (*25*) results produced from the gene-model derived polypeptides as described in (*20*), and, selecting using bedtools and seqkit, those polypeptide regions that received the InterPro annotation IPR016039, IPR036736, IPR036291, IPR042104 or IPR020807, IPR029063, and IPR020843 matches for the ketosynthase (KS), acyl-carrier-protein (ACP), ketoreductase (KR), dehydratase (DH), SAM methyltransferase (MT), and enoyl-reductase (ER) domains, respectively. The polypeptide sequences of these selected domains were then used as a tblastn query against the v0.3 12B1 genomic assembly, using parameters \`-outfmt 5 -evalue 0.001\`. The blast XML results were converted to GFF using a custom script (https://github.com/photocyte/general_scripts/blob/master/blastxml2gff.py). Next, overlapping and redundant tblastn hits on the same strand were collapsed using 'bedtools merge -s'. These bed alignments, representing same-strand coding evidence for PKS domains, were clustered into "hotspots" i.e. regions with evidence of PKS coding sequence without explicit hypotheses as to the gene model structures, via merging adjacent alignments through 'bedtools slop -b' and 'bedtools merge'. The bedtools slop -b parameter was iterated over a range of values, and 5000 bp was determined as the most interpretable where each hotspot likely arose from 1 gene. These v0.3 assembly and annotation derived hotspots were lifted over to the v1.1 assembly without issue using seqkit and bedtools, and are available on Zenodo (doi:10.5281/zenodo.10309063).

**Construction of PKZILLA gene models:**
The construction of the PKZILLA gene models was originally based off an unpublished intermediate version v0.3 of the 12B1 reference assembly. Supervised gene models for the PKZILLAs were constructed by joining or extending predicted exons from the hisat2/stringtie RNA-Seq alignment and reference guided transcriptome assembly (see "RNA-Seq alignment, reference guided transcriptome assembly, and quantification" methods section), with exons from *ab initio* gene models that were generated as previously described (*20*) but based off an unpublished intermediate version v0.3 of the 12B1 reference assembly. For additional guiding evidence, splice-aware protein to genome alignments of PKS protein domains sourced from the *P. parvum* 12B1 gene annotation (*20*) were produced with a

pre-search using tblastn from BLAST+ v2.13.0 (*64*) with the following parameters: "-threshold 13 -word_size 2 -max_intron_length 1000 -outfmt '6 sseqid slen sstart send qseqid qlen qstart qend pident length evalue score' -evalue 0.01", followed by an intron-aware alignment using the prot2genome utility "funannotate-p2g.py", with parameters "--maxintron 500 --exonerate_pident 10" from v1.8.14 of the funannotate genome annotation software (*65*). Funannotate-p2g.py calls exonerate v2.4.0 (*66*) internally. Evidence for potential introns was created using the "hints.BAM.gff" file generated by funannotate v1.8.14 from the hisat2 aligned reads (see "RNA-Seq alignment, reference guided transcriptome assembly, and quantification" methods). Notably, we found that the coding regions of the presumed PKZILLAs were flagged as repeats by the repeatmasking process used to generate the *ab initio* gene models, thus our decision to perform a supervised gene-model construction based on protein-alignments and RNA-seq alignments. This combined evidence was viewed as tracks against the genomic assembly in Integrated Genomics Viewer (IGV) v2.14 (*51*). The PKZILLA gene models (in GFF format) were then iteratively edited with a text editor, with iterative inspection in IGV confirming compatibility with the evidence. As visibility of features in IGV was sensitive to their exact ordering in the GFF file, GFF files were validated for correctness and double sorted with genometools and igvtools before viewing, using a custom Nextflow script (https://github.com/photocyte/PPYR_OGS/blob/master/utility_scripts/doubleSort.nf). After the tracks were validated, features sequences (i.e. gene, mRNA, CDS, polypeptides) were extracted using a custom Nextflow script (https://github.com/photocyte/PPYR_OGS/blob/master/utility_scripts/extract_gff_features.nf).

**Integration of PKZILLA gene models into the reference 12B1 v1.1 gene annotation:**

As the PKZILLA gene models were originally based off an unpublished intermediate version v0.3 of the 12B1 reference assembly (see "Construction of PKZILLA gene models" methods), it was necessary to integrate the PKZILLA gene models into a reference gene model annotation based off the v1.1 genome assembly (see "Creation of v1.1 P. parvum 12B1 reference genome by targeted reassembly" methods). Only scaffolds 12B1-Scaf8 and 12B1-Scaf32 changed between genome assembly version v0.3 used for PKZILLA model creation (see "Construction of PKZILLA gene models" methods section) and the published genome assembly version v1.0 [12B1_scaffolds_v1.fasta ; seqkit.v0.1_DLS_k0_25524daa93426ca36fa6f688345c06d0] (*20*). As described in the "Creation of v1.1 P. parvum 12B1 reference genome by targeted reassembly" methods section, scaffold 12B1-Scaf17 v1.0 was modified to produce genome assembly v1.1 [12B1_scaffolds_v1.1.fasta ; seqkit.v0.1_DLS_k0_99243f7852f0aa7f150e6169739e61a0]. non-PKZILLA-1 v1.0 gene models on the modified 12B1-Scaf17 were lifted over using the flo pipeline (https://github.com/wurmlab/flo) (*53*), which uses the ucsc-kent tools (http://hgdownload.cse.ucsc.edu/admin/exe/), blat (*54*), GNU parallel (*55*), and genometools (*56*) internally. Comparisons of extracted feature sequences (i.e. gene, mRNA, CDS, polypeptides), produced using a custom Nextflow (*57*) workflow (https://github.com/photocyte/PPYR_OGS/blob/master/utility_scripts/extract_gff_features.nf), from the v1.0 and v1.1 gene models confirmed they were successfully lifted over without fragmentation. Next, PKZILLA-1 was aligned and lifted over from v0.3 to the v1.1 assembly using the flo pipeline as described above. Given the lack of changes on the PKZILLA-2 and -3 containing scaffolds, 12B1-Scaf7 and 12B1-Scaf10 respectively, between genome assembly versions v0.3 and v1.1, PKZILLA-2 and -3 were directly lifted over to the v1.1 assembly via concatenation of the GFF files. Fragmented gene models from the v1.0 annotation which overlapped with the lifted over PKZILLA gene models, were removed using bedtools subtract. The identifiers for the removed gene models are available as 'all.remove.list.sorted.txt' on Zenodo (doi:10.5281/zenodo.10023330). In total, 25 gene models were removed. Successful liftover of the PKZILLA gene models was validated by graphical inspection using Integrated Genomics Viewer v2.16 (*51*) and by comparison of seqkit sum hashes of the extracted sequence features. The resulting GFF file of the v1.1 gene models is available as '12B1_v1.1.gff3' on Zenodo (doi:10.5281/zenodo.10023330). The extracted sequence features (gene, mRNA, translated polypeptide, individual translated CDSs, individual exons, individual introns, etc.), were extracted using a custom Nextflow script (https://github.com/photocyte/PPYR_OGS/blob/master/utility_scripts/extract_gff_features.nf) that used genometools (*56*) internally. These extracted v1.1 assembly & v1.1 annotation sequence features are available on Zenodo (doi:10.5281/zenodo.10023330).

***P. parvum*** **growth, harvesting, and lyophilization for proteomic mass spectrometry:**
*Prymnesium parvum* strain 12B1 were grown in a semi-continuous batch format, i.e. removing a subset of culture at late-exponential to early-stationary phase and diluting the remaining culture with fresh media. *P. parvum* were grown on the laboratory benchtop in a 20L polycarbonate carboy [Thermo Fisher Scientific, CAT# 2251-0050] with media prepared as described in the "Media preparation for proteomic mass spectrometry" methods and under a 14:10 light:dark cycle with ~3200 lux from above (measured at shoulder of carboy) from full spectrum LED grow lights [SpiderFarmer, CAT# SF1000], ambient laboratory temperature control (~18-20˚C), and 6 liters per minute (LPM) of aeration with laboratory pressurized air. Air was delivered below the water surface via the interior 6" silicone tubing attached to a 10 mL glass pipette [Fisher Scientific, CAT# 13-678-27F]. The interior 30" silicone tubing remained coiled below the water surface and was used for daily culture sampling (1 mL) using a pipette controller [Corning, CAT# 4099], large scale culture harvesting (1L+) using a peristaltic pump, and addition of fresh autoclaved media using a peristaltic pump. Daily cell concentration measurements were recorded with a Muse flow cytometer [Luminex, CAT# 0500-3115]. For proteomics, cells were harvested at a concentration of 400,000 cells / mL via two rounds of 500 mL centrifugation with parameters: 3500×g, 10 minutes, accel=5, decel=5, temperature=18˚C, in 500 mL polypropylene centrifuge tubes (CFTs) [Corning, CAT# 431123] using a X-14R swinging bucket centrifuge [Beckman Coulter, CAT# A99465]. The centrifugation also used a blue adapter sleeve [Beckman Coulter, CAT# 349846], a gray conical adaptor base [Beckman Coulter], and a compatible rotor and buckets [Beckman Coulter, CAT# SX4750A]. Cell concentration measurements of the supernatant indicated a 54% harvesting efficiency. The pelleted cell mass was flash frozen with liquid $N_2$ inside the 500 mL CFTs and lyophilized for >48 hours using a nominally -103˚C, 12 mTorr lyophilizer [SP Industries, CAT# 4KBTZL-105], backed by a rotary vane vacuum pump [Edwards, CAT# RV5]. Four 500 mL CFTs were lyophilized at a time in a single Multi-Tainer container [FTS Systems]. Next, the 500 mL CFTs were placed on dry ice, and the lyophilized cell mass was scraped using a spatula/scoopula into a dry-ice cooled 50 mL polypropylene CFT [Corning, CAT# 352070], and stored long-term at -80˚C.

**Sample preparation for proteomic mass spectrometry:**
*Preparation of proteomic subsample:*
A 50 mL CFT containing lyophilized *P. parvum* 12B1 biomass (see "*P. parvum* growth, harvesting and lyophilization for proteomic mass spectrometry" method) was removed from the -80˚C and allowed to equilibrate to room temperature under lyophilizer vacuum. A 4.8 mg sample of lyophilized biomass was removed using a spatula, transferred into a 1.5 mL polypropylene microcentrifuge tube (ppMCT), & had its mass measured using an analytical balance [Sartorius, CAT# CPA124S]. The ppMCT containing sample was then transported to the UCSD Collaborative Center for Multiplexed Proteomics (CCMP) proteomics core facility on dry ice and stored at -80˚C until use.

*Cell lysis, reduction, and alkylation:*
The sample was next suspended in 500 µL of Lysis buffer (6M urea, 7% SDS, 50 mM tetraethylammonium bicarbonate (TEAB) [Sigma-Aldrich, CAT# T7408-500ML], pH 7.7 adjusted with phosphoric acid) supplemented with 4-(2-aminoethyl)benzenesulfonyl fluoride (AEBSF) and phenylmethylsulfonyl fluoride (PMSF) containing cOmplete ULTRA Mini Tablets, EDTA-free (½ tablet for 10 mL of buffer ) [Roche, CAT# 05892791001] protease inhibitors, and PhosSTOP (½ tablet for 10 mL of buffer) [Roche, CAT# 4906845001] phosphatase inhibitors. Greater than typical quantities of

biomass were used as input into the method to maximize sensitivity, thus non-solubilized material was still present in the tube after addition of lysis buffer. Next, sedimented non-solubilized material was removed via pipette aspiration (~50 µL) and the remaining cell suspension transferred to a 2 mL Protein LoBind Tube [Eppendorf, CAT# 022431102] ppMCT. The sample was mixed for 10 minutes in a Vortex Genie 2 vortex mixer [Scientific Industries, CAT# 00-SI-0236] and sonicated using Qsonica Sonicator Q125 [Qsonica, CAT# Q125-110] equipped with a 1.6 mm microtip probe [Qsonica, CAT# 4417] with the following parameters: amplitude 20; 1 second pulse; 5 second break; 5 pulses. The lysate was centrifuged for 5 minutes at 16,000×*g*, room temperature (RT), and 250 µL of the supernatant was transferred into a new 2 mL Protein LoBind Tube. The sample was then chemically reduced by addition of 5 µL 0.5 M dithiothreitol (DTT) (9.8 mM final concentration) [Invitrogen, CAT# 15508-013] (prepared in a HPLC-grade water [Fisher Chemical, CAT# W5-4]), and incubated for 30 minutes at 47˚C. Next, the sample was cooled on ice for 5 minutes and the solubilized proteins were alkylated by addition of 15 µL 0.5 M iodoacetamide (IAA) (27.8 mM final concentration) [Sigma-Aldrich, CAT# I1149] (prepared in a HPLC-grade water) with incubation for 45 minutes at RT in the dark. The alkylation reaction was quenched by addition of 5 µL 0.5 M DTT (9.09 mM final concentration) and incubated for 15 minutes at RT. The sample was centrifuged for 5 minutes at 16,000×*g*, RT, and the resulting supernatant containing solubilized chemically reduced and alkylated proteins was transferred into a new 2 mL Protein LoBind Tube.

*Protein digestion:*

The protein sample (~275 µL) was next acidified with 27 µL of 12% phosphoric acid (final concentration: ~100 mM) [Sigma-Aldrich, CAT# 49685-500ML] and mixed with 1500 µL of Binding buffer (90% methanol [Fisher Chemicals, CAT# A452-4], 50 mM TEAB, pH 7.1 adjusted with phosphoric acid). The acidified sample was loaded onto an SDS binding S-Trap mini column [ProtiFi, CAT# C02–mini-80] and centrifuged for 1 minute at 1000×*g*, RT. The column was washed 5 times with 500 µL of Binding buffer with 1 minute centrifugation at 1000×*g*, RT, per each wash. The column was centrifuged for an additional 2 minutes at 2000×*g*, RT, to remove residual methanol. Next, the column was transferred into a new ppMCT collection tube (provided with the S-Trap mini columns) and prepared for on-column digestion by addition of 10 µg (20 µL) of Sequencing Grade Modified Trypsin [Promega, CAT# V5113] and 105 µL of 50 mM TEAB. The column was briefly centrifuged (<2000×g), and the resulting flow-through reapplied onto the column. Next, the proteins were on-column trypsin digested for 3 hours at 47˚C. Peptides were eluted from the column with 125 µL of 50 mM TEAB (1 minute centrifugation at 2000×g, RT), 125 µL of 5% formic acid [Fisher Chemicals, CAT# A118P-500] (1 minute centrifugation at 2000×g, RT), and 125 µL of 50% acetonitrile [Fisher Chemicals, CAT# A998-4] (5 minute centrifugation at 2000×g, RT), resulting in a ~500 µL solution of peptides in ~25 mM TEAB, ~1.25% formic acid, ~12.5% acetonitrile.

*Peptide desalting:*

The peptide solution was frozen at -80˚C and lyophilized using a Savant SPD111V SpeedVac Concentrator [Thermo Scientific, CAT# SPD111V-115], in line with -105˚C Savant RVT5105 Refrigerated Vapor Trap [Thermo Scientific, CAT# RVT5105-115], with vacuum supplied by a Fisher Scientific Maxima D4A Rotary Vane Pump [Thermo Scientific, CAT# 010574A]. The resulting solids were resuspended in 500 µL of 0.1% trifluoroacetic acid (TFA) [Thermo Scientific, CAT# 28901] in HPLC-grade $H_2O$ via vortexing for 20 minutes in a Vortex Genie 2 vortex mixer. A Sep-Pak tC18 1 cc Vac Cartridge (50 mg sorbent) [Waters, CAT# WAT054960] was placed in a NucleoVac 24 Vacuum Manifold [MACHEREY-NAGEL, CAT# 740299] and washed with 1 mL of 100% acetonitrile and 2 mL of 0.1% TFA using laboratory vacuum. The peptide solution was centrifuged for 5 minutes at 16000×*g*, RT, and the supernatant was applied onto the tC18 cartridge. Column-bound peptides were washed with 5 mL of 0.1% TFA and eluted with 750 µL of 40% acetonitrile, 0.5% acetic acid and 750 µL of 80% acetonitrile, 0.5% acetic acid into a 2 mL Protein LoBind Tube.

*Peptide quantification:*

The resulting desalted peptide sample was frozen at -80˚C, lyophilized (as described above), and the solids resuspended in 500 µL of 50% acetonitrile via vortexing in a Vortex Genie 2 vortex mixer for 20 minutes. Notably, an undissolved precipitate was observed at this step, which is unusual for this protocol. The undissolved precipitate may indicate a high proportion of post-translationally modified peptides with poor solubility in 50% acetonitrile, i.e. glycosylated peptides, or other non-peptide molecules that were carried over in the protocol. The peptide solution was next centrifuged for 5 minutes at 16000×*g*, RT, and the supernatant containing only soluble peptides was transferred into a new 2 mL Protein LoBind Tube. The peptide concentration was determined with the BCA-like Biuret-reaction-based Pierce Quantitative Colorimetric Peptide Assay kit [Thermo Scientific, CAT# 23275].

*Sample preparation for mass-spectrometry analysis:*

A 20 µg sample of peptides was frozen at -80˚C and lyophilized (as described above), and resuspended in 20 µL of 5% acetonitrile, 5% formic acid, (final concentration 1 µg/µL) and vortexed at room temperature for 20 minutes in a vortex mixer. Samples were transferred to 300 µL Target Polyspring glass Inserts [Thermo Scientific, CAT# C4010-630] within 9 mm glass autosampler vials with pierceable PTFE septa [Thermo Scientific, CAT# C5000-580W].

**Proteomic liquid chromatography, mass spectrometry:**

1 µL of sample (1 µg of peptides) was injected onto an Easy-Spray PepMap Neo 2 µm C18 75 µm X 150 mm column [Thermo Fisher, CAT# ES75150PN] using an Thermo Easy-nLC 1000 liquid chromatography instrument (HPLC) [Thermo Fisher, CAT# LC120]. Peptides were separated via reverse-phase chromatography at a flow rate of 0.4 µL/minute. Solvent A of the mobile phase consisted of 0.1% formic acid in HPLC-grade water, while solvent B consisted of 0.1% formic acid in acetonitrile. An initial 2-min isocratic gradient flow of 3% B was followed by a linear increase up to 25% B for 85 min, increased to 45% B over 15 min, and a final increase to 95% B over 15 min, whereupon B was held for 6 min and returned to baseline (2 min) and held for 10 min, for a total of 183 min. Sample spectra were collected on an Orbitrap Fusion tribrid mass spectrometer (Thermo Scientific, Wikidata:Q120754733) that collected $MS^1$ data in positive ion mode within the 375 to 1,500 m/z range, Orbitrap resolution: 120000, data type: centroid**,** precursor ion isolation window (m/z): 0.7**,** and HCD collision energy set at 30%. Data dependent $MS^2$ fragmentation spectra were recorded in the linear ion trap. A vendor-software exported report on the used $MS^1$ and $MS^2$ methods is available on Zenodo (doi:10.5281/zenodo.10023360). 3 separate injections & MS runs, i.e. “technical replicates” were performed off the same resuspended peptide sample.

**Proteomic peptide & protein identification:**

Peak lists obtained from MS/MS spectra were identified using X!Tandem version X! Tandem Vengeance (2015.12.15.2) (*67*) and Comet version 2023.01 rev. 2 (*68*). The search was conducted using SearchGUI version 4.2.17 (*69*).

Protein identification was conducted against a concatenated target/decoy (*70*) version of the *Prymnesium parvum* 12B1 polypeptides (predicted from gene models) including 24938 (target) or 49876 (target+decoy) sequences. The filename of the used database was: simple_nostops_12B1v1.1.fasta_concatenated_target_decoy.fasta , the seqkit (v2.5.0) (*71*) checksum was: seqkit.v0.1_PLS_k0_1221bf0c34f4b151e5f4d2c96f391bcd , the MD5 checksum was: 688eaf7347ce157ff94322be6c3d29ff . C-terminal stop codons were removed from the predicted polypeptides before decoy creation. The decoy sequences were created by reversing the target sequences in SearchGUI.

The identification settings were as follows: Trypsin, Specific, with a maximum of 2 missed cleavages 15.0 ppm as $MS^1$ and 0.2 Da (Th) as $MS^2$ tolerances; fixed modifications:

Carbamidomethylation of C (+57.021464 Da), variable modifications: Oxidation of M (+15.994915 Da), fixed modifications during refinement procedure: Carbamidomethylation of C (+57.021464 Da), variable modifications during refinement procedure: Acetylation of protein N-term (+42.010565 Da), Pyrolidone from E (-18.010565 Da), Pyrolidone from Q (-17.026549 Da), Pyrolidone from carbamidomethylated C (-17.026549 Da). All algorithm specific settings are listed in the Protein Identification Certificate of Analysis available as 'certificate_of_analysis.txt' on Zenodo (doi:10.5281/zenodo.10023441).

Peptides and proteins were inferred from the spectrum identification results using PeptideShaker version 2.2.25 (*72*). Peptide Spectrum Matches (PSMs), peptides and proteins were validated at a 1.0% False Discovery Rate (FDR) estimated using the decoy hit distribution. All validation thresholds are listed in the Certificate of Analysis available as 'certificate_of_analysis.txt' on Zenodo (doi:10.5281/zenodo.10023441). Spectrum counting abundance indexes were estimated using the Normalized Spectrum Abundance Factor (*73*) adapted for better handling of protein inference issues and peptide detectability.

Source data files (.raw, .mzML files) are available via ProteomeXchange with identifier PXD044632 (doi:10.6019/PXD044632), while resulting files from the downstream SearchGUI & PeptideShaker proteomic analysis are available on Zenodo (doi:10.5281/zenodo.10023441).

**Analysis of proteomic results:**

The 'Default Peptide Report.txt' TSV from the SearchGUI & PeptideShaker analysis was filtered down to those peptides that confidently matched (as scored by SearchGUI & PeptideShaker) against the PKZILLAs using a custom Jupyter notebook. The vast majority (68/70) of confident peptides that matched against the PKZILLAs, cross-matched only to the PKZILLAs, and the remaining 2/70 that also matched non-PKZILLA proteins, were removed from downstream analyses, as their removal minorly decreased the confidence of presented interpretations but also simplified explanations of this analysis. The Jupyter notebook next classified confident peptides as protein-unique (only matched a single PKZILLA protein), or those that were protein-multimatch (matched >= 2 PKZILLA proteins), and further sub-classified those categories down to those peptides that matched/were translated from a single PKZILLA CDS/exon region (exon-unique), or those that arose from multiple PKZILLA CDS/exon regions (exon-multimatch). This Jupyter notebook and spreadsheets of the resulting peptide categorizations are available on Zenodo (doi:10.5281/zenodo.10028959). See also fig. S8, S9.

**IntroProScan annotation of PKZILLA domains:**

The 3 PKZILLA polypeptides were extracted from all gene model polypeptides of the 12B1 gene annotation, and annotated with InterProScan v5.64-96.0 (*25*) using Singularity/Apptainer v3.5.8 (*74*, *75*) execution of the InterProScan Docker image, via a custom Nextflow wrapper workflow (*76*). In brief, all available InterProScan searches were performed, and post-processing off the GFF annotations was applied to remove certain PKS domain annotations for the KR and ER domains (Table S3) that gave erroneous duplicate annotations from shared homology. The workflow and InterProScan results for the PKZILLA polypeptides are available on Zenodo (doi:10.5281/zenodo.10023460). Specific InterProScan annotations that were deemed to be diagnostic for and representative of particular PKS domains are described in the "run2_filter.tsv" file of the aforementioned Zenodo dataset, and renaming from the InterPro entry identifiers to human readable identifiers are defined in "renaming.tsv" and "renaming2.tsv". Full graphical plots of all domains on the PKZILLAs are available in the "./results/pdf_2_PDF_A_1B/output/*.pdf" files, whereas graphical plots of a length normalized representation of the diagnostic PKS domains are available in the "./results/beads_on_string:pdf_2_PDF_A_1B/output/*.pdf" files. See also (maintext Fig. 2). Non length normalized representations are shown in fig. S6, S7, and the source analysis is available on Zenodo (doi:10.5281/zenodo.10093617). Graphical plots were generated with DNA Features Viewer (*77*). Non-

PKZILLA polypeptides were independently annotated with the same process but with a separate execution and a simplified workflow entry-point of ‘do_simple_nonparallel_scan’ that did not include graphical plotting. The workflow and InterProScan results for the non-PKZILLA polypeptides are available on Zenodo (doi:10.5281/zenodo.10011739).

**Analysis of active site residues within PKZILLA PKS domains:**
The presence or absence of expected conserved active site residues (catalytic or prosthetic group attachment sites) within domains of the PKZILLAs, were determined for the polyketide ketosynthase (KS), acyl carrier protein (ACP), dehydratase (DH), and ketoreductase (KR), flavin oxygenase (FLX), and enoyl (ER) domains, using either multiple sequence alignment liftover of the active site location & interpretation using the single- or multi-residue motif as defined by InterProScan annotations, or, locations & motifs as defined from literature sources, namely Fig. S4 of (*17*), Fig. S35 of (*78*), and Fig. S5 of (*79*). Notably active site residues for the ER and FLX domains were not available via InterProScan or literature sources. Essential and universally conserved active site residues for the ERs have not been reported despite determined efforts as of 2015 (*80*). For the FLX domains, which via AlphaFold2 (*81*, *82*) structural predictions and FoldSeek (*83*) structural similarity searches (doi: 10.5281/zenodo.10120640) have significant yet divergent structural homology with flavin-containing monooxygenase (FMO) proteins, catalytic active site residues have been claimed for certain structural homologs (*84*), but are not conserved across the full set of active FMO structural homologs (*85*), and thus the FLXs were not included in the PKZILLA active site analysis.

Annotated PKS domains were sourced from the ‘gt_extractfeat’ process results of the PKZILLA InterProScan available on Zenodo (doi:10.5281/zenodo.10023460), and renamed according to their increasing order N-terminal to C-terminal on the PKZILLA polypeptides, i.e. KS1, KS2, KS3, etc. A custom bash script using seqkit (*71*), bedtools (*52*), and kalign2 (*58*), was used to create multiple sequence alignments of,  and extract the residues at matching active sites, from the PKZILLA PKS domains. All ACP, and DH domains for PKZILLA-1 and -2 had the expected active site residues, while the ER and FLX domains were not included due to a lack of active site information. KS domains were scored as $KS^0$ aka “non-elongating”/“condensation-incompetent”/“transacylase” KSs based on the deviation of the histidine in the HGTGT active site motif (*29*, *86*, *87*). In contrast, KR domains had two unusual aspects: **(1)** They came in two classes - contiguous or split. The KR domain is made up of two homologous N-terminal structural and C-terminal catalytic subdomains. Typically, these two subdomains are contiguous or nearly so. However certain fatty acid synthase (FAS) or PKS proteins show a split configuration, where the two KR subdomains are intervened and split by another PKS domain. This configuration can be seen with the InterPro domain annotations for the Porcine (*Sus scrofa*) FAS protein (https://www.ebi.ac.uk/interpro/protein/UniProt/I3LCW1/), where the ER domain intervenes between the two KR subdomains. In the literature, the subdomains showing this phenomena have been dubbed as the N-terminal $KR^s$ or $KR_S$ (structural) or C-terminal $KR^c$ or $KR_C$ (catalytic) subdomains (*88–90*). We use the nomenclature KR or KRf (full-length), KRn (N-terminal), and KRc (C-terminal) to distinguish our 3 subclasses of ketoreductase (sub)domains. Notably, some unannotated gaps in the PKZILLA are similar to the KRn subdomains and show a nearby annotated KRc subdomain, suggesting that KRn domains may have diverged away from bioinformatic detection (table S6, S7). **(2)** The catalytic residues of several KRf and KRc domains do not match with expectations for activity, and thus we interpret their source domains as catalytically inactive. Some KR domains showed expected catalytic residues for activity, yet we anticipated they would be inactive based on our biosynthetic model - thus a 2nd tier of analysis was applied where KR domains predicted to phylogenetically cluster with and descend from the last common ancestor for PKZILLA-1__KR31c0 (inactivity assigned based on active side residues) and PKZILLA-1__KR14c (fig. S12, Table S6, S7, S8, S9) were assigned as inactive. This clade was selected based on their relatively long phylogenetic branches (suggesting relaxation of selection from inactivity), and the matching with expectation of inactivity from the biosynthetic model. An alternative hypothesis is they are catalytically active domains but catalyze non-standard reactions. Throughout the manuscript and figures we dub these putatively inactive or diverged domains as $KR^0$/$KRf^0$/KRf0, $KRn^0$/KRn0, or $KRc^0$/KRc0. The workflow and results for this analysis, including Excel spreadsheets of the sequences at active site residues, is available on Zenodo (doi:10.5281/zenodo.10028517).

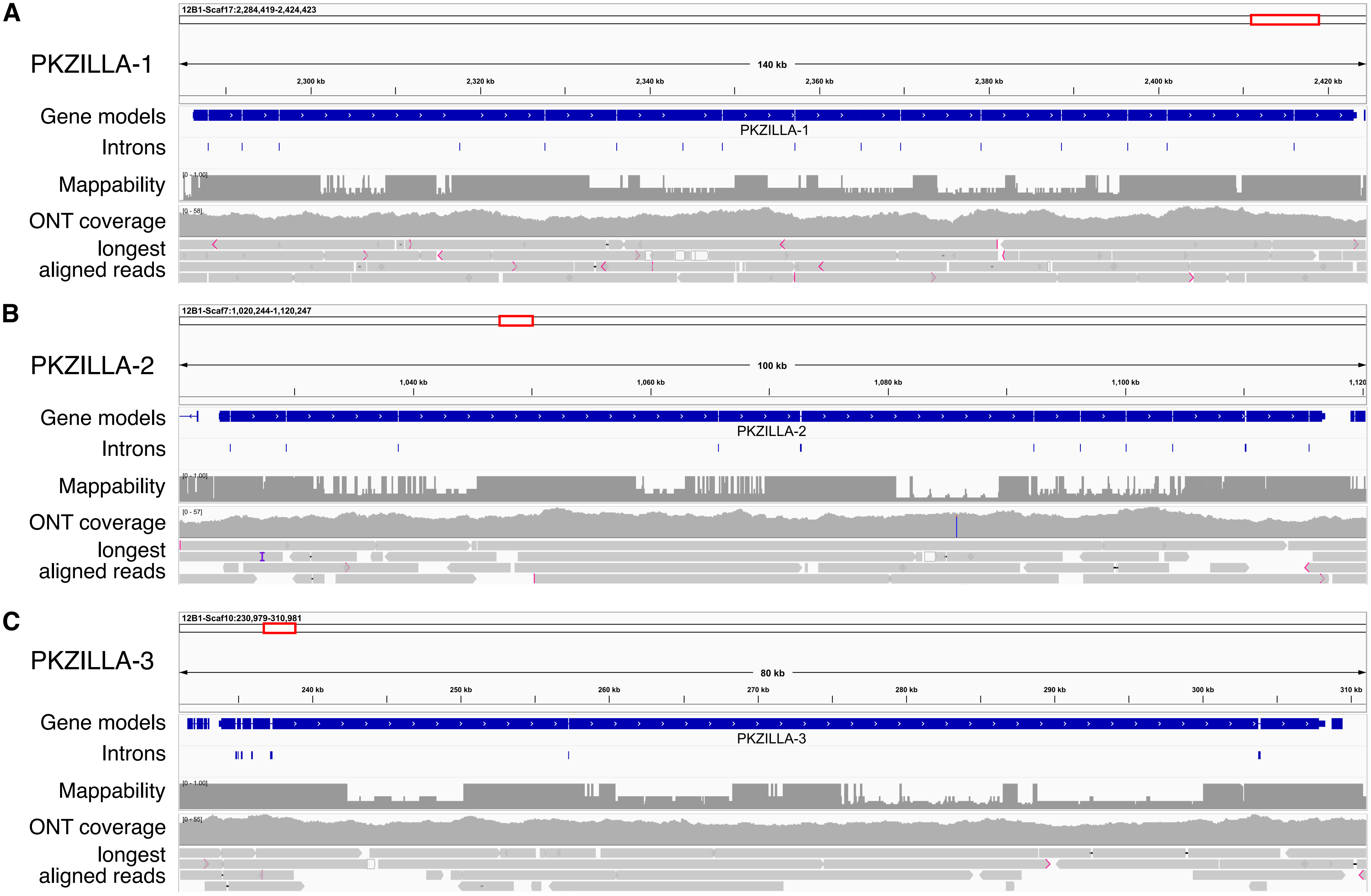
A
PKZILLA-1
12B1-Scaf17:2,284,419-2,424,423
140 kb
2,300 kb
2,320 kb
2,340 kb
2,360 kb
2,380 kb
2,400 kb
2,420 kb
Gene models
PKZILLA-1
Introns
Mappability
[0 - 1.00]
ONT coverage
[0 - 58]
longest aligned reads
B
PKZILLA-2
12B1-Scaf7:1,020,244-1,120,247
100 kb
1,040 kb
1,060 kb
1,080 kb
1,100 kb
1,120
Gene models
PKZILLA-2
Introns
Mappability
[0 - 1.00]
ONT coverage
[0 - 57]
longest aligned reads
C
PKZILLA-3
12B1-Scaf10:230,979-310,981
80 kb
240 kb
250 kb
260 kb
270 kb
280 kb
290 kb
300 kb
310 kb
Gene models
PKZILLA-3
Introns
Mappability
[0 - 1.00]
ONT coverage
[0 - 55]
longest aligned reads

**Fig. S1:** Expanded genomic detail for PKZILLA gene models. PKZILLA-1,-2,-3 are shown as panel A,B,C. Aligned Oxford Nanopore (ONT) reads (*20*) show no evidence of assembly errors like sharp coverage discontinuities, frequent coincident insertions or deletions, or frequent coincident single nucleotide variations (SNVs), excluding the synonymous SNV T→C (L20550L) in PKZILLA-2. Graphical key: Pink arrows=read soft-clips. Vertical purple bar=read insertions. Black horizontal line=read deletions. IGV parameters: Allele Frequency Threshold=0.70, Hide Small Indels=Yes, Sort Alignments by=aligned read length. Show mismatched bases=Off. Mappability was calculated with genmap v1.3.0 (*91*), with parameters K=100 and E=2. Drops from maximum of the mappability track represent low short read mappability due to repetitive sequences. Figure produced with Integrated Genomics Viewer (IGV) v2.16.2 (*51*).

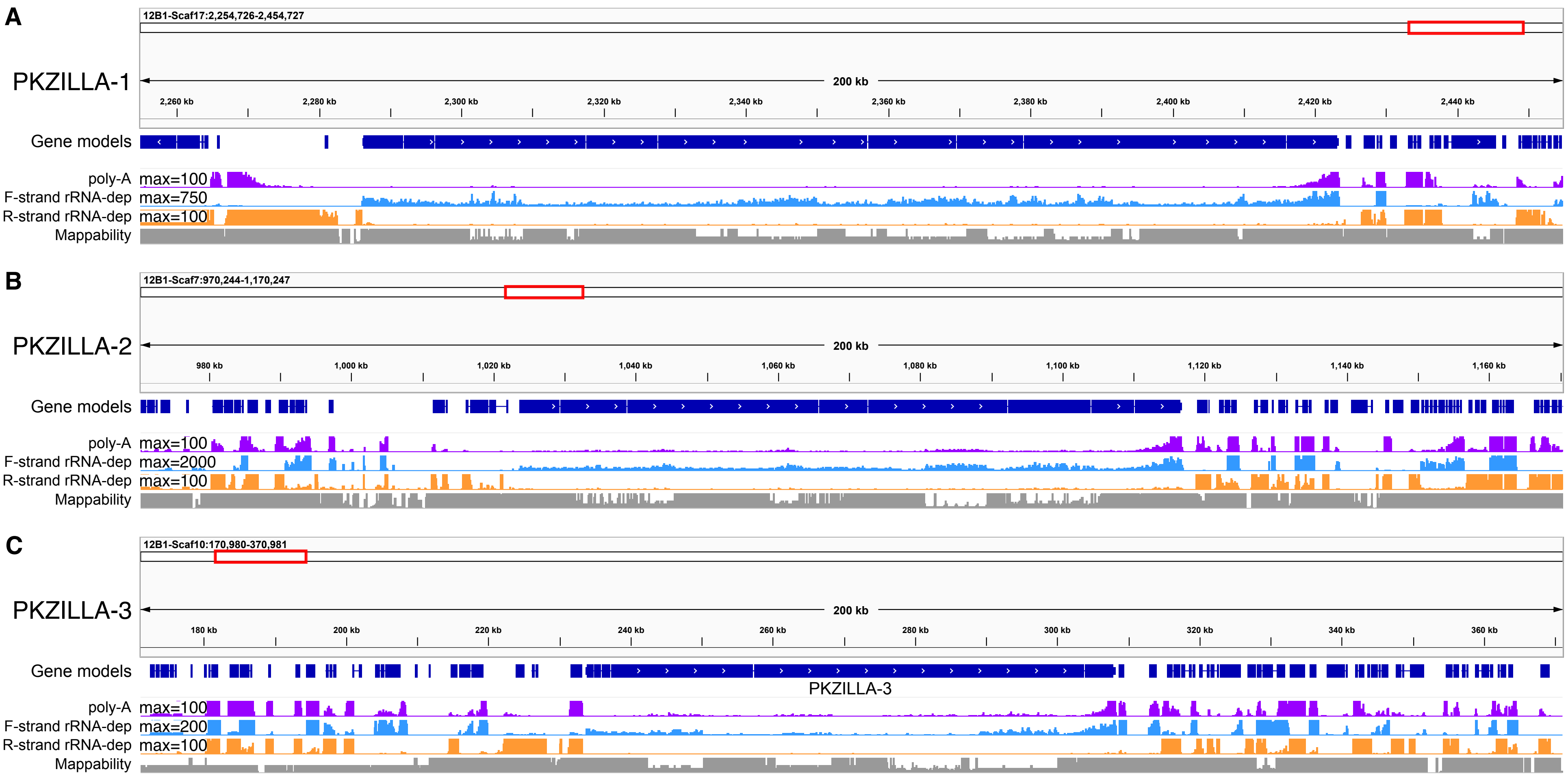


**Fig. S2**: Expanded transcriptomic detail for PKZILLA gene models. PKZILLA-1,-2,-3 are shown as panel A,B,C. poly-A=Coverage of hisat2 aligned poly-A pulldown RNA-Seq reads. F-strand rRNA-dep=Coverage of extracted forward-stranded reads for hisat2 aligned rRNA depletion RNA-Seq reads. R-strand rRNA-dep=Coverage of extracted reverse-stranded reads for hisat2 aligned rRNA depletion RNA-Seq reads. RNA-Seq coverage tracks have a linear scale, scaled to the maximum values annotated on the tracks. Note that the poly-A and R-strand tracks are scaled to a lower maximum to enhance sensitivity. Note that the antisense strand has negligible coverage, supporting one transcriptional start site (TSS) per hotspot. Mappability was calculated with genmap v1.3.0 (*91*), with parameters K=100 and E=2. Drops from maximum of the mappability track represent low short read mappability due to repetitive sequences. Figure produced with Integrated Genomics Viewer (IGV) v2.16.2 (*51*).

#### A - PKZILLA-1 Intron #1

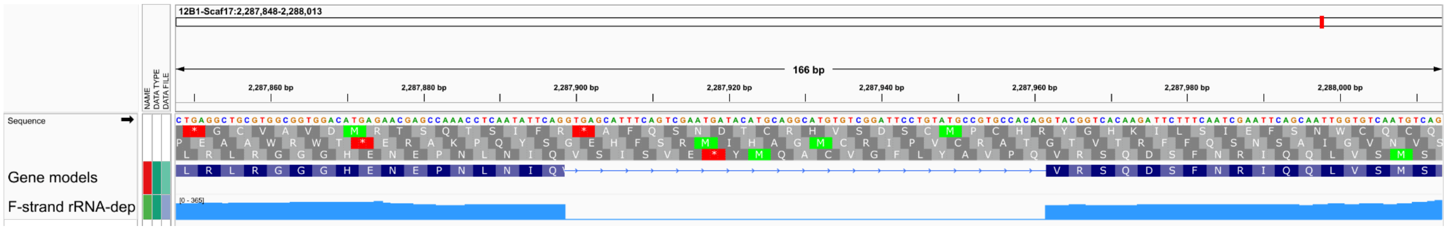


#### B - PKZILLA-1 Intron #2

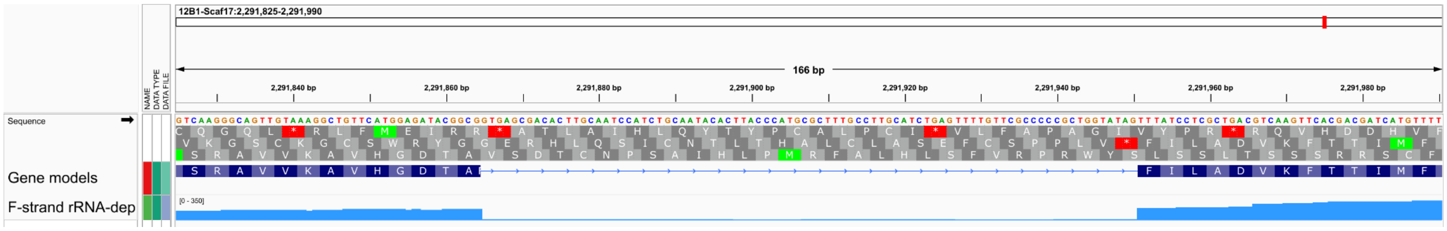


#### C - PKZILLA-1 Intron #3

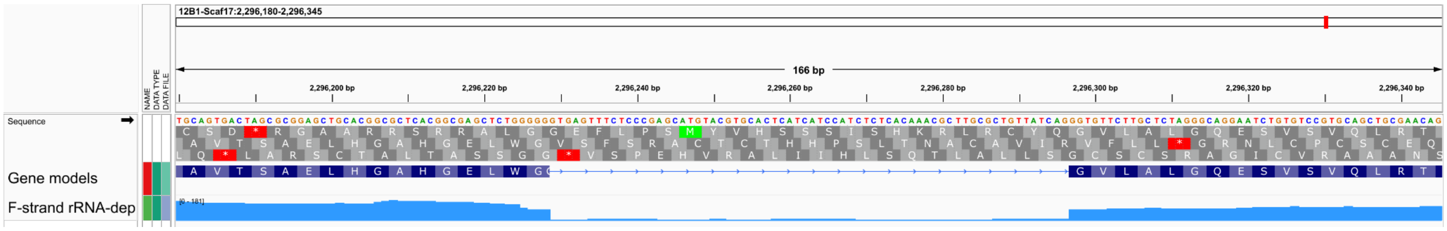


#### D - PKZILLA-1 Intron #4

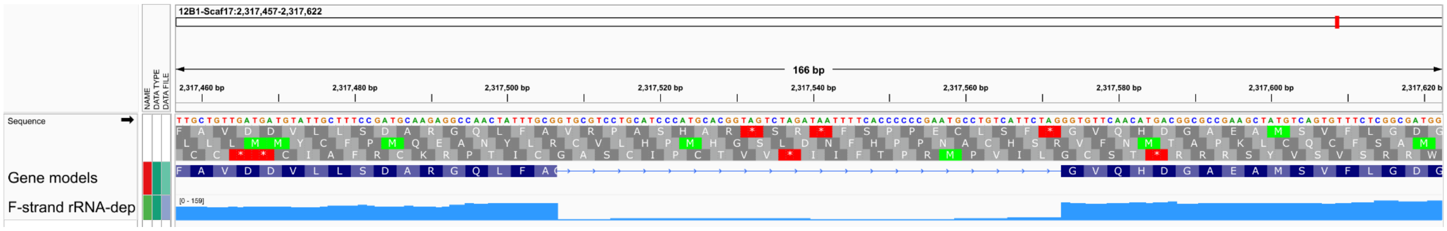


#### E - PKZILLA-1 Intron #5

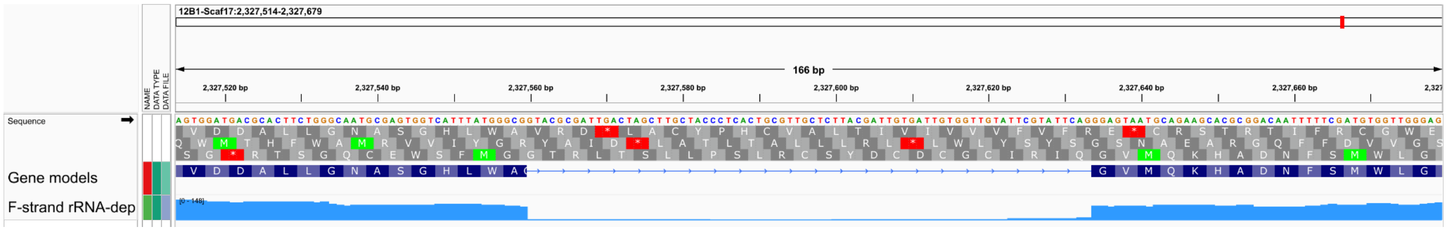


#### F - PKZILLA-1 Intron #6

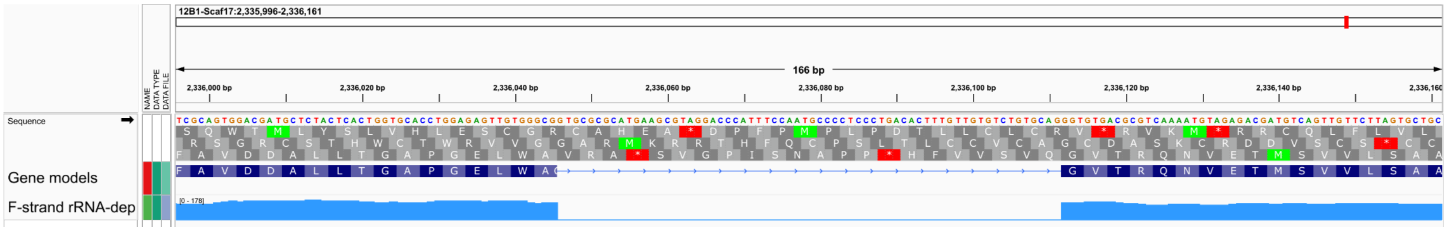

#### G - PKZILLA-1 Intron #7

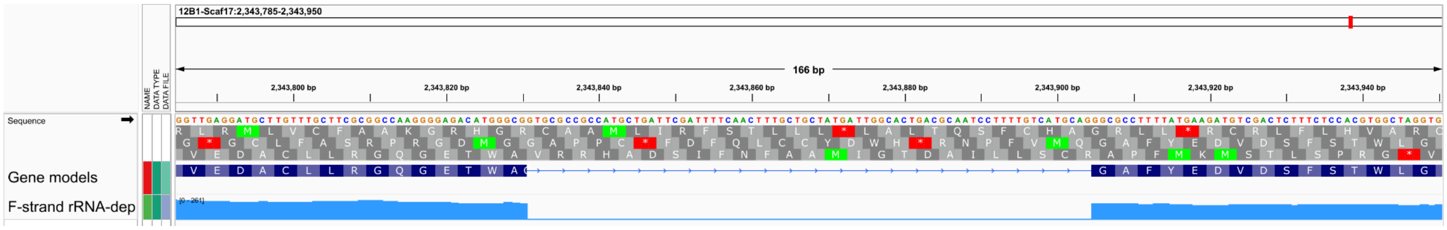


#### H - PKZILLA-1 Intron #8

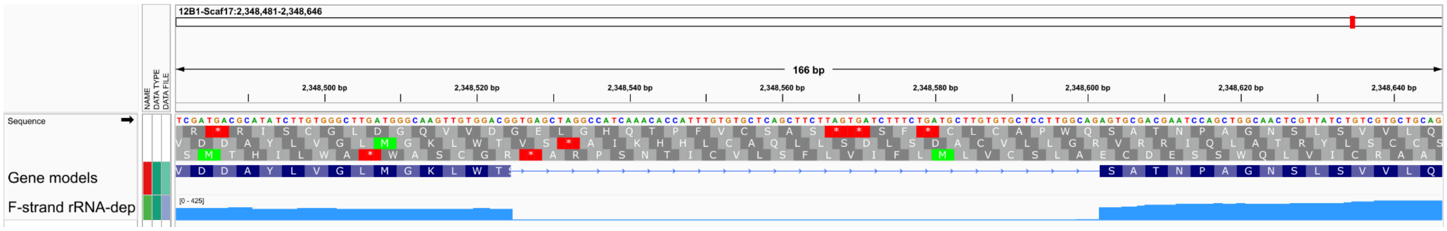


#### I - PKZILLA-1 Intron #9

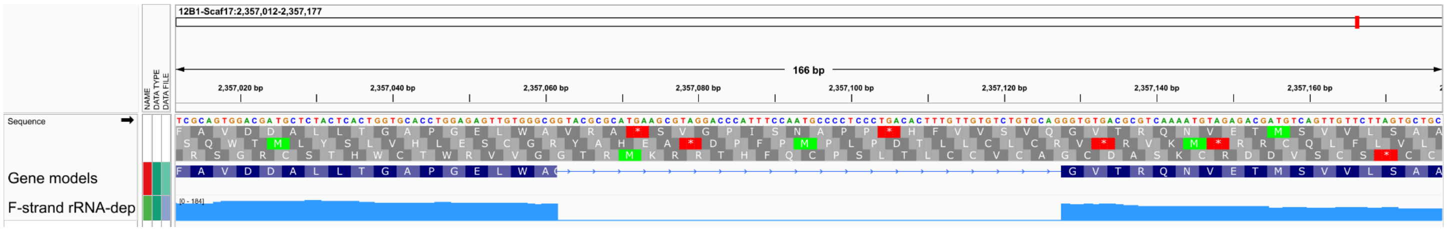


#### J - PKZILLA-1 Intron #10

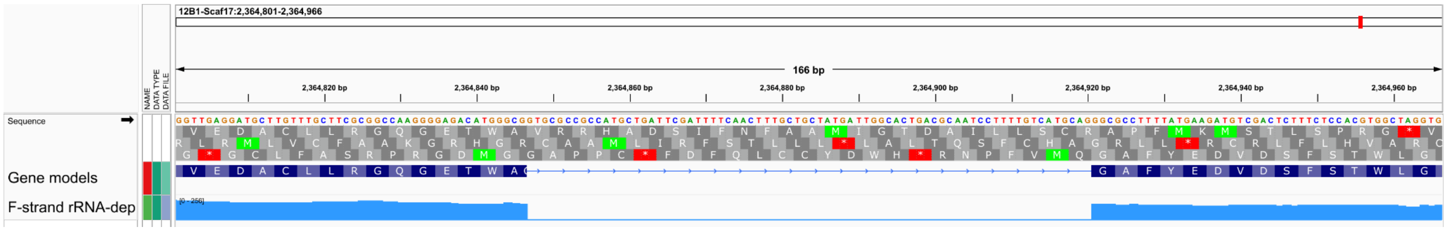


#### K - PKZILLA-1 Intron #11

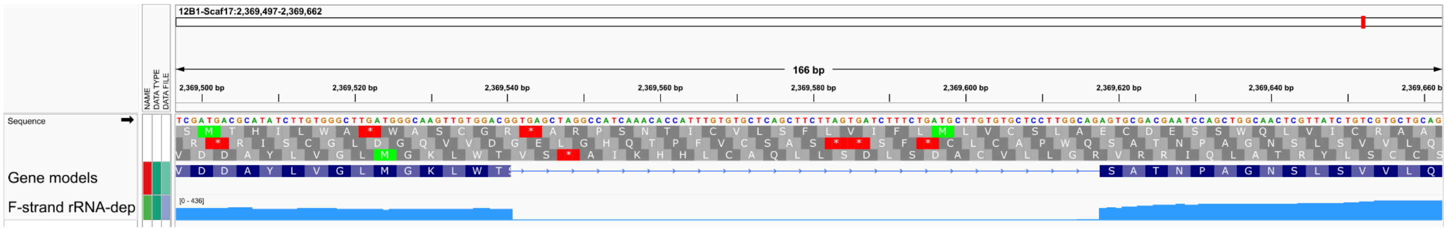


#### L - PKZILLA-1 Intron #12

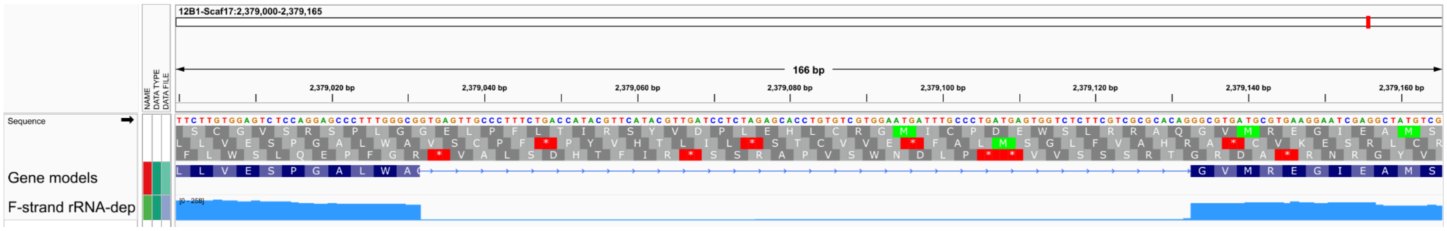


#### M - PKZILLA-1 Intron #13

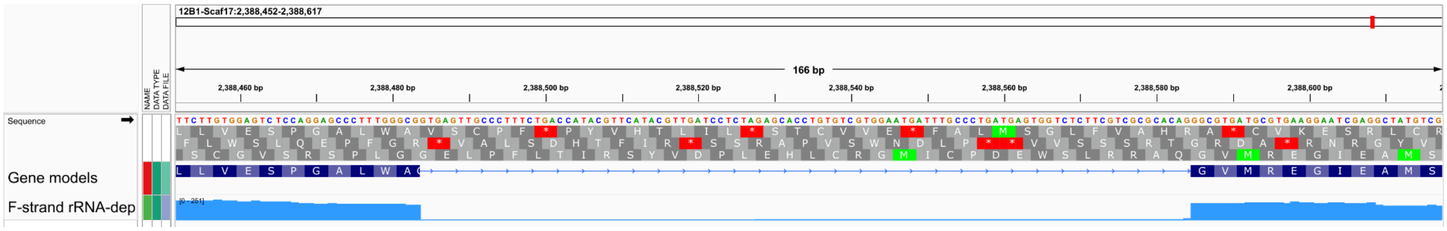

#### N - PKZILLA-1 Intron #14

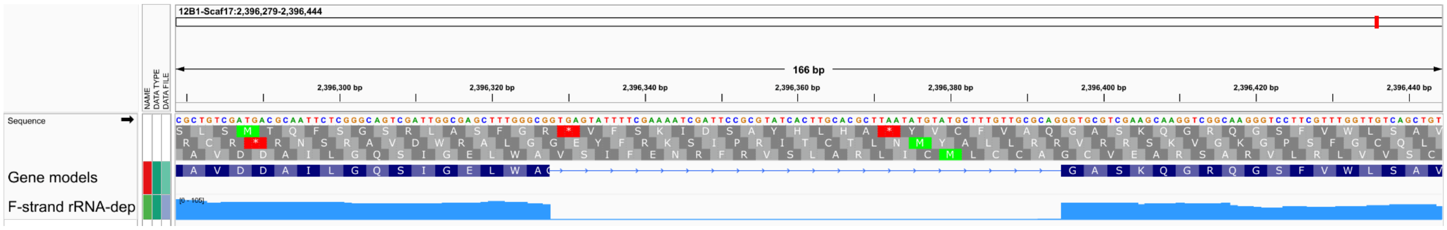


#### O - PKZILLA-1 Intron #15

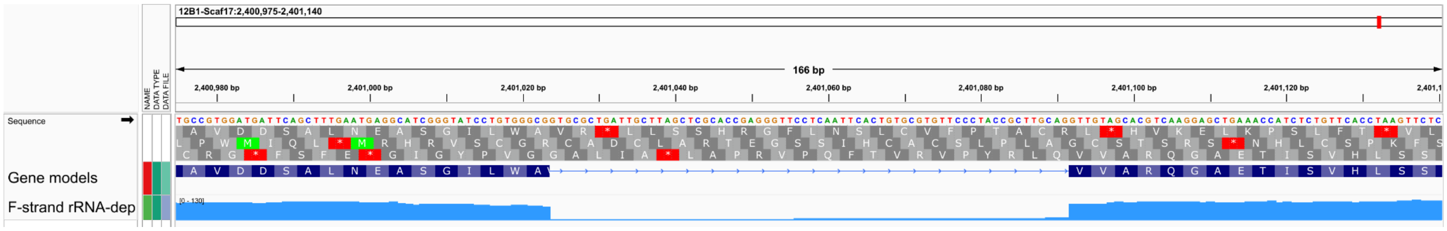


#### P - PKZILLA-1 Intron #16

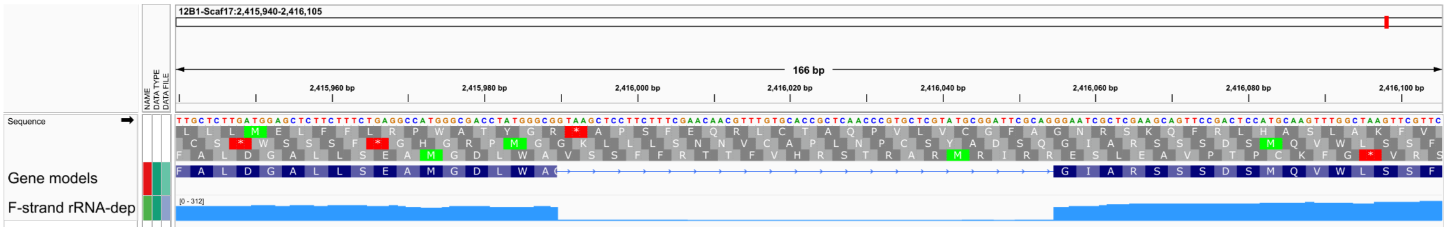


**Fig. S3:** Detail view of PKZILLA-1 introns. Note the drop in aligned rRNA depletion RNA-Seq coverage, and presence of canonical GT-AG splice donor and acceptor sites. Figure produced with Integrated Genomics Viewer (IGV) v2.16.2 (*51*).

#### A - PKZILLA-2 Intron #1

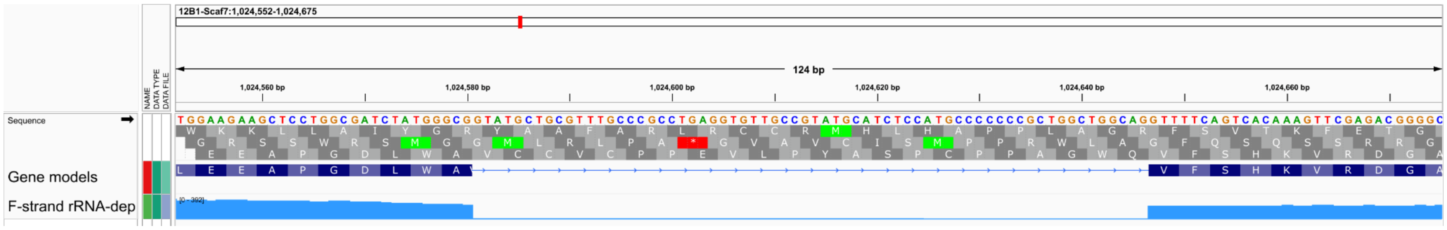


#### B - PKZILLA-2 Intron #2

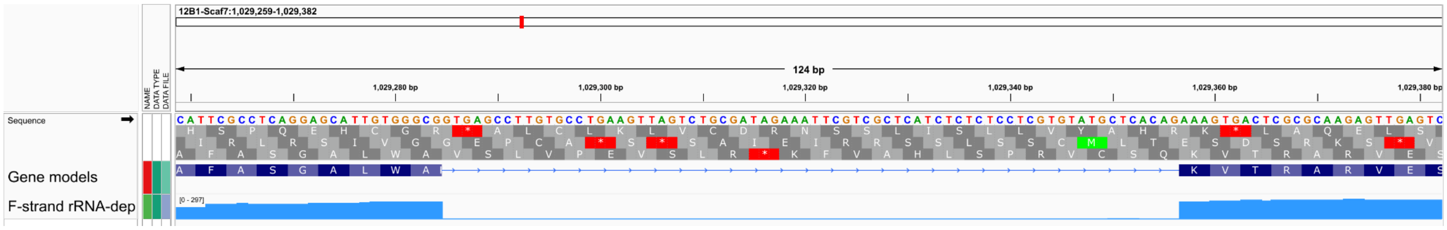


#### C - PKZILLA-2 Intron #3

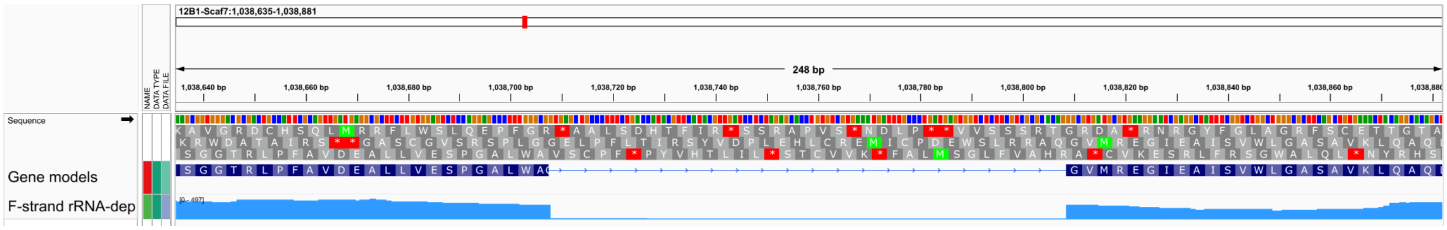


#### D - PKZILLA-2 Intron #4

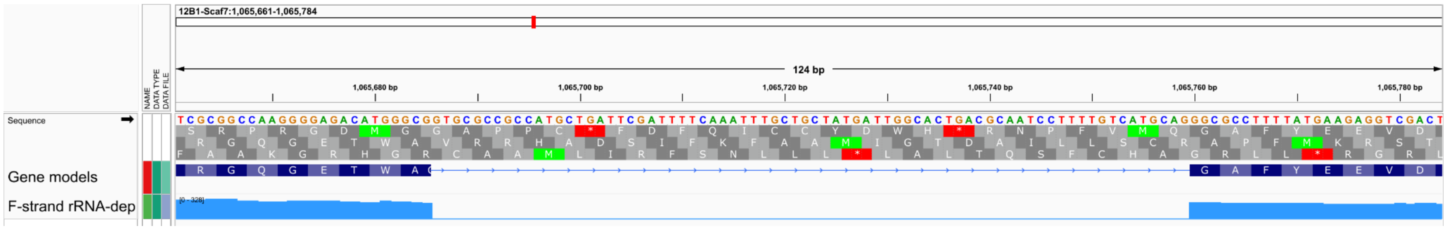


#### E - PKZILLA-2 Intron #5

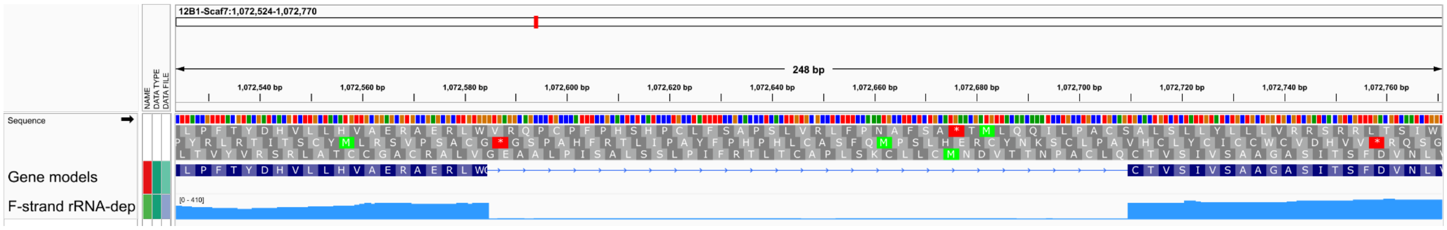


#### F - PKZILLA-2 Intron #6

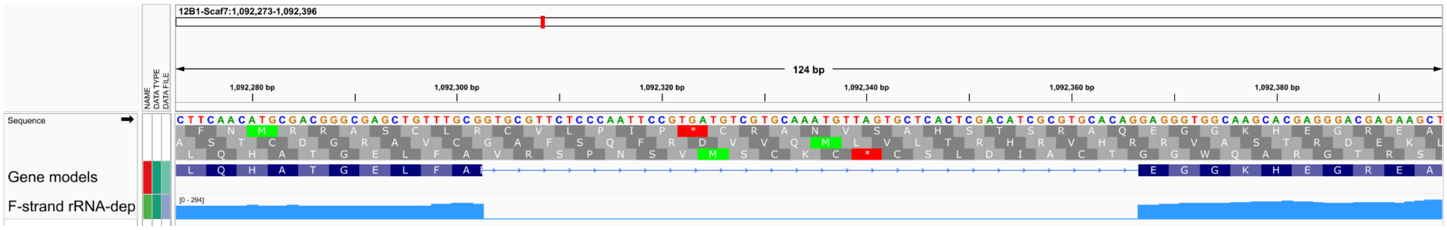

#### G - PKZILLA-2 Intron #7

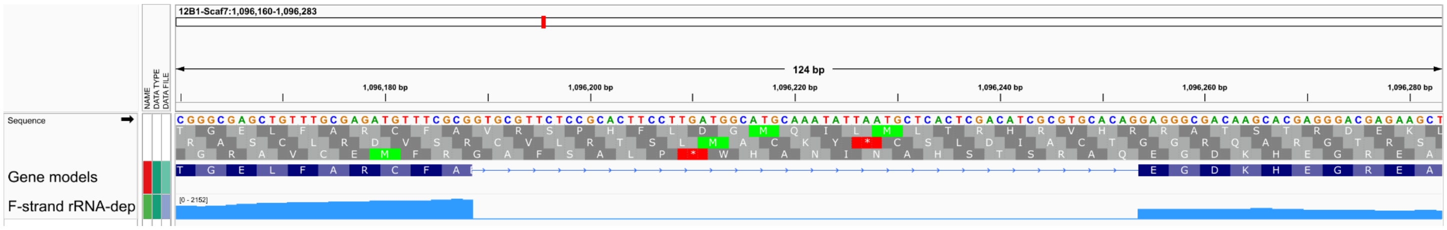


#### H - PKZILLA-2 Intron #8

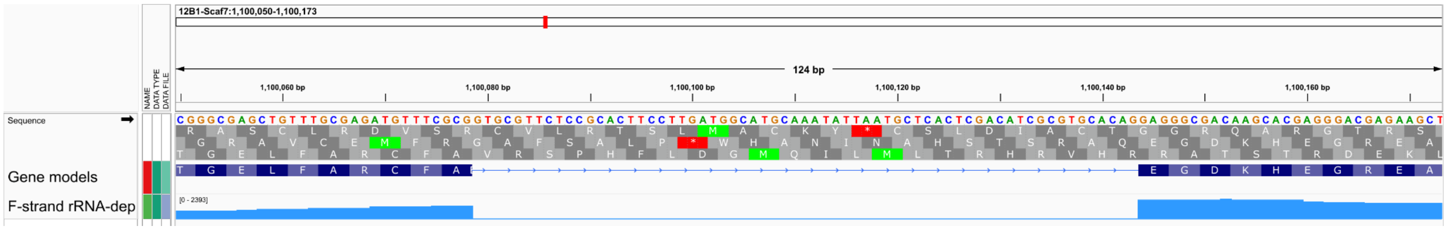


#### I - PKZILLA-2 Intron #9

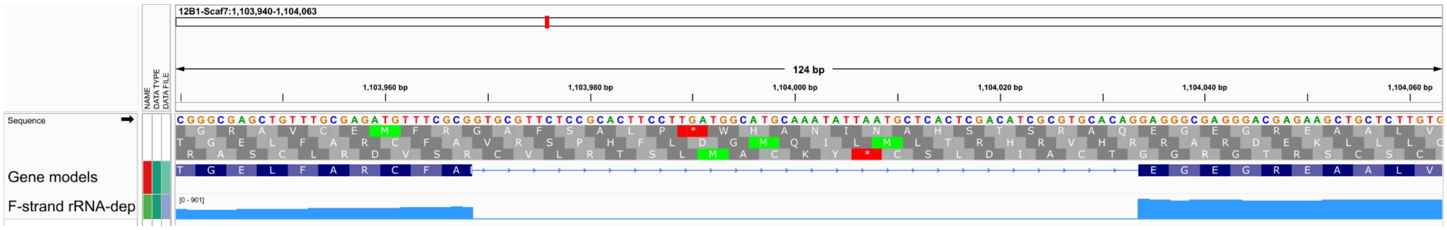


#### J - PKZILLA-2 Intron #10

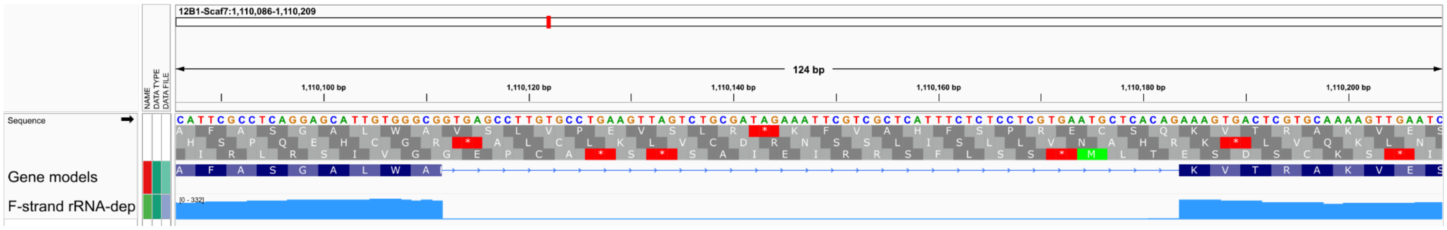


#### K - PKZILLA-2 Intron #11

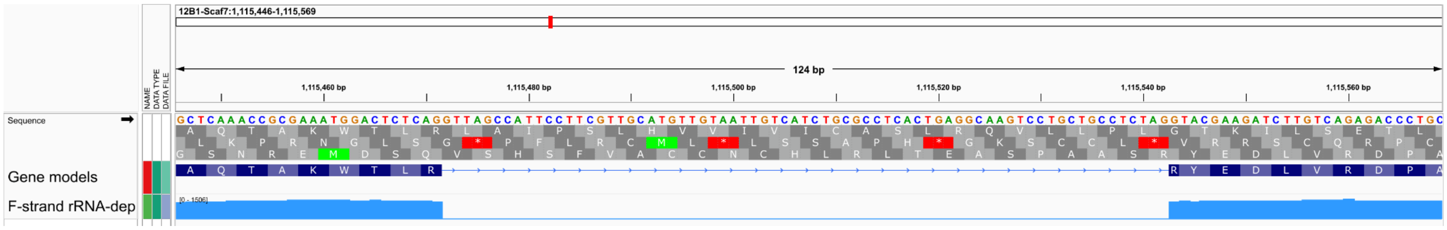


**Fig. S4:** Detail view of PKZILLA-2 introns. Note the drop in aligned rRNA depletion RNA-Seq coverage, and presence of canonical GT-AG splice donor and acceptor sites. Figure produced with Integrated Genomics Viewer (IGV) v2.16.2 (*51*).

#### A - PKZILLA-3 Intron #1

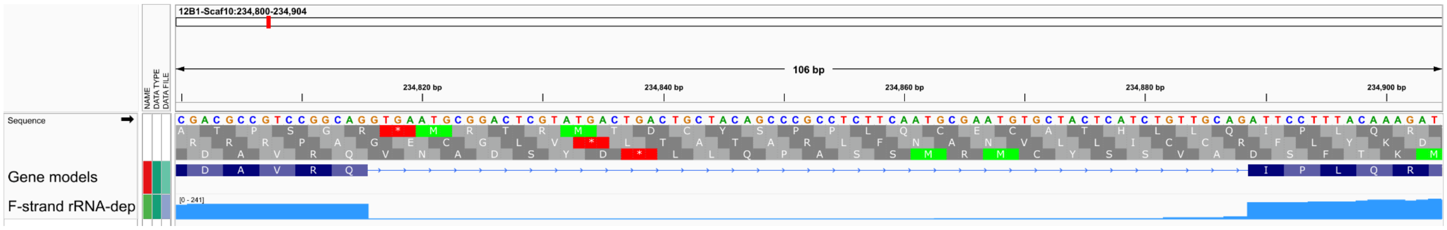


#### B - PKZILLA-3 Intron #2

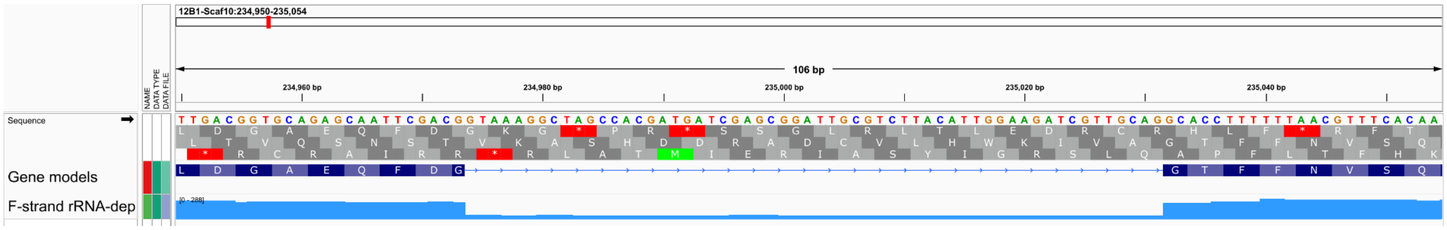


#### C - PKZILLA-3 Intron #3

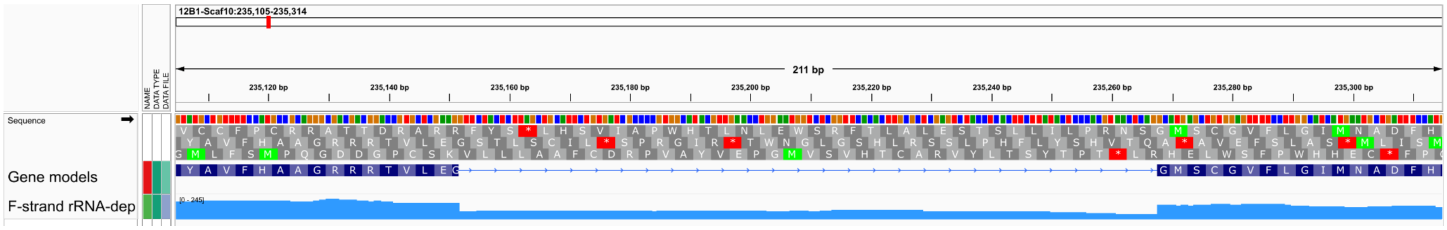


#### D - PKZILLA-3 Intron #4

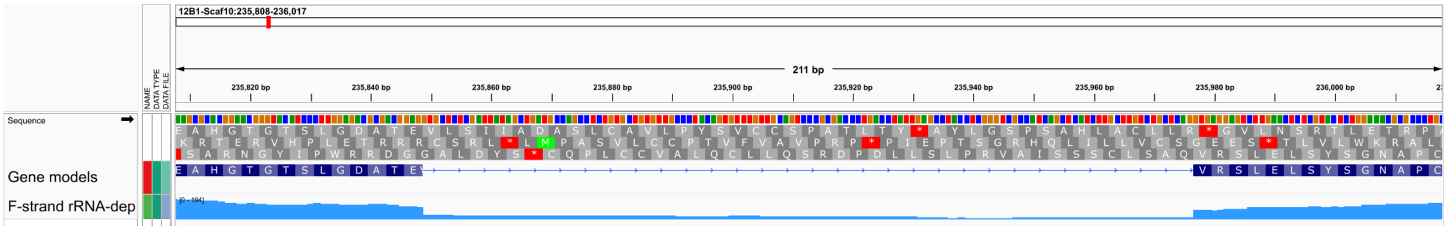


#### E - PKZILLA-3 Intron #5

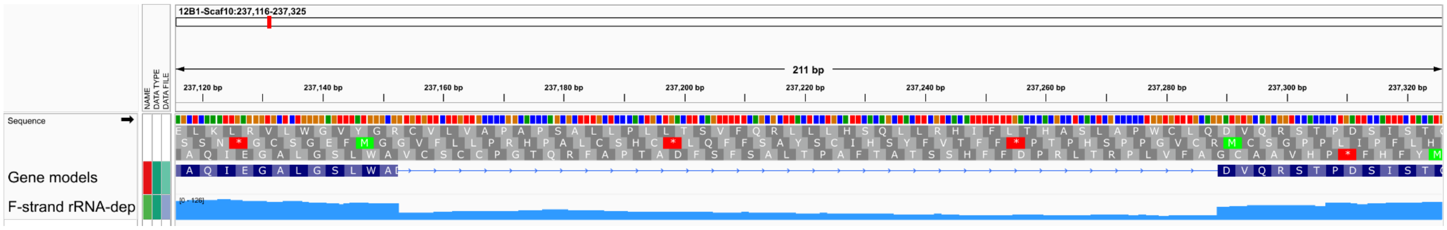


#### F - PKZILLA-3 Intron #6

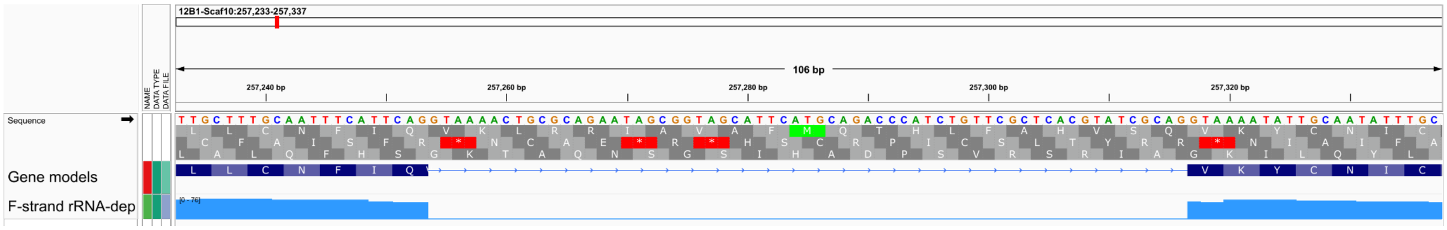

**G - PKZILLA-3 Intron #7**

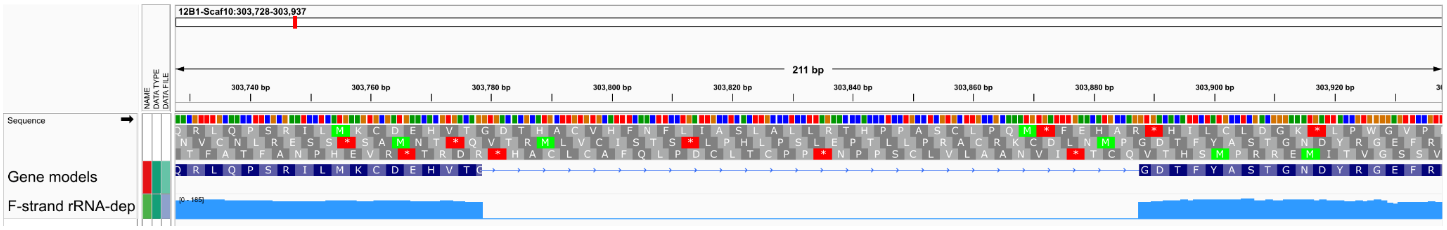


**Fig. S5:** Detail view of PKZILLA-3 introns. Note the drop in aligned rRNA depletion RNA-Seq coverage, and presence of canonical GT-AG splice donor and acceptor sites. Figure produced with Integrated Genomics Viewer (IGV) v2.16.2 (*51*).

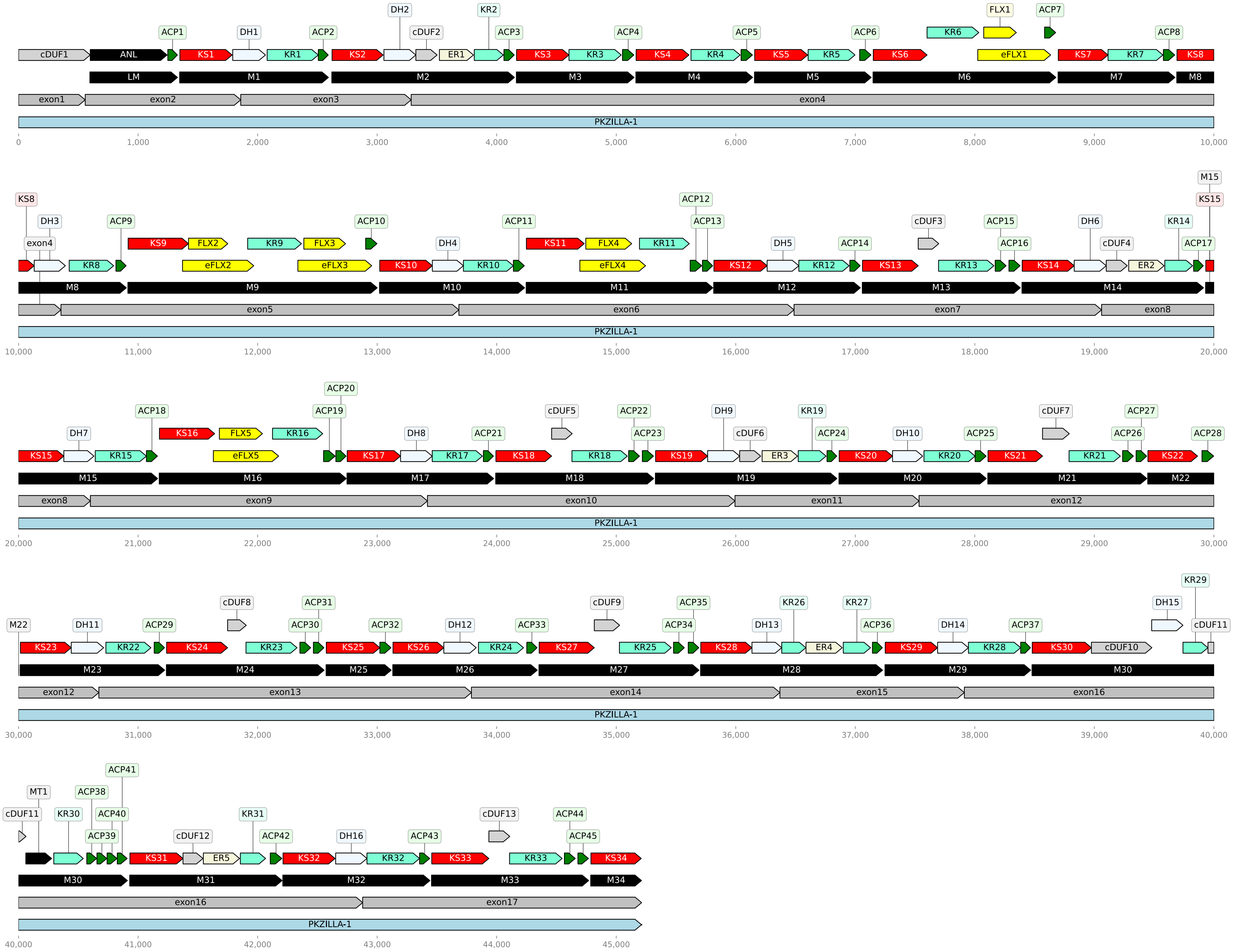

DH2 KR2 FLX1 ACP7
ACP1 DH1 ACP2 cDUF2 ACP3 ACP4 ACP5 ACP6 KR6 ACP8
cDUF1 ANL KS1 KR1 KS2 ER1 KS3 KR3 KS4 KR4 KS5 KR5 KS6 eFLX1 KS7 KR7 KS8
LM M1 M2 M3 M4 M5 M6 M7 M8
exon1 exon2 exon3 exon4
PKZILLA-1
0 1,000 2,000 3,000 4,000 5,000 6,000 7,000 8,000 9,000 10,000
M15
KS8 ACP12 KS15
DH3 ACP9 ACP10 ACP11 ACP13 cDUF3 ACP15 DH6 KR14
exon4 KS9 FLX2 KR9 FLX3 DH4 KS11 FLX4 KR11 DH5 ACP14 ACP16 cDUF4 ACP17
KR8 eFLX2 eFLX3 KS10 KR10 eFLX4 KS12 KR12 KS13 KR13 KS14 ER2
M8 M9 M10 M11 M12 M13 M14
exon5 exon6 exon7 exon8
PKZILLA-1
10,000 11,000 12,000 13,000 14,000 15,000 16,000 17,000 18,000 19,000 20,000
ACP20
ACP18 ACP19 cDUF5 ACP22 DH9 KR19 cDUF7 ACP27
DH7 DH8 ACP21 ACP23 cDUF6 ACP24 DH10 ACP25 ACP26 ACP28
KS16 FLX5 KR16
eFLX5
KS15 KR15 KS17 KR17 KS18 KR18 KS19 ER3 KS20 KR20 KS21 KR21 KS22
M15 M16 M17 M18 M19 M20 M21 M22
exon8 exon9 exon10 exon11 exon12
PKZILLA-1
20,000 21,000 22,000 23,000 24,000 25,000 26,000 27,000 28,000 29,000 30,000
KR29
cDUF8 ACP31 cDUF9 ACP35 KR26 KR27 DH15
M22 DH11 ACP29 ACP30 ACP32 DH12 ACP33 ACP34 DH13 ACP36 DH14 ACP37 cDUF11
KS23 KR22 KS24 KR23 KS25 KS26 KR24 KS27 KR25 KS28 ER4 KS29 KR28 KS30 cDUF10
M23 M24 M25 M26 M27 M28 M29 M30
exon12 exon13 exon14 exon15 exon16
PKZILLA-1
30,000 31,000 32,000 33,000 34,000 35,000 36,000 37,000 38,000 39,000 40,000
ACP41
MT1 ACP38
cDUF11 KR30 ACP40 KR31 cDUF13 ACP44
ACP39 cDUF12 ACP42 DH16 ACP43 ACP45
KS31 ER5 KS32 KR32 KS33 KR33 KS34
M30 M31 M32 M33 M34
exon16 exon17
PKZILLA-1
40,000 41,000 42,000 43,000 44,000 45,000

**Fig. S6:** Graphical plot of non-length-normalized PKZILLA-1 polypeptide (light blue), PKS modules (black), domains (varied colors), and unannotated cDUFs (light gray), and the relationship of shown regions with their source exons (dark gray). Plot generated with DNA Features Viewer (*77*). Source files, scripts, and resulting plots available on Zenodo (doi:10.5281/zenodo.10093617).

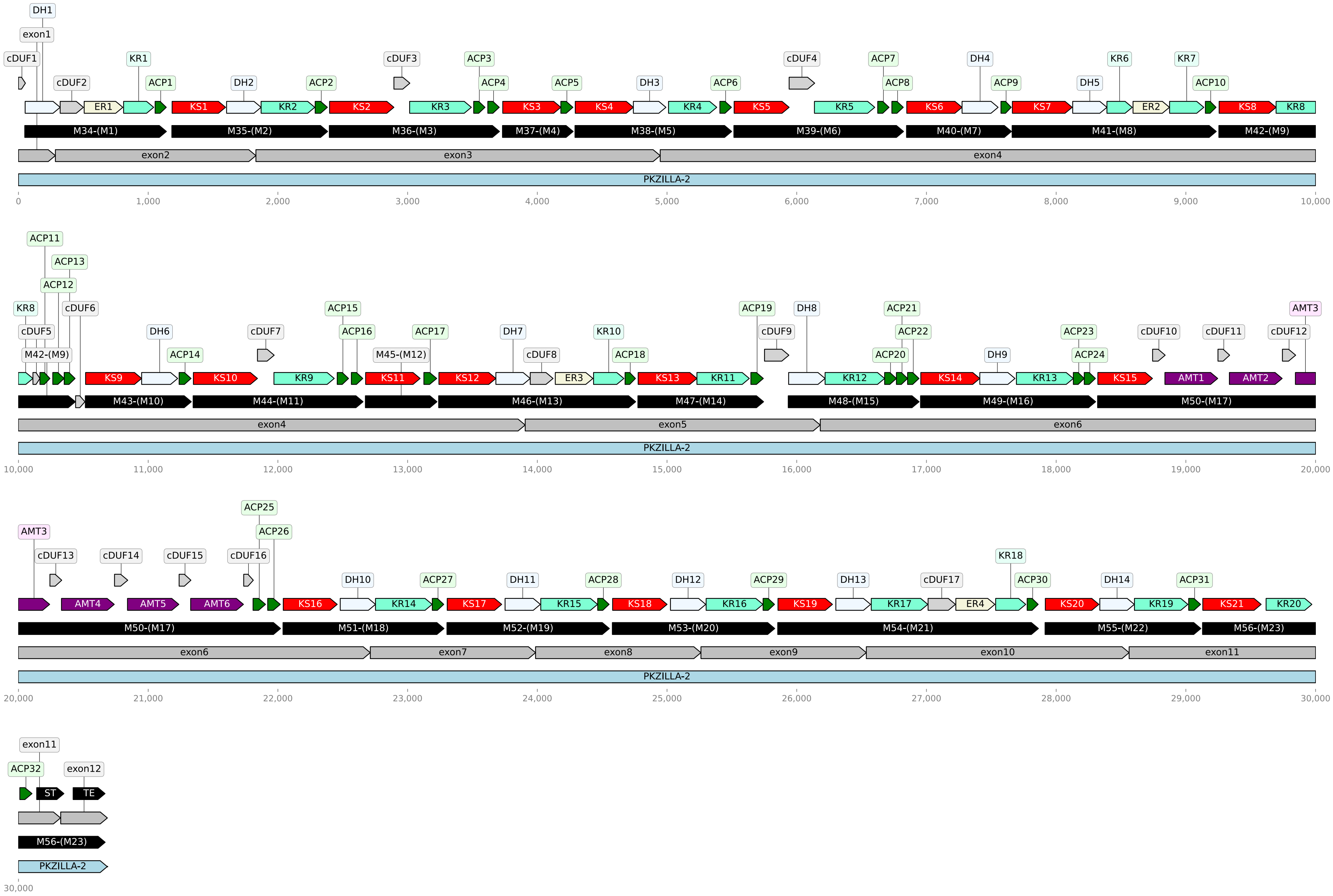

DH1
exon1
cDUF1
cDUF2
KR1
ACP1
DH2
ACP2
cDUF3
ACP3
ACP4
ACP5
DH3
ACP6
cDUF4
ACP7
ACP8
DH4
ACP9
DH5
KR6
KR7
ACP10
ER1
KS1
KR2
KS2
KR3
KS3
KS4
KR4
KS5
KR5
KS6
KS7
ER2
KS8
KR8
M34-(M1)
M35-(M2)
M36-(M3)
M37-(M4)
M38-(M5)
M39-(M6)
M40-(M7)
M41-(M8)
M42-(M9)
exon2
exon3
exon4
PKZILLA-2
0
1,000
2,000
3,000
4,000
5,000
6,000
7,000
8,000
9,000
10,000
ACP11
ACP13
ACP12
KR8
cDUF6
cDUF5
M42-(M9)
DH6
ACP14
cDUF7
ACP15
ACP16
M45-(M12)
ACP17
DH7
cDUF8
KR10
ACP18
ACP19
DH8
cDUF9
ACP21
ACP22
ACP20
DH9
ACP23
ACP24
cDUF10
cDUF11
AMT3
cDUF12
KS9
KS10
KR9
KS11
KS12
ER3
KS13
KR11
KR12
KS14
KR13
KS15
AMT1
AMT2
M43-(M10)
M44-(M11)
M46-(M13)
M47-(M14)
M48-(M15)
M49-(M16)
M50-(M17)
exon4
exon5
exon6
PKZILLA-2
10,000
11,000
12,000
13,000
14,000
15,000
16,000
17,000
18,000
19,000
20,000
ACP25
ACP26
AMT3
cDUF13
cDUF14
cDUF15
cDUF16
DH10
ACP27
DH11
ACP28
DH12
ACP29
DH13
cDUF17
KR18
ACP30
DH14
ACP31
AMT4
AMT5
AMT6
KS16
KR14
KS17
KR15
KS18
KR16
KS19
KR17
ER4
KS20
KR19
KS21
KR20
M50-(M17)
M51-(M18)
M52-(M19)
M53-(M20)
M54-(M21)
M55-(M22)
M56-(M23)
exon6
exon7
exon8
exon9
exon10
exon11
PKZILLA-2
20,000
21,000
22,000
23,000
24,000
25,000
26,000
27,000
28,000
29,000
30,000
exon11
ACP32
exon12
ST
TE
M56-(M23)
PKZILLA-2
30,000

**Fig. S7:** Graphical plot of non-length-normalized PKZILLA-2 polypeptide (light blue), PKS modules (black), domains (varied colors), and unannotated cDUFs (light gray), and the relationship of shown regions with their source exons (dark gray). Plot generated with DNA Features Viewer (*77*). Source files, scripts, and resulting plots available on Zenodo (doi:10.5281/zenodo.10093617).

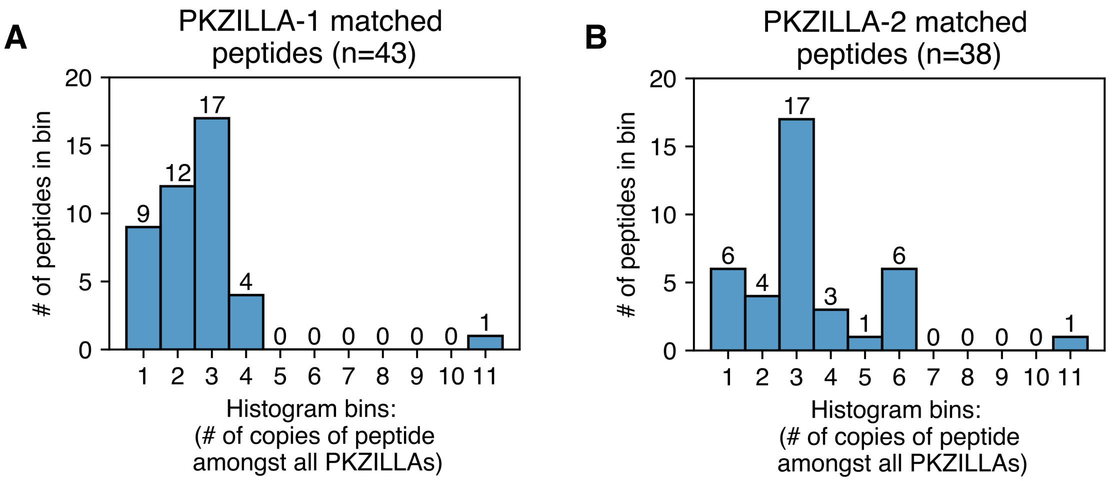


**Fig. S8:** Histogram of proteomics detected peptides binned by the number of times the specific peptide occurs within the PKZILLA polypeptides of the *Prymnesium parvum* 12B1 proteome. The majority of detected peptides from the PKZILLAs are present >=2 times. These peptides were only present in the PKZILLA gene models and were not found anywhere else in 6-frame translations of the 12B1 genome (doi:10.5281/zenodo.10028959). Source data, workflow, and plots available on Zenodo (doi:10.5281/zenodo.10028959).

**A** # of confident peptides matching **PKZILLA-1**: 43

- # of protein-unique peptides: 30
  - # of exon-unique: 14
    - # of single-match: 9
    - # of multimatch: 5
  - # of exon-multimatch: 16
- # of protein-multimatch peptides: 13

**B** # of confident peptides matching **PKZILLA-2**: 38

- # of protein-unique peptides: 25
  - # of exon-unique: 16
    - # of single-match: 6
    - # of multimatch: 10
  - # of exon-multimatch: 9
- # of protein-multimatch peptides: 13

**Fig. S9**: Hierarchical classification of proteomics detected peptides. **(A)** PKZILLA-1, source values from Zenodo file 'PKZILLA-1_classify_peptides.txt' and **(B)** PKZILLA-2, source values from Zenodo file 'PKZILLA-2_classify_peptides.txt'. Source data, workflow, and these derived data available on Zenodo (doi:10.5281/zenodo.10028959).

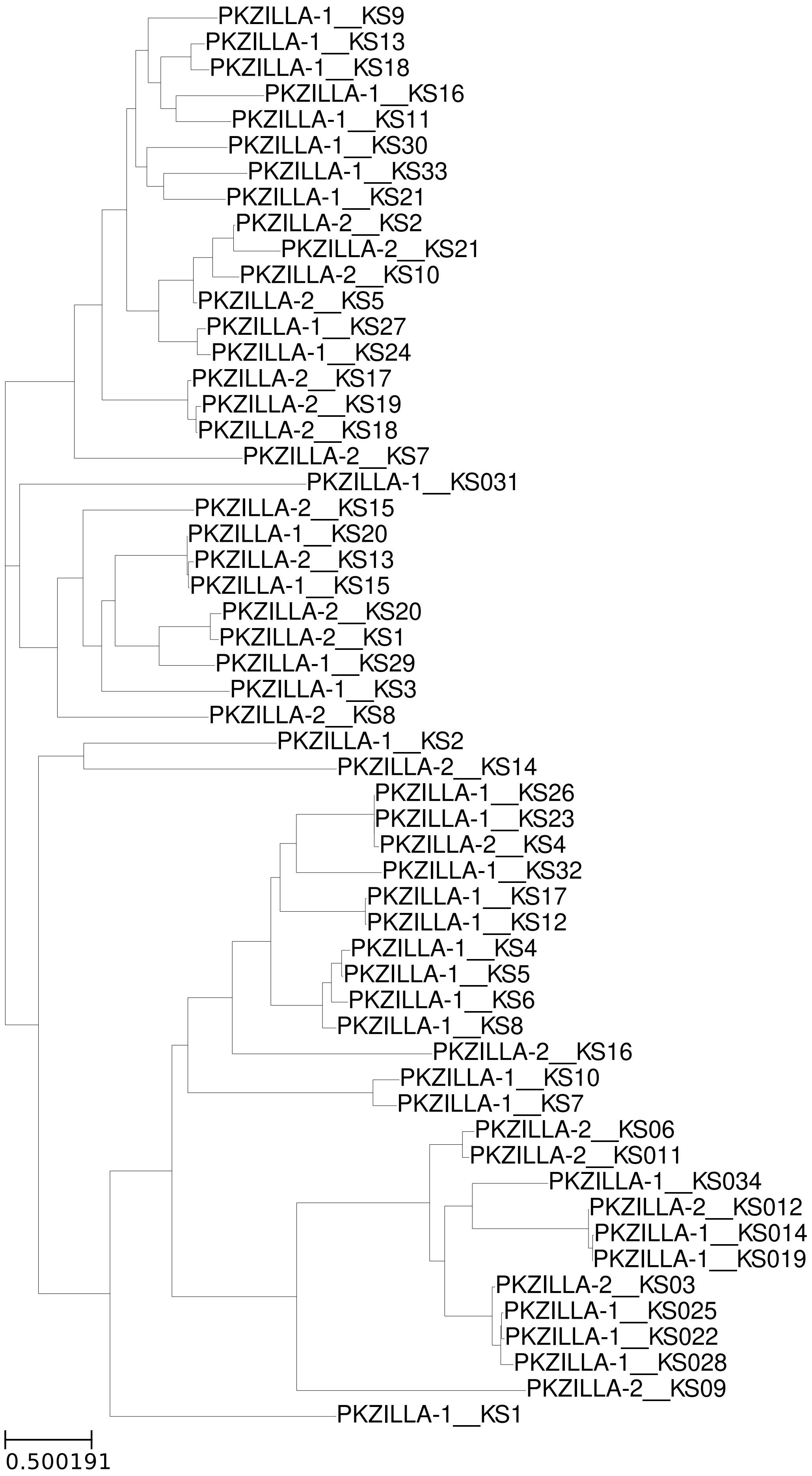
PKZILLA-1__KS9
PKZILLA-1__KS13
PKZILLA-1__KS18
PKZILLA-1__KS16
PKZILLA-1__KS11
PKZILLA-1__KS30
PKZILLA-1__KS33
PKZILLA-1__KS21
PKZILLA-2__KS2
PKZILLA-2__KS21
PKZILLA-2__KS10
PKZILLA-2__KS5
PKZILLA-1__KS27
PKZILLA-1__KS24
PKZILLA-2__KS17
PKZILLA-2__KS19
PKZILLA-2__KS18
PKZILLA-2__KS7
PKZILLA-1__KS031
PKZILLA-2__KS15
PKZILLA-1__KS20
PKZILLA-2__KS13
PKZILLA-1__KS15
PKZILLA-2__KS20
PKZILLA-2__KS1
PKZILLA-1__KS29
PKZILLA-1__KS3
PKZILLA-2__KS8
PKZILLA-1__KS2
PKZILLA-2__KS14
PKZILLA-1__KS26
PKZILLA-1__KS23
PKZILLA-2__KS4
PKZILLA-1__KS32
PKZILLA-1__KS17
PKZILLA-1__KS12
PKZILLA-1__KS4
PKZILLA-1__KS5
PKZILLA-1__KS6
PKZILLA-1__KS8
PKZILLA-2__KS16
PKZILLA-1__KS10
PKZILLA-1__KS7
PKZILLA-2__KS06
PKZILLA-2__KS011
PKZILLA-1__KS034
PKZILLA-2__KS012
PKZILLA-1__KS014
PKZILLA-1__KS019
PKZILLA-2__KS03
PKZILLA-1__KS025
PKZILLA-1__KS022
PKZILLA-1__KS028
PKZILLA-2__KS09
PKZILLA-1__KS1
0.500191

**Fig. S10:** Phylogenetic analysis of PKZILLA ketosynthase (KS) and non-elongating KS (KS0) domains. Sequences were multiple sequence aligned with kalign2 (*58*) and the unrooted phylogeny calculated with raxml-ng (*92*) using model LG+G4m without bootstraps. Plots generated with ete3 (*93*). Branch lengths are drawn to scale and indicate the number of substitutions per site. Source data, workflow, results, and plots available on Zenodo (doi:10.5281/zenodo.10152638).

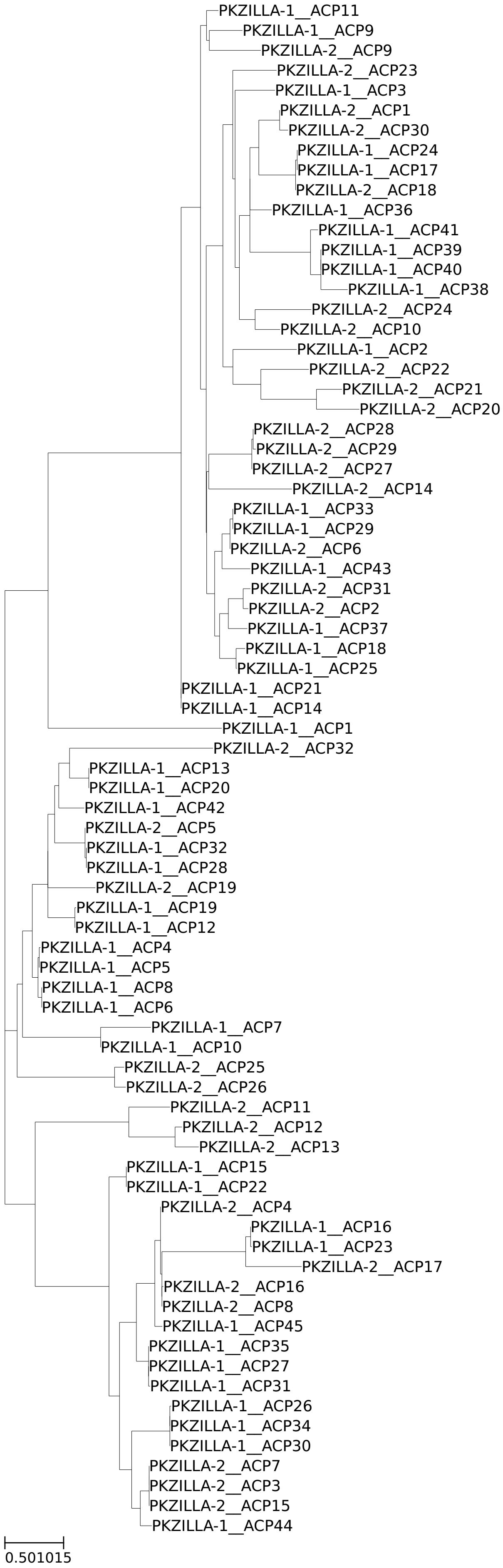
PKZILLA-1__ACP11
PKZILLA-1__ACP9
PKZILLA-2__ACP9
PKZILLA-2__ACP23
PKZILLA-1__ACP3
PKZILLA-2__ACP1
PKZILLA-2__ACP30
PKZILLA-1__ACP24
PKZILLA-1__ACP17
PKZILLA-2__ACP18
PKZILLA-1__ACP36
PKZILLA-1__ACP41
PKZILLA-1__ACP39
PKZILLA-1__ACP40
PKZILLA-1__ACP38
PKZILLA-2__ACP24
PKZILLA-2__ACP10
PKZILLA-1__ACP2
PKZILLA-2__ACP22
PKZILLA-2__ACP21
PKZILLA-2__ACP20
PKZILLA-2__ACP28
PKZILLA-2__ACP29
PKZILLA-2__ACP27
PKZILLA-2__ACP14
PKZILLA-1__ACP33
PKZILLA-1__ACP29
PKZILLA-2__ACP6
PKZILLA-1__ACP43
PKZILLA-2__ACP31
PKZILLA-2__ACP2
PKZILLA-1__ACP37
PKZILLA-1__ACP18
PKZILLA-1__ACP25
PKZILLA-1__ACP21
PKZILLA-1__ACP14
PKZILLA-1__ACP1
PKZILLA-2__ACP32
PKZILLA-1__ACP13
PKZILLA-1__ACP20
PKZILLA-1__ACP42
PKZILLA-2__ACP5
PKZILLA-1__ACP32
PKZILLA-1__ACP28
PKZILLA-2__ACP19
PKZILLA-1__ACP19
PKZILLA-1__ACP12
PKZILLA-1__ACP4
PKZILLA-1__ACP5
PKZILLA-1__ACP8
PKZILLA-1__ACP6
PKZILLA-1__ACP7
PKZILLA-1__ACP10
PKZILLA-2__ACP25
PKZILLA-2__ACP26
PKZILLA-2__ACP11
PKZILLA-2__ACP12
PKZILLA-2__ACP13
PKZILLA-1__ACP15
PKZILLA-1__ACP22
PKZILLA-2__ACP4
PKZILLA-1__ACP16
PKZILLA-1__ACP23
PKZILLA-2__ACP17
PKZILLA-2__ACP16
PKZILLA-2__ACP8
PKZILLA-1__ACP45
PKZILLA-1__ACP35
PKZILLA-1__ACP27
PKZILLA-1__ACP31
PKZILLA-1__ACP26
PKZILLA-1__ACP34
PKZILLA-1__ACP30
PKZILLA-2__ACP7
PKZILLA-2__ACP3
PKZILLA-2__ACP15
PKZILLA-1__ACP44
0.501015

**Fig. S11:** Phylogenetic analysis of PKZILLA acyl carrier protein (ACP) domains. Sequences were multiple sequence aligned with kalign2 (*58*) and the unrooted phylogeny calculated with raxml-ng (*92*) using model LG+G4m without bootstraps. Plots generated with ete3 (*93*). Branch lengths are drawn to scale and indicate the number of substitutions per site. Source data, workflow, results, and plots available on Zenodo (doi:10.5281/zenodo.10152638).

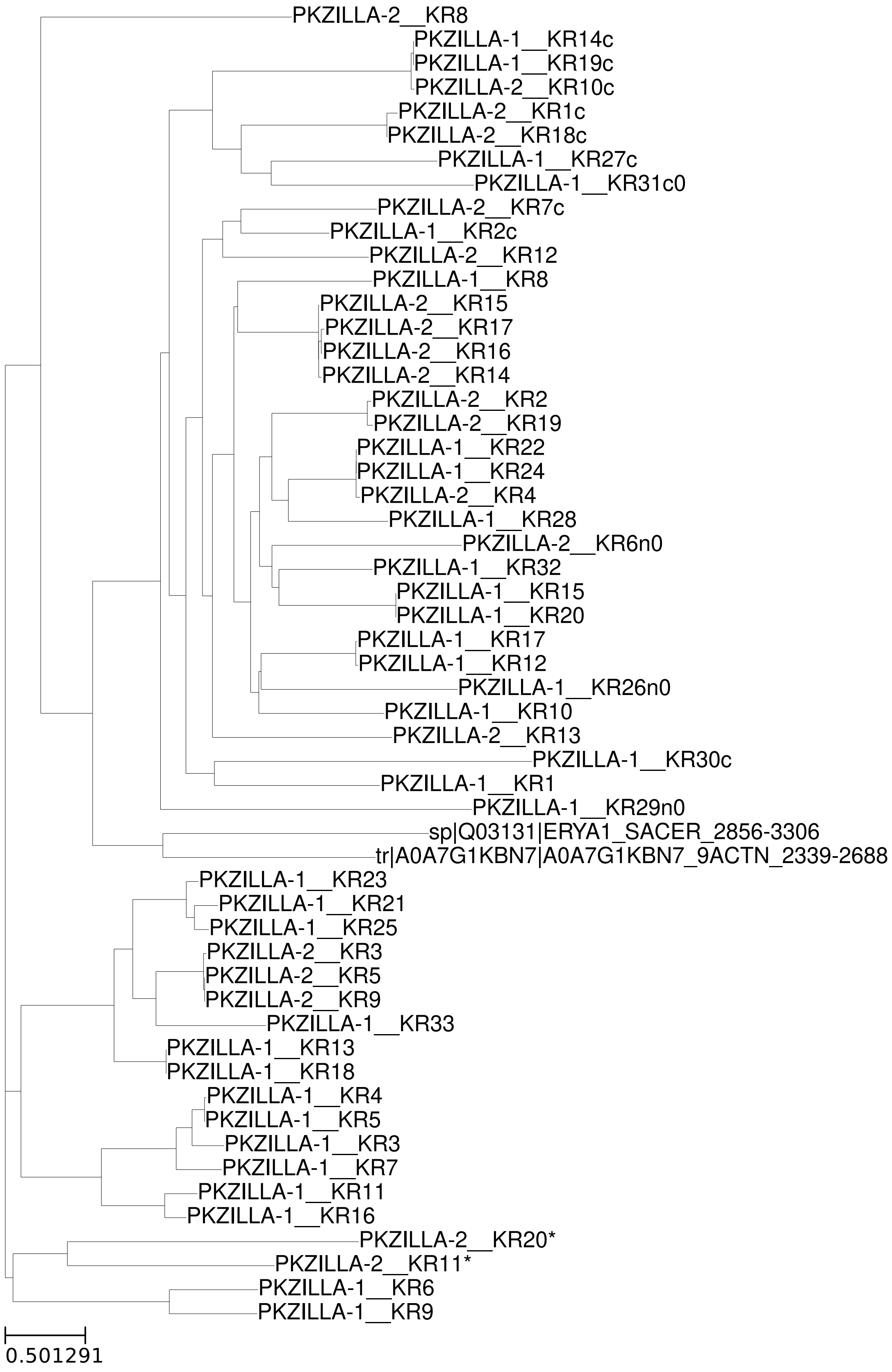
PKZILLA-2__KR8
PKZILLA-1__KR14c
PKZILLA-1__KR19c
PKZILLA-2__KR10c
PKZILLA-2__KR1c
PKZILLA-2__KR18c
PKZILLA-1__KR27c
PKZILLA-1__KR31c0
PKZILLA-2__KR7c
PKZILLA-1__KR2c
PKZILLA-2__KR12
PKZILLA-1__KR8
PKZILLA-2__KR15
PKZILLA-2__KR17
PKZILLA-2__KR16
PKZILLA-2__KR14
PKZILLA-2__KR2
PKZILLA-2__KR19
PKZILLA-1__KR22
PKZILLA-1__KR24
PKZILLA-2__KR4
PKZILLA-1__KR28
PKZILLA-2__KR6n0
PKZILLA-1__KR32
PKZILLA-1__KR15
PKZILLA-1__KR20
PKZILLA-1__KR17
PKZILLA-1__KR12
PKZILLA-1__KR26n0
PKZILLA-1__KR10
PKZILLA-2__KR13
PKZILLA-1__KR30c
PKZILLA-1__KR1
PKZILLA-1__KR29n0
sp|Q03131|ERYA1_SACER_2856-3306
tr|A0A7G1KBN7|A0A7G1KBN7_9ACTN_2339-2688
PKZILLA-1__KR23
PKZILLA-1__KR21
PKZILLA-1__KR25
PKZILLA-2__KR3
PKZILLA-2__KR5
PKZILLA-2__KR9
PKZILLA-1__KR33
PKZILLA-1__KR13
PKZILLA-1__KR18
PKZILLA-1__KR4
PKZILLA-1__KR5
PKZILLA-1__KR3
PKZILLA-1__KR7
PKZILLA-1__KR11
PKZILLA-1__KR16
PKZILLA-2__KR20*
PKZILLA-2__KR11*
PKZILLA-1__KR6
PKZILLA-1__KR9
0.501291

**Fig. S12:** Phylogenetic analysis of PKZILLA ketoreductase (KR) domains and select representative KR domains from Uniprot. Sequences were multiple sequence aligned with kalign2 (*58*) and the unrooted phylogeny calculated with raxml-ng (*92*) using model LG+G4m without bootstraps. Plots generated with ete3 (*93*). Branch lengths are drawn to scale and indicate the number of substitutions per site. KR domains are annotated as KR/KRf, KRn, KRc depending on their respective classification as a full length non-split KR, a split KR N-terminal subdomain, or a split KR C-terminal subdomain, and "0" is appended to the domain name if the active site residues suggest the domain is catalytically inactive. KR* domains would be expected to be active based on the biosynthetic model, but are bioinformatically predicted to be inactive (table S9), and thus may be false positive predictions of inactivity caused by divergent catalytic properties. For further details on catalytic activity determination via active site analysis, see "Analysis of active site residues within PKZILLA PKS domains" in the Materials and Methods and the associated Zenodo item (doi:10.5281/zenodo.10028517). Source data, workflow, results, and plots available on Zenodo (doi:10.5281/zenodo.10152638).

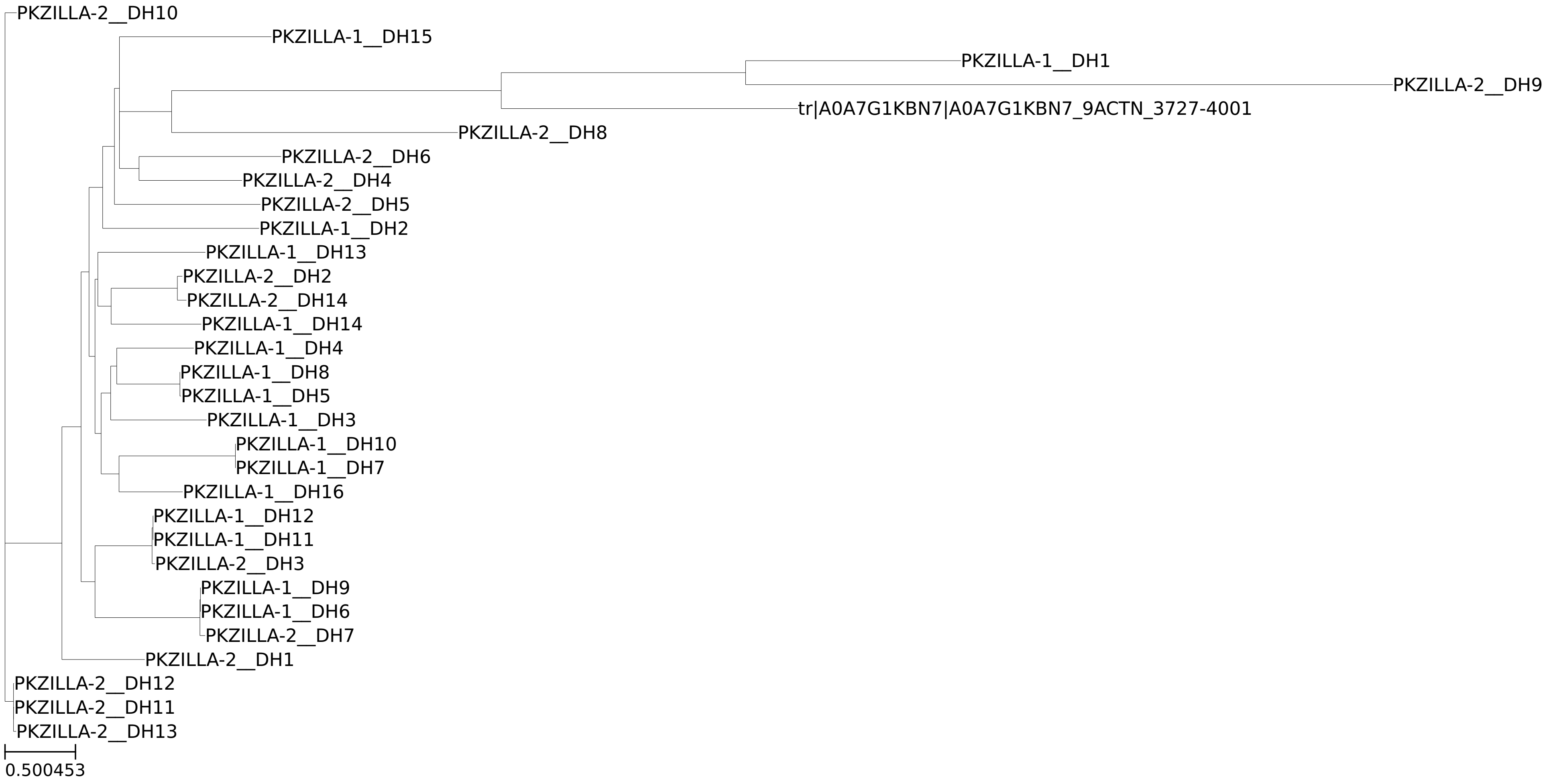


**Fig. S13:** Phylogenetic analysis of PKZILLA dehydratase (DH) domains and select representative DH domains from Uniprot. Sequences were multiple sequence aligned with kalign2 (*58*) and the unrooted phylogeny calculated with raxml-ng (*92*) using model LG+G4m without bootstraps. Plots generated with ete3 (*93*). Branch lengths are drawn to scale and indicate the number of substitutions per site. Source data, workflow, results, and plots available on Zenodo (doi:10.5281/zenodo.10152638)

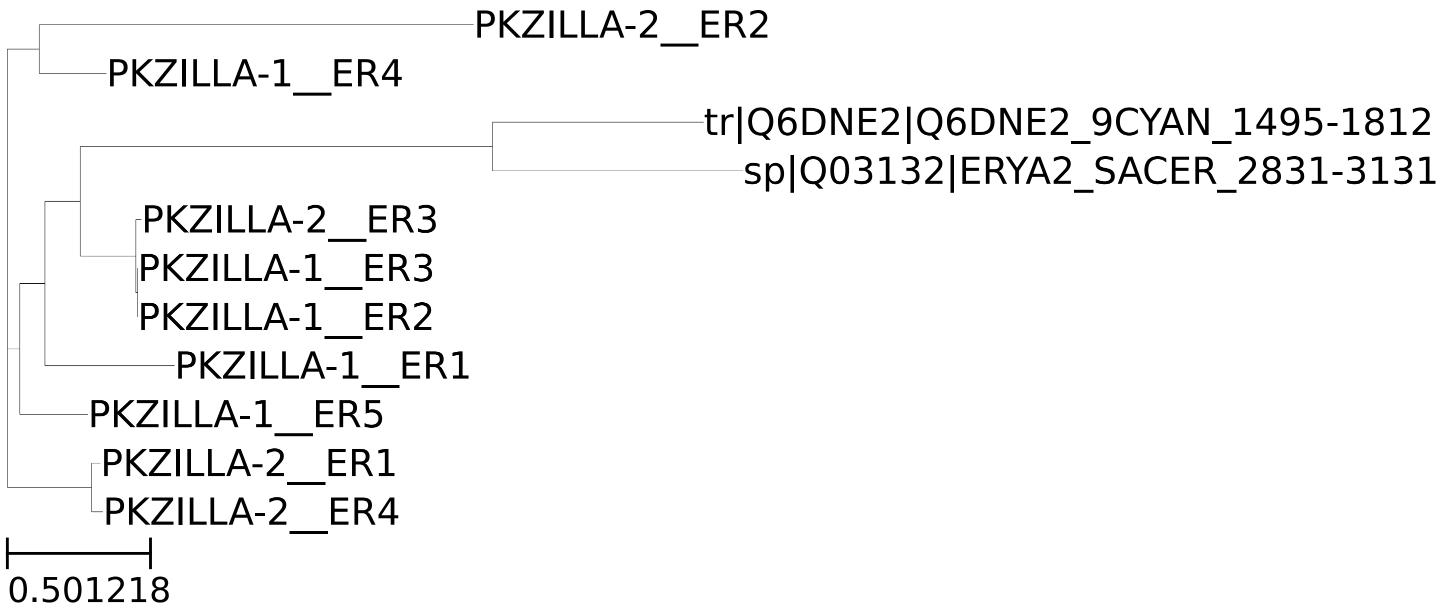


**Fig. S14:** Phylogenetic analysis of PKZILLA enoyl reductase (ER) domains and select representative ER domains from Uniprot. Sequences were multiple sequence aligned with kalign2 (*58*) and the unrooted phylogeny calculated with raxml-ng (*92*) using model LG+G4m without bootstraps. Plots generated with ete3 (*93*). Branch lengths are drawn to scale and indicate the number of substitutions per site. Source data, workflow, results, and plots available on Zenodo (doi:10.5281/zenodo.10152638).

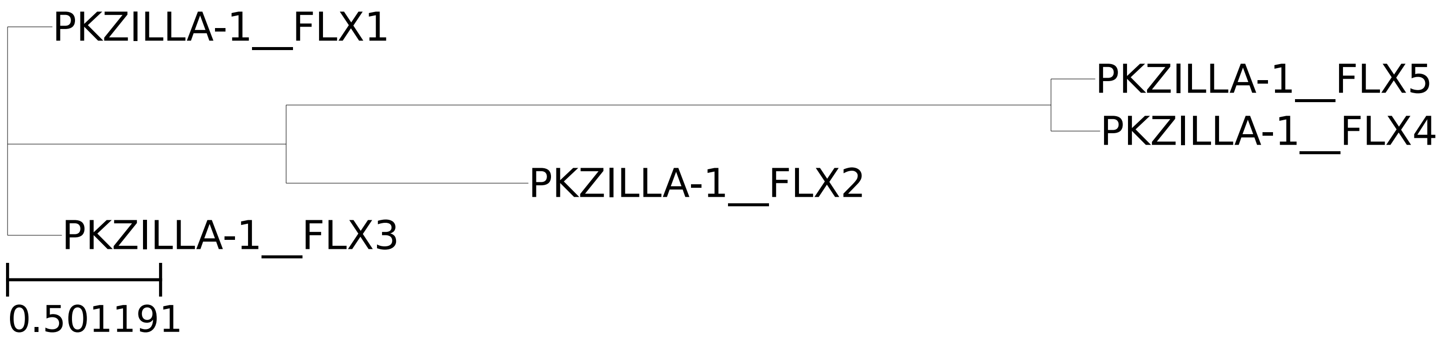


**Fig. S15:** Phylogenetic analysis of PKZILLA flavin oxygenase (FLX) domains. Sequences were multiple sequence aligned with kalign2 (*58*) and the unrooted phylogeny calculated with raxml-ng (*92*) using model LG+G4m without bootstraps. Plots generated with ete3 (*93*). Branch lengths are drawn to scale and indicate the number of substitutions per site. Source data, workflow, results, and plots available on Zenodo (doi:10.5281/zenodo.10152638).

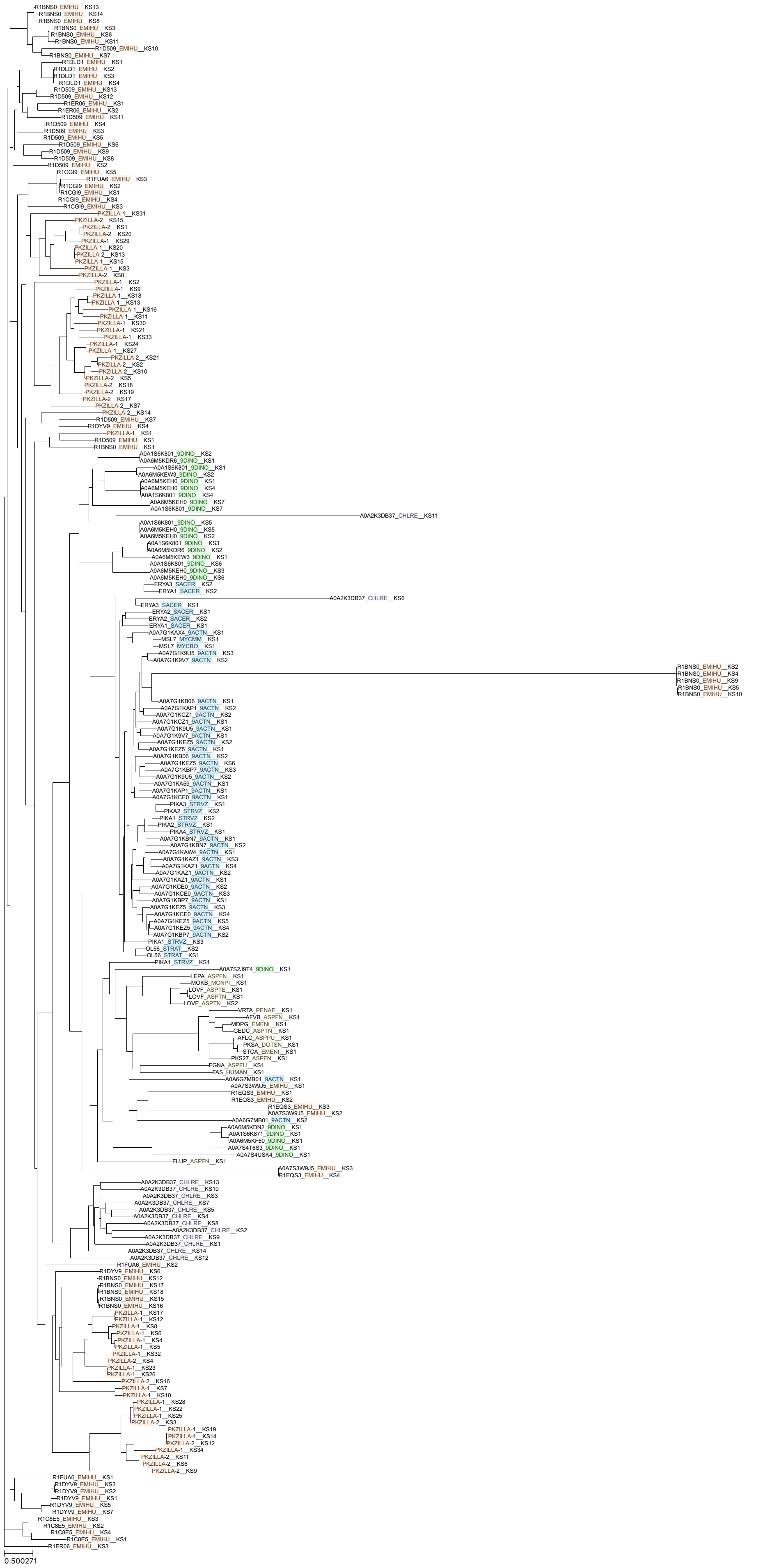
R1BNS0_EMIHU__KS13
R1BNS0_EMIHU__KS14
R1BNS0_EMIHU__KS8
R1BNS0_EMIHU__KS3
R1BNS0_EMIHU__KS6
R1BNS0_EMIHU__KS11
R1D509_EMIHU__KS10
R1BNS0_EMIHU__KS7
R1DLD1_EMIHU__KS1
R1DLD1_EMIHU__KS2
R1DLD1_EMIHU__KS3
R1DLD1_EMIHU__KS4
R1D509_EMIHU__KS13
R1D509_EMIHU__KS12
R1ER06_EMIHU__KS1
R1ER06_EMIHU__KS2
R1D509_EMIHU__KS11
R1D509_EMIHU__KS4
R1D509_EMIHU__KS3
R1D509_EMIHU__KS5
R1D509_EMIHU__KS6
R1D509_EMIHU__KS9
R1D509_EMIHU__KS8
R1D509_EMIHU__KS2
R1CGI9_EMIHU__KS5
R1FUA6_EMIHU__KS3
R1CGI9_EMIHU__KS2
R1CGI9_EMIHU__KS1
R1CGI9_EMIHU__KS4
R1CGI9_EMIHU__KS3
PKZILLA-1__KS31
PKZILLA-2__KS15
PKZILLA-2__KS1
PKZILLA-2__KS20
PKZILLA-1__KS29
PKZILLA-1__KS20
PKZILLA-2__KS13
PKZILLA-1__KS15
PKZILLA-1__KS3
PKZILLA-2__KS8
PKZILLA-1__KS2
PKZILLA-1__KS9
PKZILLA-1__KS18
PKZILLA-1__KS13
PKZILLA-1__KS16
PKZILLA-1__KS11
PKZILLA-1__KS30
PKZILLA-1__KS21
PKZILLA-1__KS33
PKZILLA-1__KS24
PKZILLA-1__KS27
PKZILLA-2__KS21
PKZILLA-2__KS2
PKZILLA-2__KS10
PKZILLA-2__KS5
PKZILLA-2__KS18
PKZILLA-2__KS19
PKZILLA-2__KS17
PKZILLA-2__KS7
PKZILLA-2__KS14
R1D509_EMIHU__KS7
R1DYV9_EMIHU__KS4
PKZILLA-1__KS1
R1D509_EMIHU__KS1
R1BNS0_EMIHU__KS1
A0A1S6K801_9DINO__KS2
A0A6M5KDR6_9DINO__KS1
A0A1S6K801_9DINO__KS1
A0A6M5KEW3_9DINO__KS2
A0A6M5KEH0_9DINO__KS1
A0A6M5KEH0_9DINO__KS4
A0A1S6K801_9DINO__KS4
A0A6M5KEH0_9DINO__KS7
A0A1S6K801_9DINO__KS7
A0A2K3DB37_CHLRE__KS11
A0A1S6K801_9DINO__KS5
A0A6M5KEH0_9DINO__KS5
A0A6M5KEH0_9DINO__KS2
A0A1S6K801_9DINO__KS3
A0A6M5KDR6_9DINO__KS2
A0A6M5KEW3_9DINO__KS1
A0A1S6K801_9DINO__KS6
A0A6M5KEH0_9DINO__KS3
A0A6M5KEH0_9DINO__KS6
ERYA3_SACER__KS2
ERYA1_SACER__KS2
A0A2K3DB37_CHLRE__KS6
ERYA3_SACER__KS1
ERYA2_SACER__KS1
ERYA2_SACER__KS2
ERYA1_SACER__KS1
A0A7G1KAX4_9ACTN__KS1
MSL7_MYCMM__KS1
MSL7_MYCBO__KS1
A0A7G1K9U5_9ACTN__KS3
A0A7G1K9V7_9ACTN__KS2
R1BNS0_EMIHU__KS2
R1BNS0_EMIHU__KS4
R1BNS0_EMIHU__KS9
R1BNS0_EMIHU__KS5
R1BNS0_EMIHU__KS10
A0A7G1KB06_9ACTN__KS1
A0A7G1KAP1_9ACTN__KS2
A0A7G1KCZ1_9ACTN__KS2
A0A7G1KCZ1_9ACTN__KS1
A0A7G1K9U5_9ACTN__KS1
A0A7G1K9V7_9ACTN__KS1
A0A7G1KEZ5_9ACTN__KS2
A0A7G1KEZ5_9ACTN__KS1
A0A7G1KB06_9ACTN__KS2
A0A7G1KEZ5_9ACTN__KS6
A0A7G1KBP7_9ACTN__KS3
A0A7G1K9U5_9ACTN__KS2
A0A7G1KA59_9ACTN__KS1
A0A7G1KAP1_9ACTN__KS1
A0A7G1KCE0_9ACTN__KS1
PIKA3_STRVZ__KS1
PIKA2_STRVZ__KS2
PIKA1_STRVZ__KS2
PIKA2_STRVZ__KS1
PIKA4_STRVZ__KS1
A0A7G1KBN7_9ACTN__KS1
A0A7G1KBN7_9ACTN__KS2
A0A7G1KAW4_9ACTN__KS1
A0A7G1KAZ1_9ACTN__KS3
A0A7G1KAZ1_9ACTN__KS4
A0A7G1KAZ1_9ACTN__KS2
A0A7G1KAZ1_9ACTN__KS1
A0A7G1KCE0_9ACTN__KS2
A0A7G1KCE0_9ACTN__KS3
A0A7G1KBP7_9ACTN__KS1
A0A7G1KEZ5_9ACTN__KS3
A0A7G1KCE0_9ACTN__KS4
A0A7G1KEZ5_9ACTN__KS5
A0A7G1KEZ5_9ACTN__KS4
A0A7G1KBP7_9ACTN__KS2
PIKA1_STRVZ__KS3
OL56_STRAT__KS2
OL56_STRAT__KS1
PIKA1_STRVZ__KS1
A0A7S2J9T4_9DINO__KS1
LEPA_ASPFN__KS1
MOKB_MONPI__KS1
LOVF_ASPTE__KS1
LOVF_ASPTN__KS1
LOVF_ASPTN__KS2
VRTA_PENAE__KS1
AFVB_ASPFN__KS1
MDPG_EMENI__KS1
GEDC_ASPTN__KS1
AFLC_ASPPU__KS1
PKSA_DOTSN__KS1
STCA_EMENI__KS1
PKS27_ASPFN__KS1
FGNA_ASPFU__KS1
FAS_HUMAN__KS1
A0A6G7MB01_9ACTN__KS1
A0A7S3W9J5_EMIHU__KS1
R1EQS3_EMIHU__KS1
R1EQS3_EMIHU__KS2
R1EQS3_EMIHU__KS3
A0A7S3W9J5_EMIHU__KS2
A0A6G7MB01_9ACTN__KS2
A0A6M5KDN2_9DINO__KS1
A0A1S6K871_9DINO__KS1
A0A6M5KF60_9DINO__KS1
A0A7S4T6S3_9DINO__KS1
A0A7S4USK4_9DINO__KS1
FLUP_ASPFN__KS1
A0A7S3W9J5_EMIHU__KS3
R1EQS3_EMIHU__KS4
A0A2K3DB37_CHLRE__KS13
A0A2K3DB37_CHLRE__KS10
A0A2K3DB37_CHLRE__KS3
A0A2K3DB37_CHLRE__KS7
A0A2K3DB37_CHLRE__KS5
A0A2K3DB37_CHLRE__KS4
A0A2K3DB37_CHLRE__KS8
A0A2K3DB37_CHLRE__KS2
A0A2K3DB37_CHLRE__KS9
A0A2K3DB37_CHLRE__KS1
A0A2K3DB37_CHLRE__KS14
A0A2K3DB37_CHLRE__KS12
R1FUA6_EMIHU__KS2
R1DYV9_EMIHU__KS6
R1BNS0_EMIHU__KS12
R1BNS0_EMIHU__KS17
R1BNS0_EMIHU__KS18
R1BNS0_EMIHU__KS15
R1BNS0_EMIHU__KS16
PKZILLA-1__KS17
PKZILLA-1__KS12
PKZILLA-1__KS8
PKZILLA-1__KS6
PKZILLA-1__KS4
PKZILLA-1__KS5
PKZILLA-1__KS32
PKZILLA-2__KS4
PKZILLA-1__KS23
PKZILLA-1__KS26
PKZILLA-2__KS16
PKZILLA-1__KS7
PKZILLA-1__KS10
PKZILLA-1__KS28
PKZILLA-1__KS22
PKZILLA-1__KS25
PKZILLA-2__KS3
PKZILLA-1__KS19
PKZILLA-1__KS14
PKZILLA-2__KS12
PKZILLA-1__KS34
PKZILLA-2__KS11
PKZILLA-2__KS6
PKZILLA-2__KS9
R1FUA6_EMIHU__KS1
R1DYV9_EMIHU__KS3
R1DYV9_EMIHU__KS2
R1DYV9_EMIHU__KS1
R1DYV9_EMIHU__KS5
R1DYV9_EMIHU__KS7
R1C8E5_EMIHU__KS3
R1C8E5_EMIHU__KS2
R1C8E5_EMIHU__KS4
R1C8E5_EMIHU__KS1
R1ER06_EMIHU__KS3
0.500271

**Fig. S16:** Phylogenetic analysis of PKZILLA ketosynthase (KS) domains with an expansive set of comparable PKS domain sequences from Uniprot representing bacteria, fungi, green algae, dinoflagellates, haptophytes, and human fatty acid synthase (FAS). See table S10 for key for taxonomic interpretation. Remainder of sequence ID is the uniprot ID and N-terminal to C-terminal ascending domain number. Source data, workflow, and resulting plots available on Zenodo (doi:10.5281/zenodo.10247217).

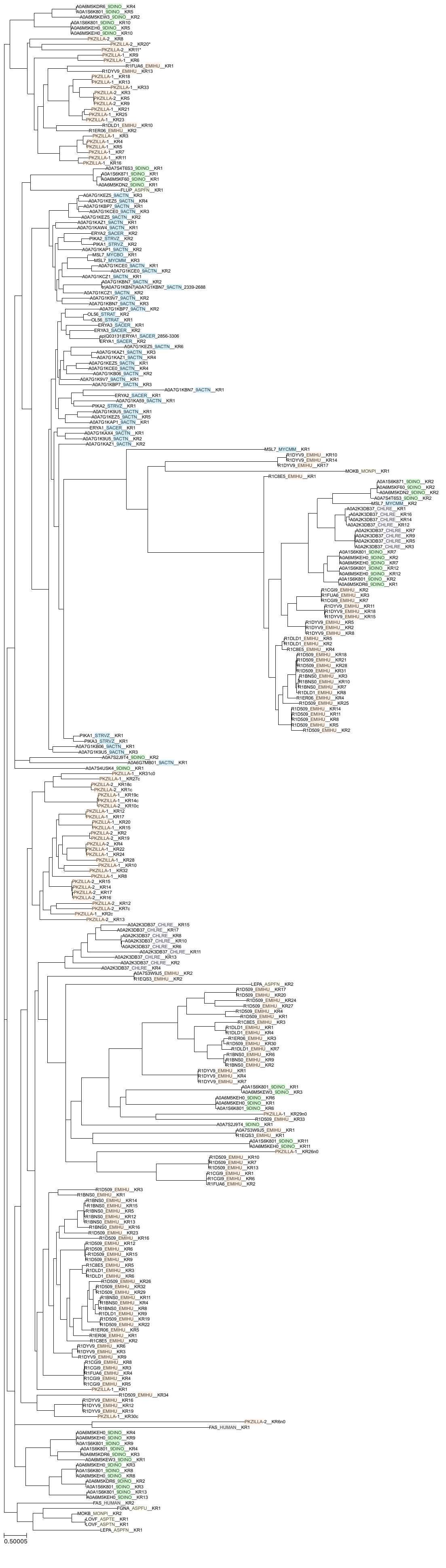
0.50005

**Fig. S17:** Phylogenetic analysis of PKZILLA ketoreductase (KR) domains with an expansive set of comparable PKS domain sequences from Uniprot representing bacteria, fungi, green algae, dinoflagellates, haptophytes, and human fatty acid synthase (FAS). See table S10 for key for taxonomic interpretation. Remainder of sequence ID is the uniprot ID and N-terminal to C-terminal ascending domain number. Source data, workflow, and resulting plots available on Zenodo (doi:10.5281/zenodo.10247217).

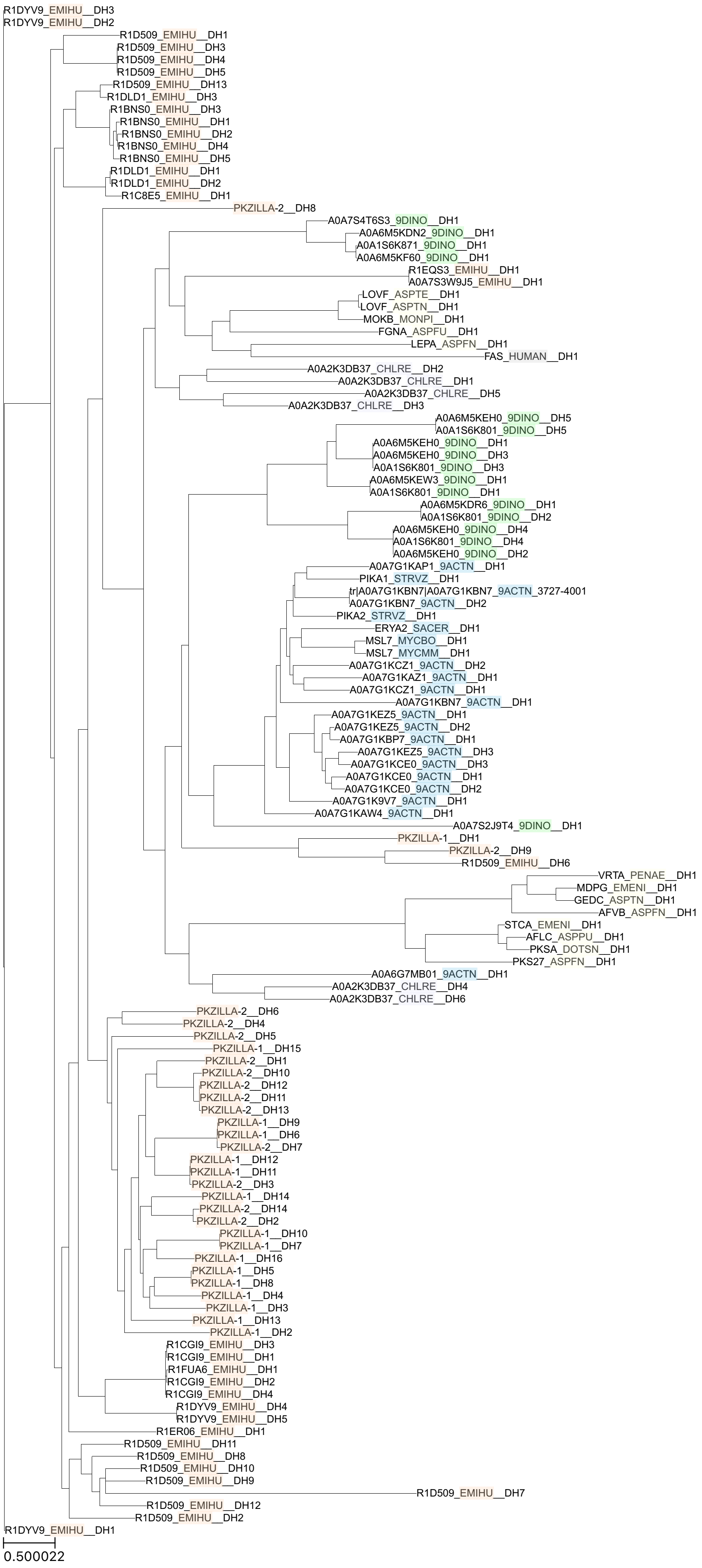
R1DYV9_EMIHU__DH3
R1DYV9_EMIHU__DH2
R1D509_EMIHU__DH1
R1D509_EMIHU__DH3
R1D509_EMIHU__DH4
R1D509_EMIHU__DH5
R1D509_EMIHU__DH13
R1DLD1_EMIHU__DH3
R1BNS0_EMIHU__DH3
R1BNS0_EMIHU__DH1
R1BNS0_EMIHU__DH2
R1BNS0_EMIHU__DH4
R1BNS0_EMIHU__DH5
R1DLD1_EMIHU__DH1
R1DLD1_EMIHU__DH2
R1C8E5_EMIHU__DH1
PKZILLA-2__DH8
A0A7S4T6S3_9DINO__DH1
A0A6M5KDN2_9DINO__DH1
A0A1S6K871_9DINO__DH1
A0A6M5KF60_9DINO__DH1
R1EQS3_EMIHU__DH1
A0A7S3W9J5_EMIHU__DH1
LOVF_ASPTE__DH1
LOVF_ASPTN__DH1
MOKB_MONPI__DH1
FGNA_ASPFU__DH1
LEPA_ASPFN__DH1
FAS_HUMAN__DH1
A0A2K3DB37_CHLRE__DH2
A0A2K3DB37_CHLRE__DH1
A0A2K3DB37_CHLRE__DH5
A0A2K3DB37_CHLRE__DH3
A0A6M5KEH0_9DINO__DH5
A0A1S6K801_9DINO__DH5
A0A6M5KEH0_9DINO__DH1
A0A6M5KEH0_9DINO__DH3
A0A1S6K801_9DINO__DH3
A0A6M5KEW3_9DINO__DH1
A0A1S6K801_9DINO__DH1
A0A6M5KDR6_9DINO__DH1
A0A1S6K801_9DINO__DH2
A0A6M5KEH0_9DINO__DH4
A0A1S6K801_9DINO__DH4
A0A6M5KEH0_9DINO__DH2
A0A7G1KAP1_9ACTN__DH1
PIKA1_STRVZ__DH1
tr|A0A7G1KBN7|A0A7G1KBN7_9ACTN_3727-4001
A0A7G1KBN7_9ACTN__DH2
PIKA2_STRVZ__DH1
ERYA2_SACER__DH1
MSL7_MYCBO__DH1
MSL7_MYCMM__DH1
A0A7G1KCZ1_9ACTN__DH2
A0A7G1KAZ1_9ACTN__DH1
A0A7G1KCZ1_9ACTN__DH1
A0A7G1KBN7_9ACTN__DH1
A0A7G1KEZ5_9ACTN__DH1
A0A7G1KEZ5_9ACTN__DH2
A0A7G1KBP7_9ACTN__DH1
A0A7G1KEZ5_9ACTN__DH3
A0A7G1KCE0_9ACTN__DH3
A0A7G1KCE0_9ACTN__DH1
A0A7G1KCE0_9ACTN__DH2
A0A7G1K9V7_9ACTN__DH1
A0A7G1KAW4_9ACTN__DH1
A0A7S2J9T4_9DINO__DH1
PKZILLA-1__DH1
PKZILLA-2__DH9
R1D509_EMIHU__DH6
VRTA_PENAE__DH1
MDPG_EMENI__DH1
GEDC_ASPTN__DH1
AFVB_ASPFN__DH1
STCA_EMENI__DH1
AFLC_ASPPU__DH1
PKSA_DOTSN__DH1
PKS27_ASPFN__DH1
A0A6G7MB01_9ACTN__DH1
A0A2K3DB37_CHLRE__DH4
A0A2K3DB37_CHLRE__DH6
PKZILLA-2__DH6
PKZILLA-2__DH4
PKZILLA-2__DH5
PKZILLA-1__DH15
PKZILLA-2__DH1
PKZILLA-2__DH10
PKZILLA-2__DH12
PKZILLA-2__DH11
PKZILLA-2__DH13
PKZILLA-1__DH9
PKZILLA-1__DH6
PKZILLA-2__DH7
PKZILLA-1__DH12
PKZILLA-1__DH11
PKZILLA-2__DH3
PKZILLA-1__DH14
PKZILLA-2__DH14
PKZILLA-2__DH2
PKZILLA-1__DH10
PKZILLA-1__DH7
PKZILLA-1__DH16
PKZILLA-1__DH5
PKZILLA-1__DH8
PKZILLA-1__DH4
PKZILLA-1__DH3
PKZILLA-1__DH13
PKZILLA-1__DH2
R1CGI9_EMIHU__DH3
R1CGI9_EMIHU__DH1
R1FUA6_EMIHU__DH1
R1CGI9_EMIHU__DH2
R1CGI9_EMIHU__DH4
R1DYV9_EMIHU__DH4
R1DYV9_EMIHU__DH5
R1ER06_EMIHU__DH1
R1D509_EMIHU__DH11
R1D509_EMIHU__DH8
R1D509_EMIHU__DH10
R1D509_EMIHU__DH9
R1D509_EMIHU__DH7
R1D509_EMIHU__DH12
R1D509_EMIHU__DH2
R1DYV9_EMIHU__DH1
0.500022

**Fig. S18:** Phylogenetic analysis of PKZILLA dehydratase (DH) domains with an expansive set of comparable PKS domain sequences from Uniprot representing bacteria, fungi, green algae, dinoflagellates, haptophytes, and human fatty acid synthase (FAS). See table S10 for key for taxonomic interpretation. Remainder of sequence ID is the uniprot ID and N-terminal to C-terminal ascending domain number. Source data, workflow, and resulting plots available on Zenodo (doi:10.5281/zenodo.10247217).

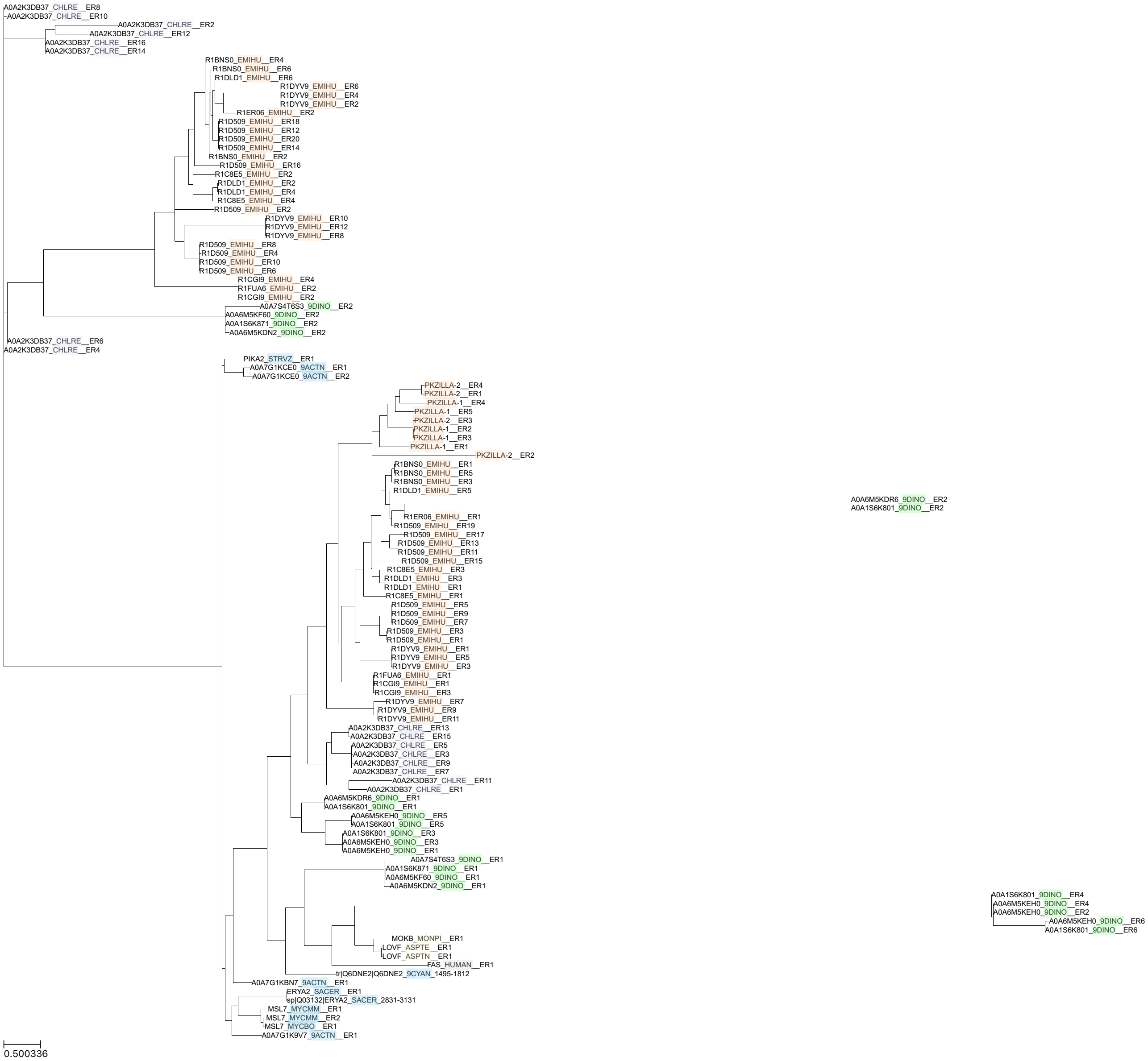


**Fig. S19:** Phylogenetic analysis of PKZILLA enoyl reductase (ER) domains with an expansive set of comparable PKS domain sequences sequences from Uniprot representing bacteria, fungi, green algae, dinoflagellates, haptophytes, and human fatty acid synthase (FAS). See table S10 for key for taxonomic interpretation. Remainder of sequence ID is the uniprot ID and N-terminal to C-terminal ascending domain number. Source data, workflow, and resulting plots available on Zenodo (doi:10.5281/zenodo.10247217).

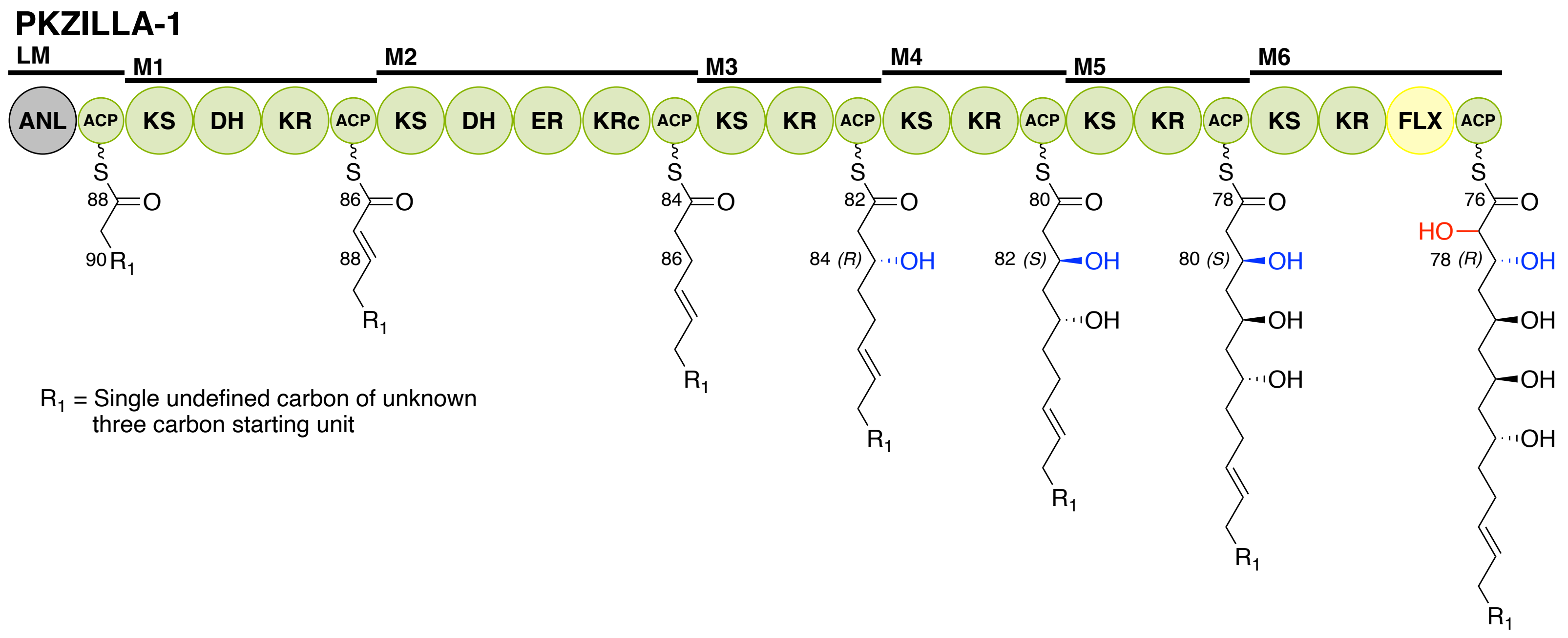
PKZILLA-1
LM
M1
M2
M3
M4
M5
M6
ANL
ACP
KS
DH
KR
ER
KRc
FLX
R1 = Single undefined carbon of unknown three carbon starting unit

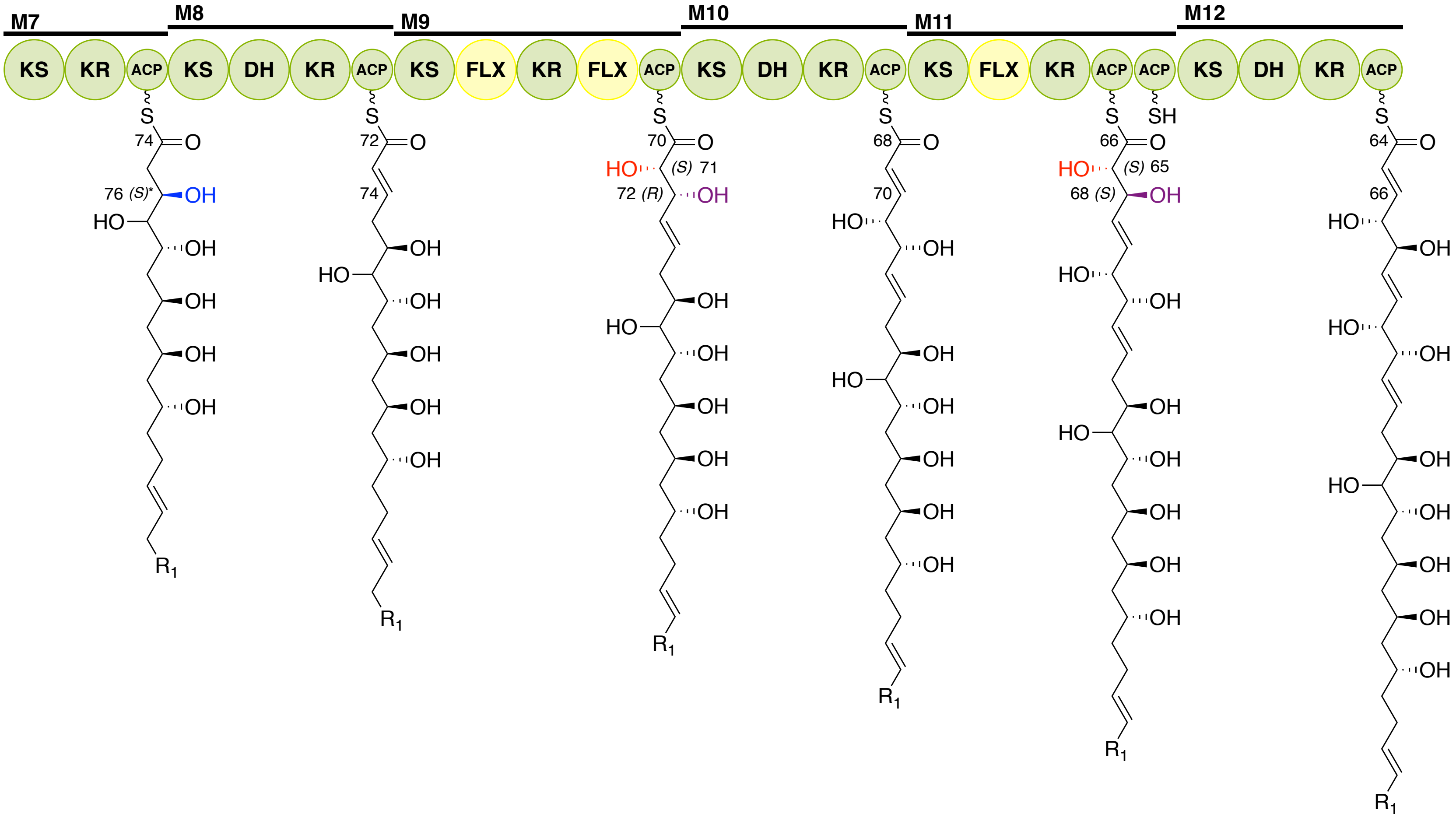
M7
M8
M9
M10
M11
M12
KS
KR
ACP
DH
FLX
SH
R1

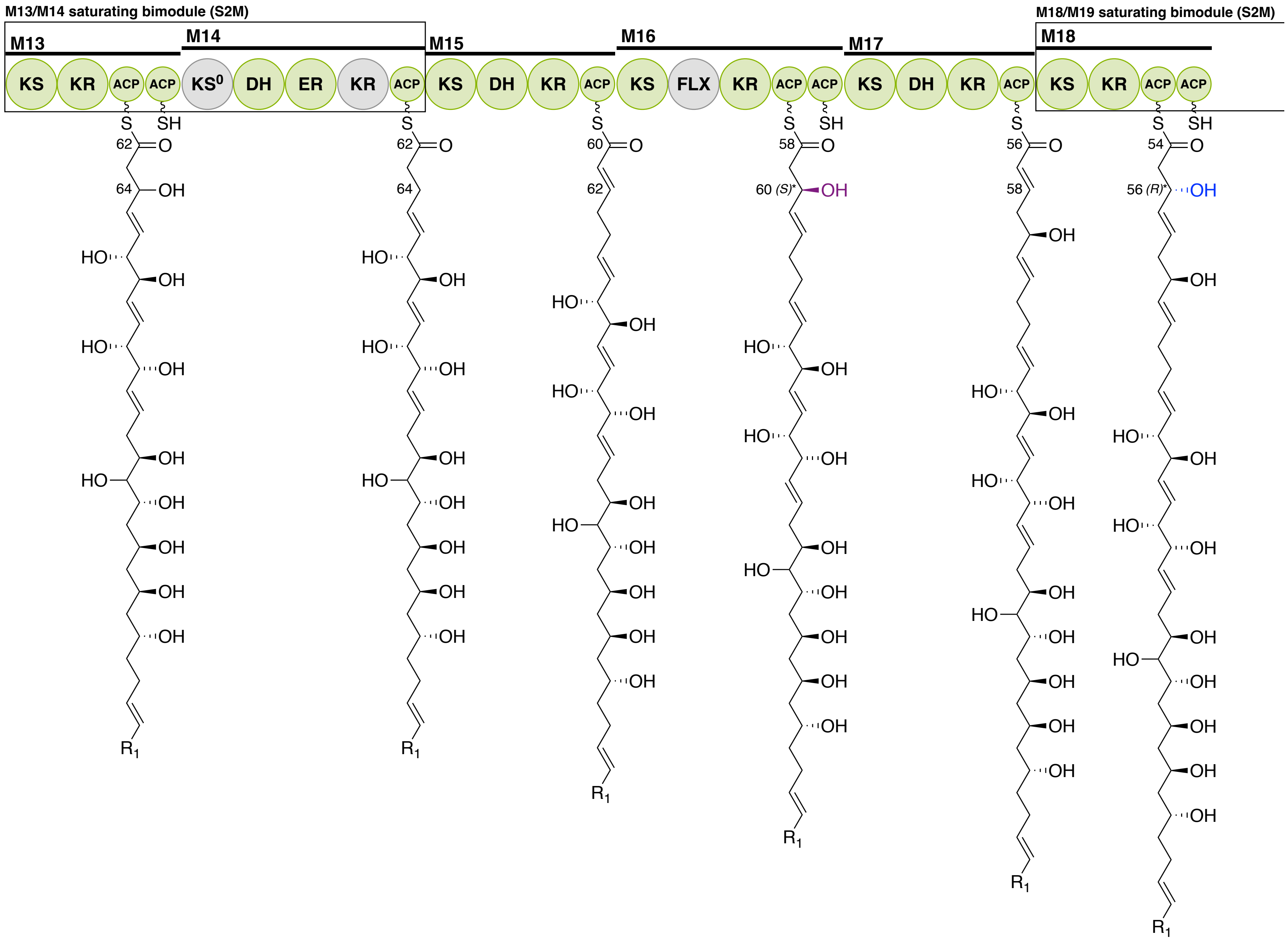

M13/M14 saturating bimodule (S2M)
M18/M19 saturating bimodule (S2M)
M13
M14
M15
M16
M17
M18
KS
KR
ACP
KS0
DH
ER
FLX
S
SH
OH
HO
R1
60 (S)*
56 (R)*

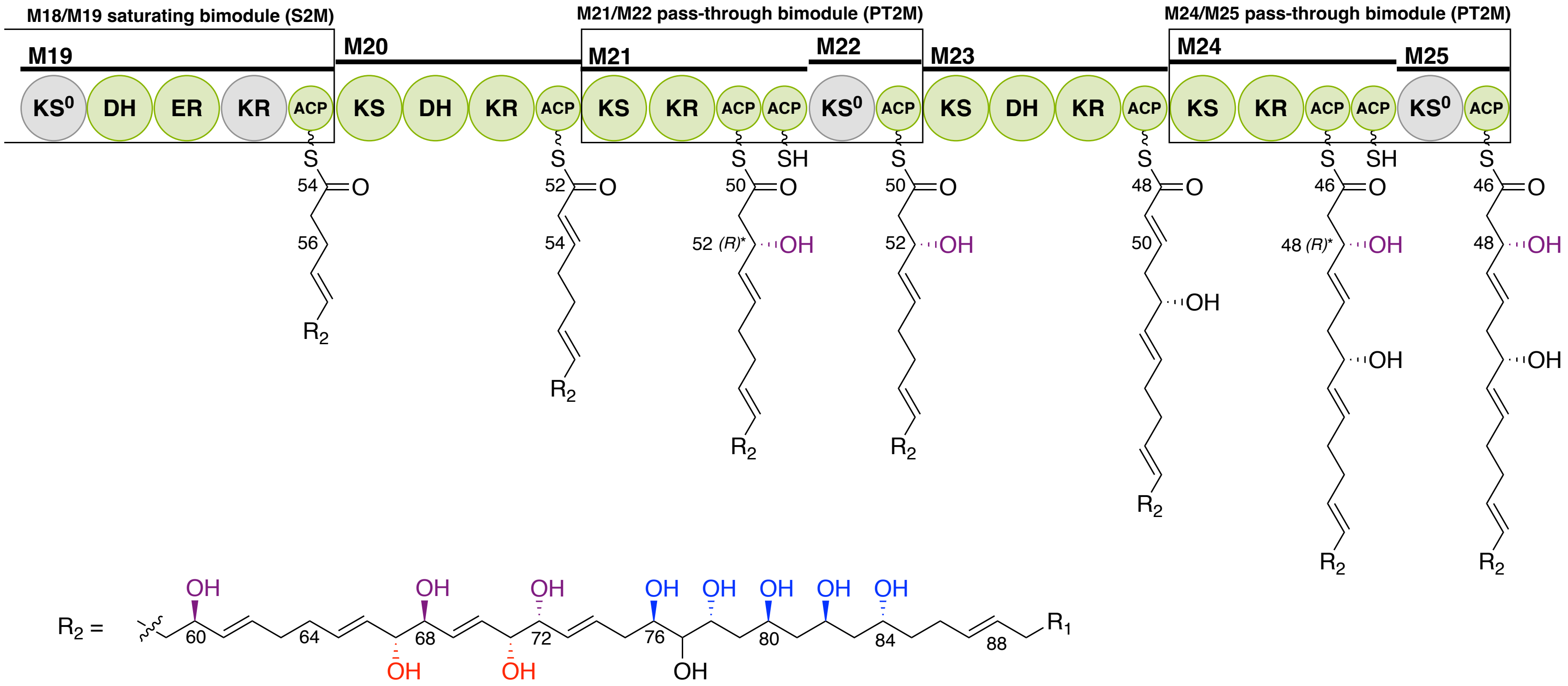

M18/M19 saturating bimodule (S2M)
M21/M22 pass-through bimodule (PT2M)
M24/M25 pass-through bimodule (PT2M)
M19
M20
M21
M22
M23
M24
M25
KS0 DH ER KR ACP KS DH KR ACP KS KR ACP ACP KS0 ACP KS DH KR ACP KS KR ACP ACP KS0 ACP
S
SH
54
56
52
54
50
52 (R)*
OH
48
48 (R)*
46
R2
R2 =
60
64
68
72
76
80
84
88
R1

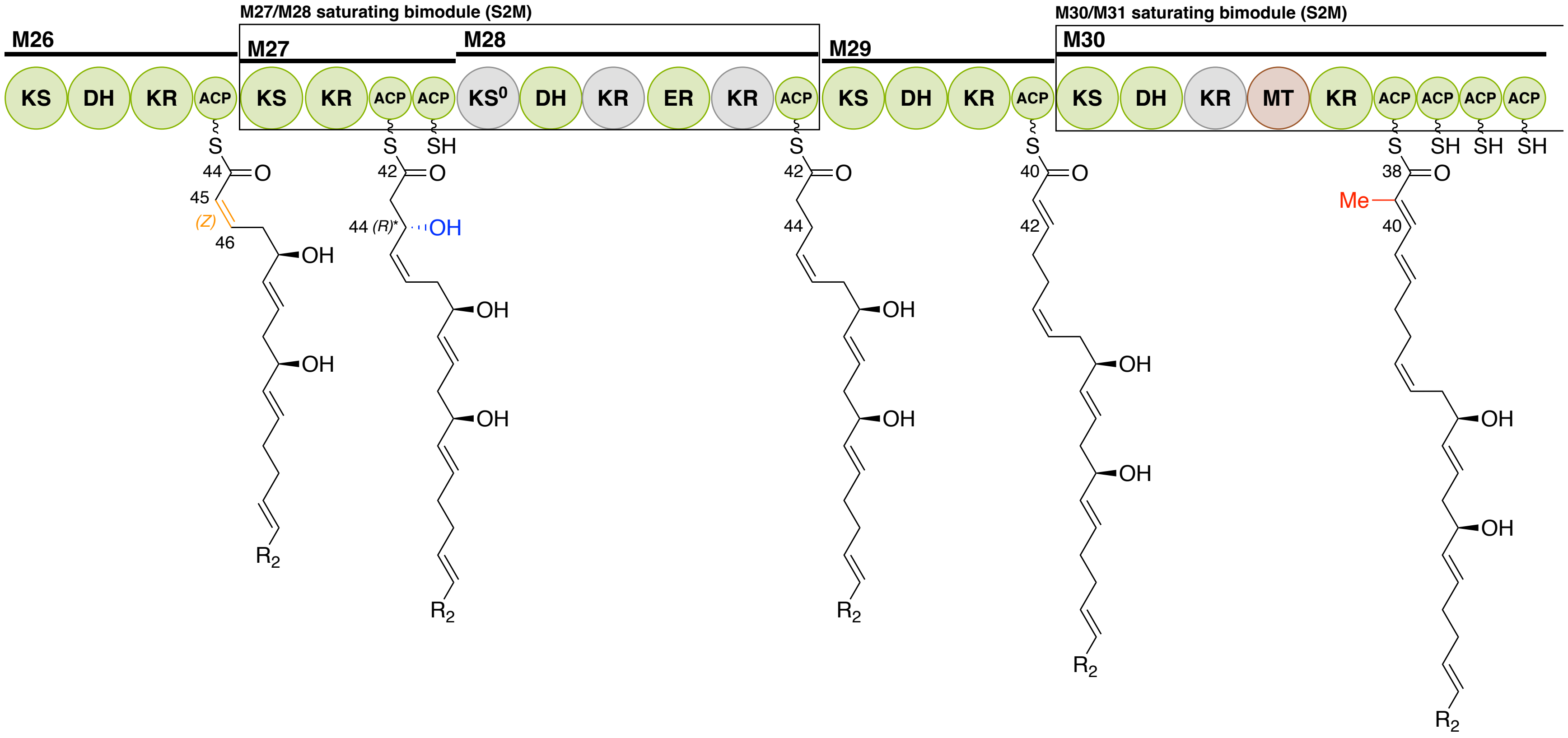
M27/M28 saturating bimodule (S2M)
M30/M31 saturating bimodule (S2M)
M26
M27
M28
M29
M30
KS
DH
KR
ACP
KS⁰
ER
MT
S
SH
O
OH
Me
(Z)
(R)*
R2
44
45
46
42
40
38

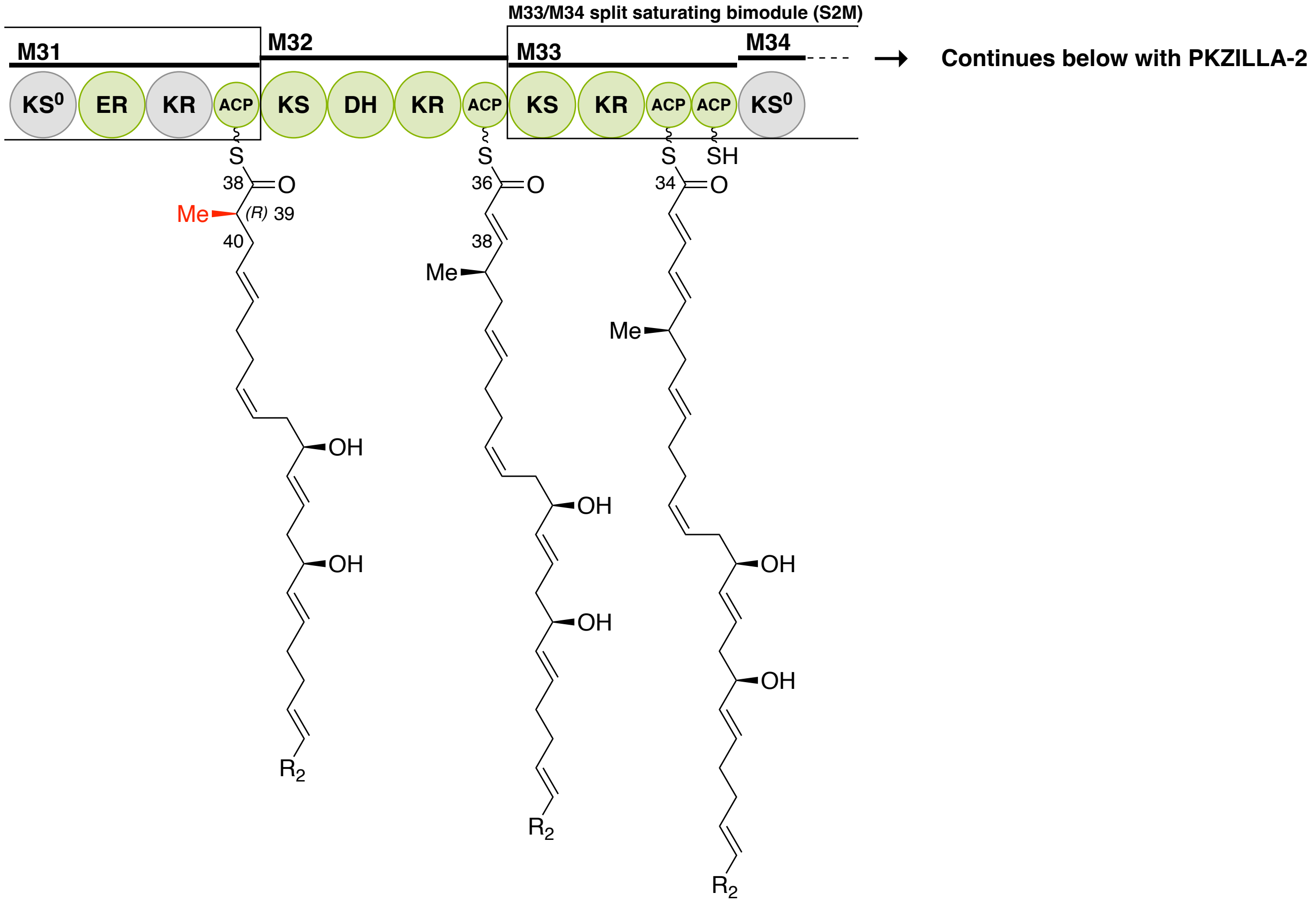
M33/M34 split saturating bimodule (S2M)
M31
M32
M33
M34
KS0
ER
KR
ACP
KS
DH
KR
ACP
KS
KR
ACP
ACP
KS0
S
S
S
SH
38
O
Me
(R) 39
40
36
38
34
Me
Me
OH
OH
R2
Continues below with PKZILLA-2

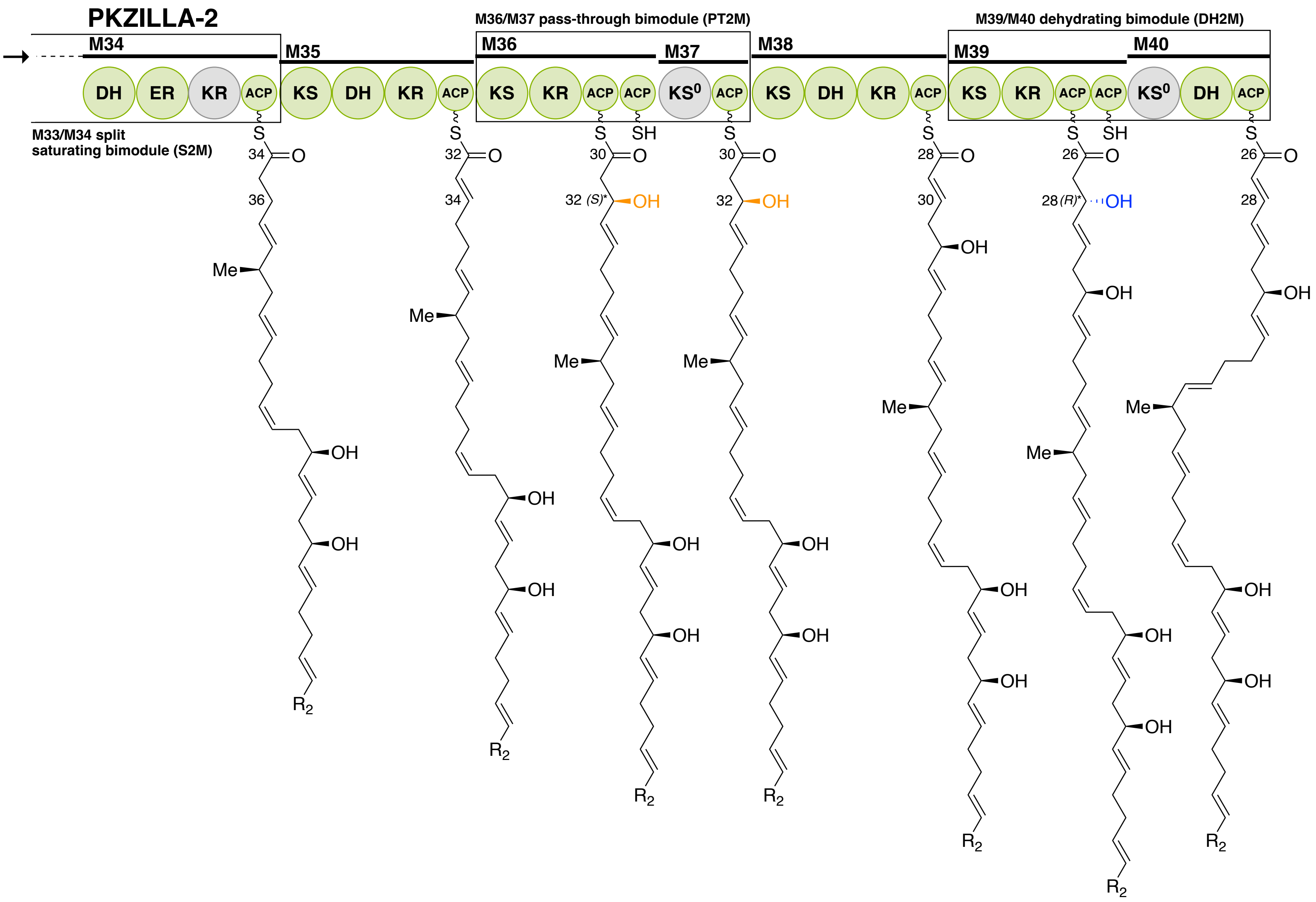

PKZILLA-2
M34
M35
M36/M37 pass-through bimodule (PT2M)
M36
M37
M38
M39/M40 dehydrating bimodule (DH2M)
M39
M40
DH
ER
KR
ACP
KS
DH
KR
ACP
KS
KR
ACP
ACP
KS0
ACP
KS
DH
KR
ACP
KS
KR
ACP
ACP
KS0
DH
ACP
M33/M34 split
saturating bimodule (S2M)
32 (S)*
28 (R)*

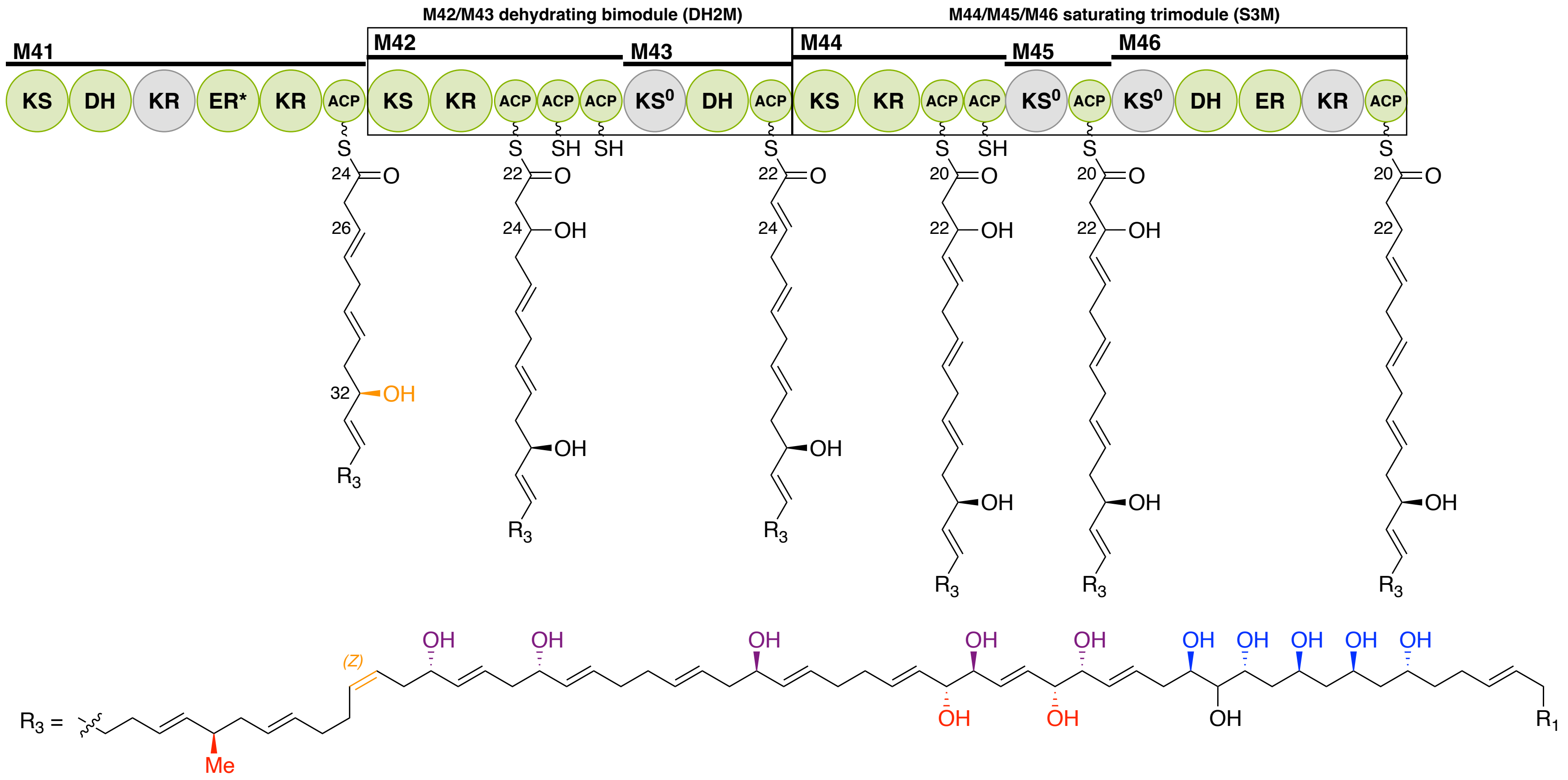

M42/M43 dehydrating bimodule (DH2M)
M44/M45/M46 saturating trimodule (S3M)
M41
M42
M43
M44
M45
M46
KS DH KR ER* KR ACP KS KR ACP ACP ACP KS0 DH ACP KS KR ACP ACP KS0 ACP KS0 DH ER KR ACP
R3 =
Me
(Z)
R1

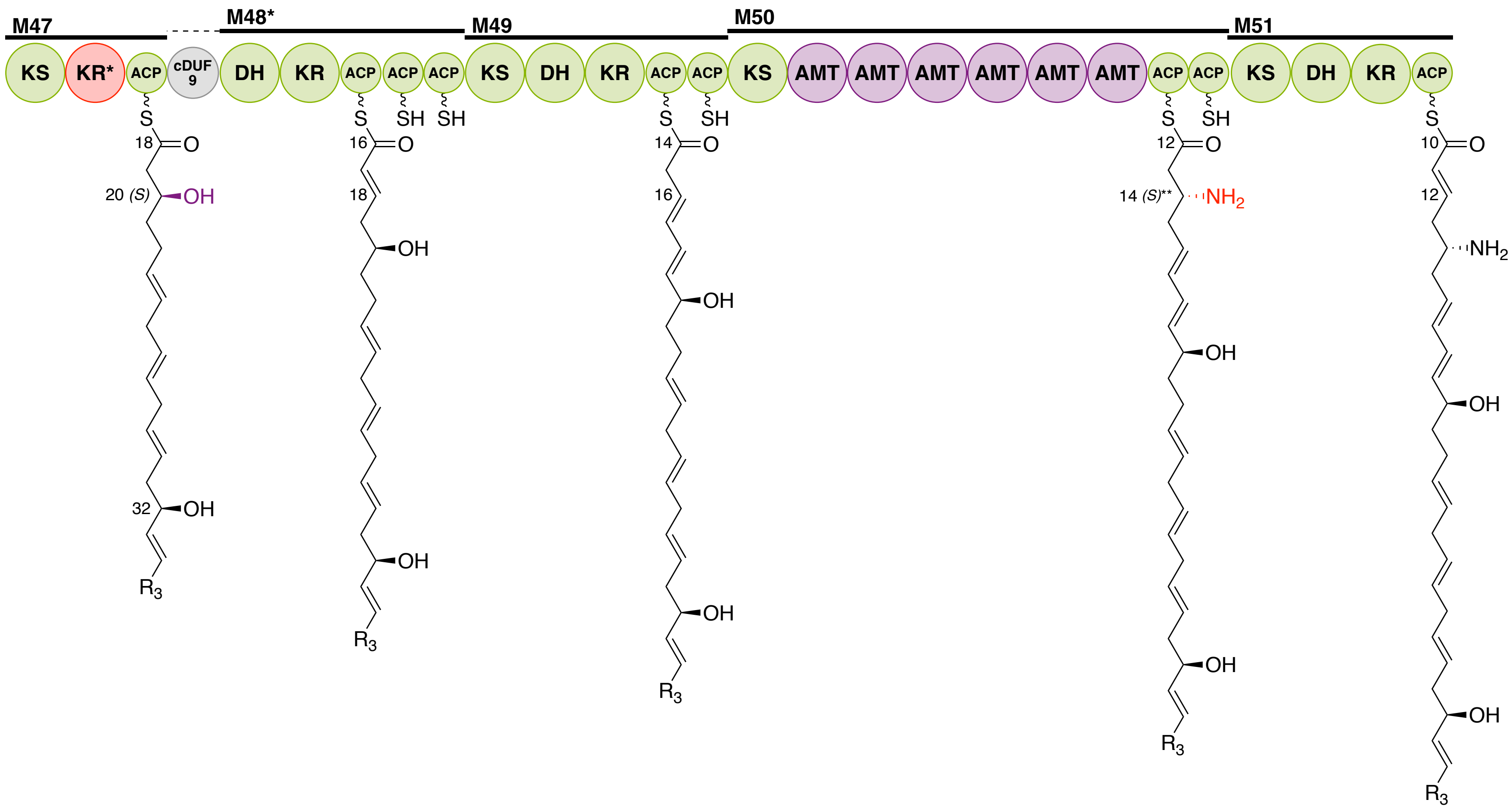
M47
M48*
M49
M50
M51
KS
KR*
ACP
cDUF 9
DH
KR
ACP
ACP
ACP
KS
DH
KR
ACP
ACP
KS
AMT
AMT
AMT
AMT
AMT
AMT
ACP
ACP
KS
DH
KR
ACP
18
20 (S)
OH
32
OH
R3
16
18
OH
OH
R3
14
16
OH
OH
R3
12
14 (S)**
NH2
OH
OH
R3
10
12
NH2
OH
OH
R3

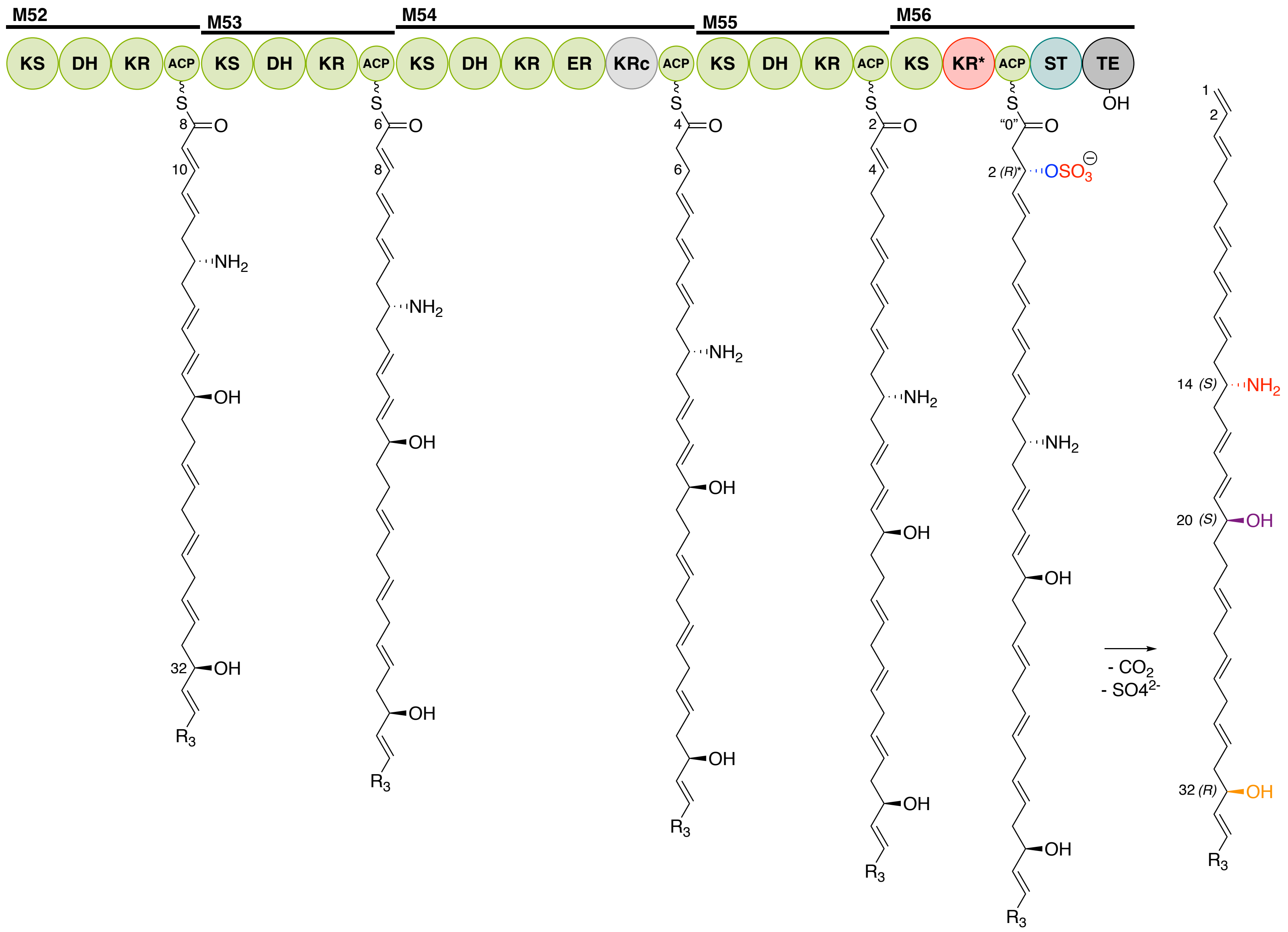

M52
M53
M54
M55
M56
KS
DH
KR
ACP
ER
KRc
KR*
ST
TE
OH
S
O
NH2
OH
R3
8
10
32
6
4
2
"0"
2 (R)*
OSO3
14 (S)
20 (S)
32 (R)
1
- CO2
- SO4 2-

**Fig. S20:** Polyketide synthase assembly line model for PKZILLA-1/2. Domain abbreviations and color coding follows those used in Fig. 2, excepting that all KS, ACP, KR, DH, ER domains are shown as green. Light-gray colored domains are either predicted to be catalytically inactive from bioinformatic analysis (KRs), or trans-acylating rather than fully condensating ($KS^0$s), or did not have an apparent attributable catalytic activity (i.e. M16 FLX) despite bioinformatic expectations of catalytic capability. Red domains (KR*s) in M47 and M56, would be expected to be active based on the biosynthetic model, but are bioinformatically predicted to be inactive (table S9), and thus may be false positive predictions of inactivity caused by divergent catalytic properties. Red bond and atom coloration indicates functional groups installed by FLX, MT, AMT, and ST domains, or installed functional groups whose stereochemistry is known only from inferences from empirically reported or retrobiosynthetically inferred stereochemistry (Fig. 3) and does not have a bioinformatic prediction of stereochemistry (Table S12). Blue coloration indicates stereochemistry is only proposed from bioinformatics (Table S12). Purple coloration indicates a match between empirically reported or inferred stereochemical configurations and bioinformatic stereochemical predictions (Table S12). Orange coloration indicates a mis-match between empirically reported or inferred stereochemical/isomeric configurations and bioinformatic predictions (Table S12). *=Due to inversion of Cahn–Ingold–Prelog priority when γ,δ bond is an alkene vs saturated, or if the γ carbon has a hydroxyl, γ,δ carbons were drawn as always saturated and without hydroxyls for the purposes of stereochemical assignment. **=Stereochemical assignment inverted to match configuration on prymnesin, in both cases it is the same configuration barring an inversion of Cahn–Ingold–Prelog priority.

| Transcript ID | Gene ID | Scaffold ID | Gene coord. start | Gene coord. end | Gene length (bp) | mRNA length (nt) | Exon count | Closest PKS gene ID | Closest PKS gene distance (kbp) | tpm_day | tpm_night |
|---|---|---|---|---|---|---|---|---|---|---|---|
| PKZILLA-1 | 12B1gPKZILLA-1 | 12B1-Scaf17 | 2286087 | 2423367 | 137280 | 136088 | 17 | | | 1.11135 | 0.713157 |
| PKZILLA-2 | 12B1gPKZILLA-2 | 12B1-Scaf7 | 1023621 | 1116873 | 93252 | 92413 | 12 | 12B1g21318 | -680.4 | 2.42293 | 1.67459 |
| PKZILLA-3 | 12B1gPKZILLA-3 | 12B1-Scaf10 | 233715 | 308247 | 74532 | 73850 | 8 | | | 0.0327181 | 0.907688 |
| 12B1g6935.t1 | 12B1g6935 | 12B1-Scaf30 | 1090474 | 1117079 | 26605 | 24501 | 7 | | | 0.0122754 | 0.102986 |
| 12B1g20558.t1 | 12B1g20558 | 12B1-Scaf24 | 1258375 | 1280190 | 21815 | 19167 | 25 | | | 13.5373 | 5.5826 |
| 12B1g8858.t2 | 12B1g8858 | 12B1-Scaf14 | 381850 | 402639 | 20789 | 20367 | 6 | 12B1g8877 | 90.7 | 0.297101 | 0.496385 |
| 12B1g8858.t1 | 12B1g8858 | 12B1-Scaf14 | 381850 | 402639 | 20789 | 20448 | 5 | 12B1g8877 | 90.7 | 0.00288062 | 0.0022481 |
| 12B1g21318.t2 | 12B1g21318 | 12B1-Scaf7 | 323056 | 343225 | 20169 | 19878 | 2 | 12B1gPKZILLA-2 | 680.4 | 0.306636 | 0.169165 |
| 12B1g21318.t1 | 12B1g21318 | 12B1-Scaf7 | 323056 | 343225 | 20169 | 3753 | 3 | 12B1gPKZILLA-2 | 680.4 | 0 | 0 |
| 12B1g8877.t1 | 12B1g8877 | 12B1-Scaf14 | 493291 | 508389 | 15098 | 13578 | 7 | 12B1g8858 | -90.7 | 1.90592 | 1.52636 |
| 12B1g521.t1 | 12B1g521 | 12B1-Scaf9 | 1906255 | 1920566 | 14311 | 11718 | 19 | | | 6.72458 | 8.85982 |
| 12B1g5199.t1 | 12B1g5199 | 12B1-Scaf22 | 974085 | 988095 | 14010 | 13545 | 3 | | | 0.359611 | 0.358954 |
| 12B1g8961.t1 | 12B1g8961 | 12B1-Scaf14 | 799146 | 811847 | 12701 | 11613 | 9 | 12B1g8877 | -290.8 | 0.12426 | 0.542579 |
| 12B1g16712.t1 | 12B1g16712 | 12B1-Scaf1 | 2251945 | 2264026 | 12081 | 11883 | 2 | | | 1.17317 | 0.828558 |
| 12B1g5573.t1 | 12B1g5573 | 12B1-Scaf21 | 668309 | 680198 | 11889 | 11493 | 6 | | | 1.2576 | 1.35399 |
| 12B1g22763.t1 | 12B1g22763 | 12B1-Scaf16 | 1956638 | 1967275 | 10637 | 7761 | 12 | | | 11.6529 | 5.12936 |
| 12B1g3947.t1 | 12B1g3947 | 12B1-Scaf5 | 908335 | 918651 | 10316 | 6363 | 7 | 12B1g3948 | 0.5 | 0.751993 | 1.59325 |
| 12B1g4698.t1 | 12B1g4698 | 12B1-Scaf5 | 3795505 | 3804614 | 9109 | 8130 | 5 | 12B1g4700 | 3 | 2.1088 | 1.27798 |
| 12B1g14437.t1 | 12B1g14437 | 12B1-Scaf3 | 1062558 | 1071078 | 8520 | 8127 | 4 | | | 0.725751 | 0.42423 |
| 12B1g3948.t1 | 12B1g3948 | 12B1-Scaf5 | 919101 | 927118 | 8017 | 5415 | 10 | 12B1g3947 | -0.5 | 0.37918 | 1.12212 |
| 12B1g23642.t1 | 12B1g23642 | 12B1-Scaf26 | 980919 | 987804 | 6885 | 6585 | 4 | | | 15.6463 | 9.60298 |
| 12B1g4700.t1 | 12B1g4700 | 12B1-Scaf5 | 3807648 | 3814260 | 6612 | 4836 | 6 | 12B1g4698 | -3 | 12.1958 | 3.27771 |
| 12B1g8154.t1 | 12B1g8154 | 12B1-Scaf13 | 326108 | 332557 | 6449 | 6225 | 3 | | | 1.38845 | 0.879017 |
| 12B1g18060.t1 | 12B1g18060 | 12B1-Scaf33 | 408311 | 411415 | 3105 | 3105 | 1 | | | 5.65643 | 2.4474 |

**Table S1:** Genomic characteristics of *Prymnesium parvum* 12B1 PKS genes. coord.=coordinate. tpm_day and tpm_night = expression values in transcripts per million (TPM) for day and night rRNA depletion RNA-seq libraries, calculated with Kallisto (doi:10.5281/zenodo.10023425).

| seqid | scaffold | start | end | intron length | Upstream reading frame | Downstream reading frame | Reading frame shift from splicing? | Stop codon in intron & upstream reading frame? |
|---|---|---|---|---|---|---|---|---|
| PKZILLA-1_intron1 | 12B1-Scaf17 | 2287899 | 2287961 | 62 | 3 | 3 | no | yes |
| PKZILLA-1_intron2 | 12B1-Scaf17 | 2291865 | 2291950 | 85 | 3 | 2 | yes | no |
| PKZILLA-1_intron3 | 12B1-Scaf17 | 2296229 | 2296296 | 67 | 2 | 1 | yes | no |
| PKZILLA-1_intron4 | 12B1-Scaf17 | 2317507 | 2317572 | 65 | 1 | 1 | no | yes |
| PKZILLA-1_intron5 | 12B1-Scaf17 | 2327560 | 2327633 | 73 | 1 | 3 | yes | yes |
| PKZILLA-1_intron6 | 12B1-Scaf17 | 2336046 | 2336111 | 65 | 3 | 3 | no | yes |
| PKZILLA-1_intron7 | 12B1-Scaf17 | 2343831 | 2343904 | 73 | 3 | 2 | yes | no |
| PKZILLA-1_intron8 | 12B1-Scaf17 | 2348525 | 2348601 | 76 | 2 | 1 | yes | yes |
| PKZILLA-1_intron9 | 12B1-Scaf17 | 2357062 | 2357127 | 65 | 1 | 1 | no | yes |
| PKZILLA-1_intron10 | 12B1-Scaf17 | 2364847 | 2364920 | 73 | 1 | 3 | yes | no |
| PKZILLA-1_intron11 | 12B1-Scaf17 | 2369541 | 2369617 | 76 | 3 | 2 | yes | yes |
| PKZILLA-1_intron12 | 12B1-Scaf17 | 2379032 | 2379132 | 100 | 2 | 1 | yes | yes |
| PKZILLA-1_intron13 | 12B1-Scaf17 | 2388484 | 2388584 | 100 | 1 | 3 | yes | yes |
| PKZILLA-1_intron14 | 12B1-Scaf17 | 2396328 | 2396394 | 66 | 3 | 1 | yes | no |
| PKZILLA-1_intron15 | 12B1-Scaf17 | 2401024 | 2401091 | 67 | 1 | 3 | yes | yes |
| PKZILLA-1_intron16 | 12B1-Scaf17 | 2415990 | 2416054 | 64 | 3 | 2 | yes | no |
| PKZILLA-2_intron1 | 12B1-Scaf7 | 1024581 | 1024646 | 65 | 3 | 3 | no | no |
| PKZILLA-2_intron2 | 12B1-Scaf7 | 1029285 | 1029356 | 71 | 3 | 3 | no | yes |
| PKZILLA-2_intron3 | 12B1-Scaf7 | 1038708 | 1038808 | 100 | 3 | 2 | yes | yes |
| PKZILLA-2_intron4 | 12B1-Scaf7 | 1065686 | 1065759 | 73 | 2 | 1 | yes | yes |
| PKZILLA-2_intron5 | 12B1-Scaf7 | 1072585 | 1072709 | 124 | 1 | 3 | yes | yes |
| PKZILLA-2_intron6 | 12B1-Scaf7 | 1092303 | 1092366 | 63 | 3 | 1 | yes | yes |
| PKZILLA-2_intron7 | 12B1-Scaf7 | 1096189 | 1096253 | 64 | 1 | 3 | yes | no |
| PKZILLA-2_intron8 | 12B1-Scaf7 | 1100079 | 1100143 | 64 | 3 | 2 | yes | no |
| PKZILLA-2_intron9 | 12B1-Scaf7 | 1103969 | 1104033 | 64 | 2 | 1 | yes | no |
| PKZILLA-2_intron10 | 12B1-Scaf7 | 1110112 | 1110183 | 71 | 1 | 1 | no | yes |
| PKZILLA-2_intron11 | 12B1-Scaf7 | 1115472 | 1115542 | 70 | 1 | 3 | yes | no |
| PKZILLA-3_intron1 | 12B1-Scaf10 | 234816 | 234888 | 72 | 3 | 1 | yes | yes |
| PKZILLA-3_intron2 | 12B1-Scaf10 | 234974 | 235031 | 57 | 1 | 2 | yes | yes |
| PKZILLA-3_intron3 | 12B1-Scaf10 | 235152 | 235267 | 115 | 2 | 1 | yes | yes |
| PKZILLA-3_intron4 | 12B1-Scaf10 | 235849 | 235976 | 127 | 1 | 3 | yes | yes |
| PKZILLA-3_intron5 | 12B1-Scaf10 | 237153 | 237288 | 135 | 3 | 1 | yes | no |
| PKZILLA-3_intron6 | 12B1-Scaf10 | 257254 | 257316 | 62 | 1 | 1 | no | no |
| PKZILLA-3_intron7 | 12B1-Scaf10 | 303779 | 303887 | 108 | 1 | 2 | yes | yes |

**Table S2:** Intron characteristics of PKZILLA genes. See Fig. S3, S4, S5 for detailed graphical views of each intron.

| InterProScan annotated region | Short PKS nomenclature | Comments |
|---|---|---|
| SSF56801 | ANL | Unintegrated into InterPro, but, covers both the "N" and "C" terminal subdomains of an ANL enzyme, unlike other matchers. Alternatively, IPR042099 matches the N-terminal (sub)domain, IPR025110 matches the C-terminal (sub)domain. |
| G3DSA:1.10.1200.10 | ACP | Integrated into InterPro:IPR036736 |
| G3DSA:3.40.47.10 | KS | Integrated into InterPro:IPR016039 |
| G3DSA:3.10.129.110 | DH | Integrated into InterPro:IPR042104 |
| G3DSA:3.40.50.720 | KR | Unintegrated into InterPro. Confirmed cross-match: Ends up nested underneath G3DSA:3.90.180.10 (ER). But, the longest available matcher for the KR. |
| G3DSA:3.90.180.10 | ER | Unintegrated into InterPro. Possible cross-match: Ends up nested underneath G3DSA:3.40.50.720 (KR)?. But, the longest available matcher for the ER. IPR020843 is a possible alternative. |
| G3DSA:3.50.50.60 | FLX | Integrated into InterPro:IPR036188 |
| G3DSA:3.40.50.150 | MT | Integrated into InterPro:IPR029063 |
| G3DSA:3.90.1150.10 | AMT | Integrated into InterPro:IPR015422 |
| SSF53474 | TE | Integrated into InterPro:IPR029058 |
| PF13469 | ST | Integrated into InterPro:IPR027417. Formerly annotated with G3DSA:3.40.50.300, but PF13469/sulfotransferase is a more informative annotation. |
| PF00668 | C | Integrated into InterPro:IPR001242. NRPS-like condensation domain. In PKZILLA-3 |
| G3DSA:3.30.1610.10 | S59 | Integrated into InterPro:IPR036903. This is a weak hit on PKZILLA-3. Autoproteolytic C-terminal nucleoporin thing. |

**Table S3:** Specific InterProScan annotations that were deemed to be diagnostic for and representative of particular PKS domains within the PKZILLA polypeptides, and used to select these domains for downstream analyses and figures. The table includes all non-trivially small and/or weak matches and non-intrinsically disordered region matches that were found in the PKZLLAs. Table is derived from “run2_filter.tsv” used in Zenodo workflows for InterProScan annotations of the PKZILLAs (doi:10.5281/zenodo.10023460), and non-PKZILLAs (doi:10.5281/zenodo.10011739).

| seqid | domain sequence |
|---|---|
| PKZILLA-1 | ANL-ACP-KS-DH-KR-ACP-KS-DH-ER-KR-ACP-KS-KR-ACP-KS-KR-ACP-KS-KR-ACP-KS-KR-FLX-ACP-KS-KR-ACP-KS-DH-KR-ACP-KS-FLX-KR-FLX-ACP-KS-DH-KR-ACP-KS-FLX-KR-ACP-ACP-KS-DH-KR-ACP-KS-KR-ACP-ACP-KS-DH-ER-KR-ACP-KS-DH-KR-ACP-KS-FLX-KR-ACP-ACP-KS-DH-KR-ACP-KS-KR-ACP-ACP-KS-DH-ER-KR-ACP-KS-DH-KR-ACP-KS-KR-ACP-ACP-KS-ACP-KS-DH-KR-ACP-KS-KR-ACP-ACP-KS-ACP-KS-DH-KR-ACP-KS-KR-ACP-ACP-KS-DH-KR-ER-KR-ACP-KS-DH-KR-ACP-KS-DH-KR-MT-KR-ACP-ACP-ACP-ACP-KS-ER-KR-ACP-KS-DH-KR-ACP-KS-KR-ACP-ACP-KS |
| PKZILLA-2 | DH-ER-KR-ACP-KS-DH-KR-ACP-KS-KR-ACP-ACP-KS-ACP-KS-DH-KR-ACP-KS-KR-ACP-ACP-KS-DH-ACP-KS-DH-KR-ER-KR-ACP-KS-KR-ACP-ACP-ACP-KS-DH-ACP-KS-KR-ACP-ACP-KS-ACP-KS-DH-ER-KR-ACP-KS-KR-ACP-DH-KR-ACP-ACP-ACP-KS-DH-KR-ACP-ACP-KS-AMT-AMT-AMT-AMT-AMT-AMT-ACP-ACP-KS-DH-KR-ACP-KS-DH-KR-ACP-KS-DH-KR-ACP-KS-DH-KR-ER-KR-ACP-KS-DH-KR-ACP-KS-KR-ACP-ST-TE |
| PKZILLA-3 | KS-DH-ER-KR-ACP-KS-KR-ACP-KS-KR-ACP-KS-KR-ACP-KS-DH-MT-KR-ACP-ACP-KS-KR-ER-KR-ACP-KS-DH-KR-ACP-KS-DH-KR-ACP-KS-DH-ER-KR-ACP-KS-KR-FLX-ACP-KS-KR-ACP-KS-KR-ACP-KS-KR-ACP-KS-KR-ACP-KS-DH-KR-ACP-KS-DH-KR-ACP-KS-DH-KR-ACP-KS-KR-ACP-KS-DH-KR-ACP-ACP-S59-C |

**Table S4**: Linear sequence of polyketide synthase (PKS) domains within the PKZILLA polypeptides of *Prymnesium parvum* 12B1. Analogous rows for non-PKZILLA PKS polypeptides are not shown for brevity, but are available on Zenodo (doi:10.5281/zenodo.10028239).

| seqid | # of AA residues | M.W. (MDa) | # of modules | # of domains | # of ANLs | # of KSs | # of ACPs | # of KRs | # of DHs | # of ERs | # of FLXs | # of MTs | # of AMTs | # of STs | # of TEs | # of S59s | # of Cs |
|---|---|---|---|---|---|---|---|---|---|---|---|---|---|---|---|---|---|
| PKZILLA-1 | 45,212 | 4.73 | 35 | 140 | 1 | 34 | 45 | 33 | 16 | 5 | 5 | 1 | 0 | 0 | 0 | 0 | 0 |
| PKZILLA-2 | 30,685 | 3.23 | 23 | 99 | 0 | 21 | 32 | 20 | 14 | 4 | 0 | 0 | 6 | 1 | 1 | 0 | 0 |
| PKZILLA-3 | 24,439 | 2.59 | 19 | 76 | 0 | 19 | 21 | 20 | 9 | 3 | 1 | 1 | 0 | 0 | 0 | 1 | 1 |

**Table S5**: Counts of polyketide synthase (PKS) residues, domains, and modules within the PKZILLA polypeptides of *Prymnesium parvum* 12B1. AA=amino acid. M.W.=Molecular weight, calculated with ProtParam (https://web.expasy.org/protparam/). Analogous rows for non-PKZILLA PKS sequences are not shown for brevity, but are available on Zenodo (doi:10.5281/zenodo.10028239).

| cDUF ID | PKZILLA coordinates | Length | In module | Comments |
|---|---|---|---|---|
| PKZILLA-1__cDUF1 | PKZILLA-1:0-598 | 598 | N/A | At N-terminus of PKZILLA-1. InterProScan annotates as an intrinsically disordered region. |
| PKZILLA-1__cDUF10 | PKZILLA-1:38976-39480 | 504 | M30 | Sequence and structurally similar to FLX domains. Unclear if functional or vestigial. Dubbed cveFLX6 (cve=candidate, vestigial, extended) |
| PKZILLA-1__cDUF7 | PKZILLA-1:28567-28789 | 222 | M21 | Sequence similarity to FLX domains. Short, probably vestigial. |
| PKZILLA-1__cDUF9 | PKZILLA-1:34817-35027 | 210 | M27 | Sequence similarity to FLX domains. Short, probably vestigial. |
| PKZILLA-1__cDUF2 | PKZILLA-1:3324-3500 | 176 | M2 | Sequence similarity to KR domains. Nearby KRc. Likely an unannotated KRn structural subdomain. |
| PKZILLA-1__cDUF4 | PKZILLA-1:19100-19274 | 174 | M14 | Sequence similarity to KR domains. Nearby KRc. Likely an unannotated KRn structural subdomain. |
| PKZILLA-1__cDUF6 | PKZILLA-1:26033-26207 | 174 | M19 | Sequence similarity to KR domains. Nearby KRc. Likely an unannotated KRn structural subdomain. |
| PKZILLA-1__cDUF13 | PKZILLA-1:43935-44108 | 173 | M33 | Sequence similarity to FLX domains. Short, probably vestigial. |
| PKZILLA-1__cDUF3 | PKZILLA-1:17527-17697 | 170 | M13 | Sequence similarity to KR domains. Nearby KRf. Possibly an unannotated KRn structural subdomain. |
| PKZILLA-1__cDUF5 | PKZILLA-1:24460-24630 | 170 | M18 | Sequence similarity to KR domains. Nearby KRf. Possibly an unannotated KRn structural subdomain. |
| PKZILLA-1__cDUF12 | PKZILLA-1:41376-41541 | 165 | M31 | Sequence similarity to KR domains. Nearby KRc. Likely an unannotated KRn structural subdomain. |
| PKZILLA-1__cDUF8 | PKZILLA-1:31749-31903 | 154 | M24 | Weak sequence similarity to FLX4. Short, probably vestigial. |
| PKZILLA-1__cDUF11 | PKZILLA-1:39951-40061 | 110 | M30 | Some sequence similarity hits in 12B1, unclear role. |

**Table S6:** Analysis of InterProScan unannotated regions in PKZILLA-1 as candidate domains of unknown function (cDUFs). All contiguous unannotated regions of the PKZILLAs larger than 50 amino acid residues are shown, excepting those that seemed reasonably associated with an existing annotated domain (FLX extended C-terminal regions, i.e. eFLXs; doi:10.5281/zenodo.10120640, fig. S6). Regions were queried against other *P. parvum* 12B1 polypeptides using blastp. Source data, analysis scripts, and raw blast results are available on Zenodo (doi:10.5281/zenodo.10028178). For PKZILLA-2, see next table, and for PKZILLA-3 see Zenodo item.

| cDUF ID | PKZILLA coordinates | Length | In module | Comments |
|---|---|---|---|---|
| PKZILLA-2__cDUF17 | PKZILLA-2:27013-27222 | 209 | M54 | Sequence similarity to KR domains. Nearby KRc. Likely an unannotated KRn structural subdomain. |
| PKZILLA-2__cDUF4 | PKZILLA-2:5942-6137 | 195 | M39 | Sequence similarity to KR domains. Possibly an unannotated KRn structural subdomain. |
| PKZILLA-2__cDUF9 | PKZILLA-2:15752-15938 | 186 | M47/M48 | No hits to anything. (Including KSs, and KRs) |
| PKZILLA-2__cDUF2 | PKZILLA-2:322-499 | 177 | M34 | Sequence similarity to KR domains. Nearby KRc. Likely an unannotated KRn structural subdomain. |
| PKZILLA-2__cDUF8 | PKZILLA-2:13946-14120 | 174 | M46 | Sequence similarity to KR domains. Nearby KRc. Likely an unannotated KRn structural subdomain. |
| PKZILLA-2__cDUF7 | PKZILLA-2:11843-11970 | 127 | M44 | Sequence similarity to KS-KR-ACP-ACP modules. Possible structural role. |
| PKZILLA-2__cDUF3 | PKZILLA-2:2895-3016 | 121 | M36 | Sequence similarity to KS-KR-ACP-ACP modules. Possible structural role. |
| PKZILLA-2__cDUF12 | PKZILLA-2:19745-19845 | 100 | M50 | Associated with AMT domains. Possible structural role |
| PKZILLA-2__cDUF14 | PKZILLA-2:20741-20841 | 100 | M50 | Associated with AMT domains. Possible structural role |
| PKZILLA-2__cDUF10 | PKZILLA-2:18744-18839 | 95 | M50 | Multiple hits in PKZILLA-2, not within annotated domains or modules. Possible structural role. |
| PKZILLA-2__cDUF13 | PKZILLA-2:20243-20332 | 89 | M50 | Multiple hits in PKZILLA-2, possible structural role. |
| PKZILLA-2__cDUF15 | PKZILLA-2:21239-21328 | 89 | M50 | Multiple hits in PKZILLA-2, possible structural role. |
| PKZILLA-2__cDUF11 | PKZILLA-2:19248-19336 | 88 | M50 | Multiple hits in PKZILLA-2, possible structural role. |
| PKZILLA-2__cDUF16 | PKZILLA-2:21737-21809 | 72 | M50 | Multiple hits in PKZILLA-2, possible structural role. |
| PKZILLA-2__cDUF6 | PKZILLA-2:10441-10511 | 70 | M42/M43 | No hits. |
| PKZILLA-2__cDUF1 | PKZILLA-2:0-52 | 52 | M34 | Sequence similarity to KR domains. Possibly an unannotated KRn structural subdomain. Interesting as this is the N-terminus of PKZILLA-2, might have a protein-protein interaction role. |
| PKZILLA-2__cDUF5 | PKZILLA-2:10112-10164 | 52 | M42 | No hits. |

**Table S7:** Analysis of InterProScan unannotated regions in PKZILLA-2 as candidate domains of unknown function (cDUFs). All contiguous unannotated regions of the PKZILLAs larger than 50 amino acid residues are shown, excepting those that seemed reasonably associated with an existing annotated domain (FLX extended C-terminal regions, i.e. eFLXs; doi:10.5281/zenodo.10120640, fig. S6). Regions were queried against other *P. parvum* 12B1 polypeptides using blastp. Source data, analysis scripts, and raw blast results are available on Zenodo (doi:10.5281/zenodo.10028178). For PKZILLA-1 see previous table and for PKZILLA-3, see Zenodo item.

| module | seqid | start | end | components | KS# | ACP# | DH# | ER# | KR# | FLX# | comments |
|---|---|---|---|---|---|---|---|---|---|---|---|
| LM | PKZILLA-1 | 600 | 1242 | ANL-ACP | | ACP1 | | | | | Loading module |
| M1 | PKZILLA-1 | 1351 | 2591 | KS-DH-KR-ACP | KS1 | ACP2 | DH1 | | KR1 | | |
| M2 | PKZILLA-1 | 2624 | 4147 | KS-DH-ER-KR-ACP | KS2 | ACP3 | DH2 | ER1 | KR2c | | |
| M3 | PKZILLA-1 | 4168 | 5148 | KS-KR-ACP | KS3 | ACP4 | | | KR3 | | |
| M4 | PKZILLA-1 | 5168 | 6140 | KS-KR-ACP | KS4 | ACP5 | | | KR4 | | |
| M5 | PKZILLA-1 | 6160 | 7132 | KS-KR-ACP | KS5 | ACP6 | | | KR5 | | |
| M6 | PKZILLA-1 | 7152 | 8677 | KS-KR-FLX-ACP | KS6 | ACP7 | | | KR6 | FLX1 | |
| M7 | PKZILLA-1 | 8700 | 9673 | KS-KR-ACP | KS7 | ACP8 | | | KR7 | | |
| M8 | PKZILLA-1 | 9693 | 10900 | KS-DH-KR-ACP | KS8 | ACP9 | DH3 | | KR8 | | |
| M9 | PKZILLA-1 | 10918 | 12999 | KS-FLX-KR-FLX-ACP | KS9 | ACP10 | | | KR9 | FLX2<br>FLX3 | |
| M10 | PKZILLA-1 | 13023 | 14232 | KS-DH-KR-ACP | KS10 | ACP11 | DH4 | | KR10 | | |
| M11 | PKZILLA-1 | 14250 | 15807 | KS-FLX-KR-ACP-ACP | KS11 | ACP12<br>ACP13 | | | KR11 | FLX4 | |
| M12 | PKZILLA-1 | 15819 | 17041 | KS-DH-KR-ACP | KS12 | ACP14 | DH5 | | KR12 | | |
| M13 | PKZILLA-1 | 17062 | 18379 | KS-KR-ACP-ACP | KS13 | ACP15<br>ACP16 | | | KR13 | | Sat. bimodule (S2M)<br>M13/M14 |
| M14 | PKZILLA-1 | 18398 | 19912 | KS-DH-ER-KR-ACP | KS$^0$14 | ACP17 | DH6 | ER2 | KR14c0 | | (S2M) KR inactive via phylogeny |
| M15 | PKZILLA-1 | 19934 | 21162 | KS-DH-KR-ACP | KS15 | ACP18 | DH7 | | KR15 | | |
| M16 | PKZILLA-1 | 21180 | 22740 | KS-FLX-KR-ACP-ACP | KS16 | ACP19<br>ACP20 | | | KR16 | FLX5 | |
| M17 | PKZILLA-1 | 22751 | 23973 | KS-DH-KR-ACP | KS17 | ACP21 | DH8 | | KR17 | | |
| M18 | PKZILLA-1 | 23995 | 25312 | KS-KR-ACP-ACP | KS18 | ACP22<br>ACP23 | | | KR18 | | Sat. bimodule (S2M)<br>M18/M19 |
| M19 | PKZILLA-1 | 25331 | 26845 | KS-DH-ER-KR-ACP | KS$^0$19 | ACP24 | DH9 | ER3 | KR19c0 | | (S2M) KR inactive via phylogeny |
| M20 | PKZILLA-1 | 26867 | 28095 | KS-DH-KR-ACP | KS20 | ACP25 | DH10 | | KR20 | | |
| M21 | PKZILLA-1 | 28112 | 29438 | KS-KR-ACP-ACP | KS21 | ACP26<br>ACP27 | | | KR21 | | Pass-through bimodule (PT2M) |
| M22 | PKZILLA-1 | 29450 | 29999 | KS-ACP | KS$^0$22 | ACP28 | | | | | (PT2M) M21/M22 |
| M23 | PKZILLA-1 | 30017 | 31221 | KS-DH-KR-ACP | KS23 | ACP29 | DH11 | | KR22 | | |
| M24 | PKZILLA-1 | 31240 | 32555 | KS-KR-ACP-ACP | KS24 | ACP30<br>ACP31 | | | KR23 | | Pass-through bimodule (PT2M) |
| M25 | PKZILLA-1 | 32576 | 33116 | KS-ACP | KS$^0$25 | ACP32 | | | | | (PT2M) M24/M25 |
| M26 | PKZILLA-1 | 33134 | 34338 | KS-DH-KR-ACP | KS26 | ACP33 | DH12 | | KR24 | | |
| M27 | PKZILLA-1 | 34357 | 35691 | KS-KR-ACP-ACP | KS27 | ACP34<br>ACP35 | | | KR25 | | Sat. bimodule (S2M)<br>M27/M28 |
| M28 | PKZILLA-1 | 35709 | 37227 | KS-DH-KR-ER-KR-ACP | KS$^0$28 | ACP36 | DH13 | ER4 | KR26n0<br>KR27c0 | | S2M, KR27 inactive via phylogeny |
| M29 | PKZILLA-1 | 37252 | 38466 | KS-DH-KR-ACP | KS29 | ACP37 | DH14 | | KR28 | | |
| M30 | PKZILLA-1 | 38482 | 40909 | KS-DH-KR-MT-KR-ACP-ACP-ACP-ACP | KS30 | ACP38<br>ACP39<br>ACP40<br>ACP41 | DH15 | | KR29n0<br>KR30c | | (S2M), PKZILLA-1 cDUF10, cveFLX6 contained within |
| M31 | PKZILLA-1 | 40933 | 42203 | KS-ER-KR-ACP | KS$^0$31 | ACP42 | | ER5 | KR31c0 | | (S2M) M30/M31 |
| M32 | PKZILLA-1 | 42214 | 43440 | KS-DH-KR-ACP | KS32 | ACP43 | DH16 | | KR32 | | |
| M33 | PKZILLA-1 | 43458 | 44768 | KS-KR-ACP-ACP | KS33 | ACP44<br>ACP45 | | | KR33 | | Split sat. bimodule (S2M) M33/M34 |
| M34 | PKZILLA-1 | 44790 | 45210 | KS- | KS$^0$34 | | | | | | S2M, Split module (split w/ PKZILLA-2) |

**Table S8:** Organization of PKZILLA-1 domains into modules and the assignment of domain catalytic activity or inactivity based on active site analysis. Number, contents, and extents of modules have been modified from original bioinformatic predictions to account for non-standard or split module structures. Source data & code for bioinformatic predictions of modules are available at Zenodo (doi:10.5281/zenodo.10028239). Analogous module structures for PKZILLA-3 and non-PKZILLA polypeptides are not shown for brevity, but are available at Zenodo (doi:10.5281/zenodo.10028239). For PKZILLA-2, see next table. KR domains are annotated as KR/KRf, KRn, KRc depending on their respective classification as a full length non-split KR, a split KR N-terminal subdomain, or a split KR C-terminal subdomain, and "0" is appended to the domain name if the active site residues suggest the domain is catalytically inactive, or in case of KSs, if it is a predicted catalytically active but non-elongating domain. For further details on catalytic activity determination via active site analysis, see "Analysis of active site residues within PKZILLA PKS domains" in the Materials and Methods and the associated Zenodo item (doi:10.5281/zenodo.10028517).

| module cont. | module restart | seqid | start | end | components | KS# | ACP# | DH# | ER# | KR# | comments |
|---|---|---|---|---|---|---|---|---|---|---|---|
| M34 | (M1) | PKZILLA-2 | 52 | 1139 | -DH-ER-KR-ACP | | ACP1 | DH1 | ER1 | KR1c0 | Split module (split w/ PKZILLA-1). KR inactive via phylogeny. |
| M35 | (M2) | PKZILLA-2 | 1186 | 2382 | KS-DH-KR-ACP | KS1 | ACP2 | DH2 | | KR2 | |
| M36 | (M3) | PKZILLA-2 | 2400 | 3707 | KS-KR-ACP-ACP | KS2 | ACP3 ACP4 | | | KR3 | Pass-through bimodule (PT2M) M36/M37 |
| M37 | (M4) | PKZILLA-2 | 3735 | 4275 | KS-ACP | $KS^{0}3$ | ACP5 | | | | (PT2M) |
| M38 | (M5) | PKZILLA-2 | 4294 | 5497 | KS-DH-KR-ACP | KS4 | ACP6 | DH3 | | KR4 | |
| M39 | (M6) | PKZILLA-2 | 5519 | 6822 | KS-KR-ACP-ACP | KS5 | ACP7 ACP8 | | | KR5 | Dehydrating bimodule (DH2M) M39/M40 |
| M40 | (M7) | PKZILLA-2 | 6850 | 7654 | KS-DH-ACP | $KS^{0}6$ | ACP9 | DH4 | | | (DH2M) |
| M41 | (M8) | PKZILLA-2 | 7664 | 9233 | KS-DH-KR-ER-KR-ACP | KS7 | ACP10 | DH5 | ER2 | KR6n0 KR7c | ER2 is diverged from other ERs , ER* |
| M42 | (M9) | PKZILLA-2 | 9258 | 10437 | KS-KR-ACP-ACP-ACP | KS8 | ACP11 ACP12 ACP13 | | | KR8 | Dehydrating bimodule (DH2M) M42/M43 |
| M43 | (M10) | PKZILLA-2 | 10519 | 11331 | KS-DH-ACP | $KS^{0}9$ | ACP14 | DH6 | | | (DH2M) |
| M44 | (M11) | PKZILLA-2 | 11350 | 12655 | KS-KR-ACP-ACP | KS10 | ACP15 ACP16 | | | KR9 | Saturating trimodule (S3M) M44/M45/M46 |
| M45 | (M12) | PKZILLA-2 | 12678 | 13225 | KS-ACP | $KS^{0}11$ | ACP17 | | | | (S3M) |
| M46 | (M13) | PKZILLA-2 | 13243 | 14757 | KS-DH-ER-KR-ACP | $KS^{0}12$ | ACP18 | DH7 | ER3 | KR10c0 | (S3M), KR inactive via phylogeny |
| M47 | (M14) | PKZILLA-2 | 14779 | 15743 | KS-KR-ACP | KS13 | ACP19 | | | KR11* | KR*. Initiates polyether. |
| M48* | (M15*) | PKZILLA-2 | 15938 | 16942 | DH-KR-ACP-ACP-ACP | | ACP20 ACP21 ACP22 | DH8 | | KR12 | Missing KS. Unannotated DH- upstream-gap is PKZILLA-2__cDUF9 |
| M49 | (M16) | PKZILLA-2 | 16958 | 18303 | KS-DH-KR-ACP-ACP | KS14 | ACP23 ACP24 | DH9 | | KR13 | KS,DH diverged vs PKZILLA others |
| M50 | (M17) | PKZILLA-2 | 18323 | 22019 | KS-AMT-AMT-AMT-AMT-AMT-AMT-ACP-ACP | KS15 | ACP25 ACP26 | | | | AMT1, AMT2, …, AMT6 |
| M51 | (M18) | PKZILLA-2 | 22043 | 23279 | KS-DH-KR-ACP | KS16 | ACP27 | DH10 | | KR14 | |
| M52 | (M19) | PKZILLA-2 | 23307 | 24554 | KS-DH-KR-ACP | KS17 | ACP28 | DH11 | | KR15 | |
| M53 | (M20) | PKZILLA-2 | 24582 | 25829 | KS-DH-KR-ACP | KS18 | ACP29 | DH12 | | KR16 | |
| M54 | (M21) | PKZILLA-2 | 25857 | 27862 | KS-DH-KR-ER-KR-ACP | KS19 | ACP30 | DH13 | ER4 | KR17 KR18c0 | KR17 is full length (f). KR18 inactive via phylogeny |
| M55 | (M22) | PKZILLA-2 | 27918 | 29114 | KS-DH-KR-ACP | KS20 | ACP31 | DH14 | | KR19 | |
| M56 | (M23) | PKZILLA-2 | 29132 | 30667 | KS-KR-ACP-ST-TE | KS21 | ACP32 | | | KR20* | ST-TE=Curacin/ CurM-like termination didomain. KR* |

**Table S9:** Organization of PKZILLA-2 domains into modules and the assignment of domain catalytic activity or inactivity based on active site analysis. Number, contents, and extents of modules have been modified from original bioinformatic predictions to account for non-standard or split module structures. Source data & code for bioinformatic predictions of modules are available at Zenodo (doi:10.5281/zenodo.10028239). Analogous module structures for PKZILLA-3 and non-PKZILLA polypeptides are not shown for brevity, but are available at Zenodo (doi:10.5281/zenodo.10028239). For PKZILLA-1, see previous table. KR domains are annotated as KR/KRf, KRn, KRc depending on their respective classification as a full length non-split KR, a split KR N-terminal subdomain, or a split KR C-terminal subdomain, and a 0 is appended to the domain name if the active site residues suggest the domain is catalytically inactive, or in case of KSs, if it is a predicted catalytically active but non-elongating domain. For further details on catalytic activity determination via active site analysis, see “Analysis of active site residues within PKZILLA PKS domains” in the Materials and Methods and the associated Zenodo item (doi:10.5281/zenodo.10028517)

| Uniprot taxon mnemonic | informative taxonomic parent | explanation |
|---|---|---|
| PKZILLA | haptophyte algae | Domain arising from a PKZILLA gene from *Prymnesium parvum* strain 12B1 (haptophyte) |
| 9ACTN | bacteria | Descended taxon of Actinomycetota |
| 9CYAN | bacteria | Descended taxon of Cyanobacteriota |
| 9DINO | dinoflagellate algae | Descended taxon of Dinophyceae (dinoflagellates) |
| ASPFN | fungi | *Aspergillus flavus* |
| ASPFU | fungi | *Aspergillus fumigatus* |
| ASPPU | fungi | *Aspergillus parasiticus* |
| ASPTE | fungi | *Aspergillus terreus* |
| ASPTN | fungi | *Aspergillus terreus* (strain NIH 2624 / FGSC A1156) |
| CHLRE | green algae | *Chlamydomonas reinhardtii* |
| DOTSN | fungi | *Dothistroma septosporum* (strain NZE10 / CBS 128990) aka Red band needle blight fungus |
| EMENI | fungi | *Emericella nidulans* |
| EMIHU | haptophyte algae | *Emiliania huxleyi* (haptophyte) |
| HUMAN | animals | *Homo sapiens* |
| MONPI | fungi | *Monascus pilosus* aka Red mold |
| MYCBO | bacteria | *Mycobacterium bovis* (strain ATCC BAA-935 / AF2122/97) |
| MYCMM | bacteria | *Mycobacterium marinum* (strain ATCC BAA-535 / M) |
| PENAE | fungi | *Penicillium aethiopicum* (fungi) |
| SACER | bacteria | *Saccharopolyspora erythraea* - erythromycin producer |
| STRAT | bacteria | *Streptomyces antibioticus* - oleandomycin producer |
| STRVZ | bacteria | *Streptomyces venezuelae* - pikromycin producer |

**Table S10**: Mapping of uniprot taxonomy mnemonics to human readable descriptions. Key for taxonomic interpretation of fig. S16, S17, S18, S19.

| seqid | module | MSA sequence context around diagnostic site (+/- 10 residues) | residue at diagnostic site | predicted stereochemical outcome | If DH present, proposed ene outcome | comment |
|---|---|---|---|---|---|---|
| A9GJ40 | N/A | VIHSAIVLR-**D**RSLREMDEP- | D | *(R)* | | seqid is Uniprot ID. Corresponds to *(R)* hydroxyl forming KR "1. Etn_KR12_D-OH" from Supplementary Figure 4 of (*32*) |
| A1KQR8 | N/A | AIFSGMVFDF**E**NSIQQTSEA- | E | *(S)* | | seqid is Uniprot ID. Corresponds to *(S)* hydroxyl forming KR "12. rhi_KR3_L-OH" from Supplementary Figure 4 of (*32*) |
| PKZILLA-1__KR1 | M1 | FWHTAGVLF-**D**ALVKKQNAV- | D | *(R)* | *(E)*- | |
| PKZILLA-1__KR2c | M2 | VWHAAGVLA-**D**SMLPKQTAL- | D | *(R)* | | |
| PKZILLA-1__KR3 | M3 | VLHAAGYAV-**D**TLSIDLVAR- | D | *(R)* | | |
| PKZILLA-1__KR4 | M4 | MLHAAGASD-**K**GLLLDIVAR- | K | *(S)* | | |
| PKZILLA-1__KR5 | M5 | MLHAAGVGD-**K**GLLLDIVAR- | K | *(S)* | | |
| PKZILLA-1__KR6 | M6 | IFHMTEVLI-**D**KLIFLMPAR- | D | *(R)* | | |
| PKZILLA-1__KR7 | M7 | VLHAAGVGD-**K**GLLLDIVAR- | K | *(S)* | | |
| PKZILLA-1__KR8 | M8 | VWHAAGALA-**D**SLLPSLRAE- | D | *(R)* | *(E)*- | |
| PKZILLA-1__KR9 | M9 | VFHMTEILQ-**D**KLIFGMTSK- | D | *(R)* | | |
| PKZILLA-1__KR10 | M10 | LWHAAGVLV-**D**GVLSKQTAR- | D | *(R)* | *(E)*- | |
| PKZILLA-1__KR11 | M11 | VLHAAGTGD-**K**GLLLELVSL- | K | *(S)* | | |
| PKZILLA-1__KR12 | M12 | VWHAAGVLA-**D**AILHKQTAR- | D | *(R)* | *(E)*- | |
| PKZILLA-1__KR13 | M13 | ILHAAGVLR-**D**ALLRNTRAA- | D | *(R)* | | |
| PKZILLA-1__KR14c0 | M14 | VWHAAGM----SESMQQHTH- | - | N/A |  |  |
| PKZILLA-1__KR15 | M15 | VWHASGVLA-**D**GMVPKQDSR- | D | *(R)* | *(E)*- | |
| PKZILLA-1__KR16 | M16 | VLHAAGTGD-**K**GLLLELVSL- | K | *(S)* | | |
| PKZILLA-1__KR17 | M17 | VWHAAGVLA-**D**AILHKQTAR- | D | *(R)* | *(E)*- | |
| PKZILLA-1__KR18 | M18 | ILHAAGVLR-**D**ALLRNTRAA- | D | *(R)* | | |
| PKZILLA-1__KR19c0 | M19 | VWHAAGM----SESMQQHTH- | - | N/A |  |  |
| PKZILLA-1__KR20 | M20 | VWHASGVLA-**D**GMVPKQDSR- | D | *(R)* | *(E)*- | |
| PKZILLA-1__KR21 | M21 | ILHAAGVLR-**D**ALLRNTRAA- | D | *(R)* | | |
| PKZILLA-1__KR22 | M23 | VWHAAGVLA-**D**GMLAKQSAR- | D | *(R)* | *(E)*- | |
| PKZILLA-1__KR23 | M24 | ILHAAGVLR-**D**ALLRNTRAA- | D | *(R)* | | |
| PKZILLA-1__KR24 | M26 | VWHAAGVLA-**D**GMLAKQSAR- | D | *(R)* | *(E)*- | |
| PKZILLA-1__KR25 | M27 | ILHAAGVLR-**D**ALLRNTRAA- | D | *(R)* | | |
| PKZILLA-1__KR26n0 | M28 | --------------------- | - | N/A |  |  |
| PKZILLA-1__KR27c0 | M28 | VWHTSSSFY-**D**AEFLQQNAL- | D | *(R)* | | Note: predicted to be inactive. |
| PKZILLA-1__KR28 | M29 | VWHAAGVLA-**D**ALLPNQTAM- | D | *(R)* | *(E)*- | |
| PKZILLA-1__KR29n0 | M30 | --------------------- | - | N/A |  |  |
| PKZILLA-1__KR30c | M30 | IFHAAHRLA-**D**AVLANQKAT- | D | *(R)* | | |
| PKZILLA-1__KR31c0 | M31 | VWHTASSLS-**E**ASIPTTTSAV | E | *(S)* | | Note: predicted to be inactive. |
| PKZILLA-1__KR32 | M32 | LWHAAGVVS-**D**GVLAKQSAQ- | D | *(R)* | *(E)*- | |
| PKZILLA-1__KR33 | M33 | VLHAAGVLW-**D**ALLRNTRAP- | D | *(R)* | | |
| PKZILLA-2__KR1c0 | M34 | VWHTMGVMS-**S**AELNQQDAL- | S | *(S)* | | Note: predicted to be inactive. |
| PKZILLA-2__KR2 | M35 | LWHAAGLLS-**D**GLIPKQTAR- | D | *(R)* | *(E)*- | |
| PKZILLA-2__KR3 | M36 | VLHAAGVLR-**D**SLLRNTRAQ- | D | *(R)* | | |
| PKZILLA-2__KR4 | M38 | VWHAAGVLA-**D**GMLAKQSAR- | D | *(R)* | *(E)*- | |
| PKZILLA-2__KR5 | M39/M40 | VLHAAGVLR-**D**SLLRNTRAQ- | D | *(R)* | *(E)*- | |
| PKZILLA-2__KR6n0 | M41 | --------------------- | - | N/A |  |  |

| | | | | | | |
|---|---|---|---|---|---|---|
| PKZILLA-2__KR7c | M41 | VWHAAGVLA-**D**SLLPNQRAQ- | D | *(R)* | | |
| PKZILLA-2__KR8 | M42/M43 | FVHASGVLL-**D**SLCRNISLM- | D | *(R)* | *(E)*- | |
| PKZILLA-2__KR9 | M44 | VLHAAGVLR-**D**SLLRNTRAQ- | D | *(R)* | | |
| PKZILLA-2__KR10c0 | M46 | VWHAAGM----SEFMQQHTH- | - | N/A |  |  |
| PKZILLA-2__KR11* | M47 | VLHMPRAVE-**E**RVLVYLGAR- | E | *(S)* | | Note: possibly novel catalysis or inactive. Is a KR* |
| PKZILLA-2__KR12 | M48* | IWHAAGVIV-**D**ALLPKQTSA- | D | *(R)* | *(E)*- | |
| PKZILLA-2__KR13 | M49 | VWHAAGVLA-**D**AVLAKQDEH- | D | *(R)* | *(E)*- | |
| PKZILLA-2__KR14 | M51 | VWHAAGVLA-**D**AVLPNQTAA- | D | *(R)* | *(E)*- | |
| PKZILLA-2__KR15 | M52 | VWHAAGVLA-**D**AVLPKQTAA- | D | *(R)* | *(E)*- | |
| PKZILLA-2__KR16 | M53 | AWHSAGVLA-**D**AVLPKQTAA- | D | *(R)* | *(E)*- | |
| PKZILLA-2__KR17 | M54 | AWHSAGVLA-**D**AVLPKQTAA- | D | *(R)* | | |
| PKZILLA-2__KR18c0 | M54 | VWHTMGVMS-**N**AELNQQDAL- | N | *(S)* | | Note: predicted to be inactive. |
| PKZILLA-2__KR19 | M55 | LWHAAGLLS-**D**GLIPKQTAR- | D | *(R)* | *(E)*- | |
| PKZILLA-2__KR20* | M56 | VLQMWSVQR-**D**VALKGLTCR- | D | *(R)* | *(E)*- | Note: possibly novel catalysis or inactive. Is a KR* |

**Table S11:** Bioinformatic assignment of stereochemical outcome for PKZILLA-1/-2 ketoreductases (KRs). An *(R)* or *(S)* stereochemical outcome of ketoreduction at the hydroxyl-bound carbon (fig. S20) was predicted based on the reported heuristic of stereochemical outcome for *trans*-AT KRs (Supplementary Figure 4 of (*32*)), wherein an aspartate (D) at the designated site is predictive of *(R)* stereochemistry, while any other residue is predictive of *(S)* stereochemistry. If a DH domain was present in the module, an *(E)*- or *(Z)*- outcome for alkene formation was predicted based on the published heuristic (*94*) that an *(R)* hydroxyl is predictive of *(E)*- while an *(S)* hydroxyl is predictive of *(Z)*-. Source data, analysis code, and results are available on Zenodo (doi:10.5281/zenodo.10569208).

| Carbon with stereocenter | Type | Empirically reported stereochemistry* | Responsible module | Mapped stereochemically responsible domain(s) | Bioinformatically predicted stereochemical outcome** | Predicted stereochemistry matches empirical? |
|---|---|---|---|---|---|---|
| C85 | "α"-Cl | *(S)* | *n.d.* [**M2**] | N/A | *n.d.* | |
| C84 | β-OH | *n.d.* | **M3** | PKZILLA-1__KR3 | *(R)* | |
| C83 | α-OH | *n.d.* | *n.d.* [**M3**] | N/A | *n.d.* | |
| C82 | β-OH | *n.d.* | **M4** | PKZILLA-1__KR4 | *(S)* | |
| C81 | α-OH | *n.d.* | *n.d.* [**M5**] | N/A | *n.d.* | |
| C80 | β-OH | *n.d.* | **M5** | PKZILLA-1__KR5 | *(S)* | |
| C78 | β-OH | *n.d.* | **M6** | PKZILLA-1__KR6 | *(R)* | |
| C77 | α-OH | *n.d.* | **M6** | PKZILLA-1__KR6 /FLX1 | *n.d.* | |
| C76 | β-OH | *n.d.* | **M7** | PKZILLA-1__KR7 | *(S)* | |
| C72 | β-OH | *(R)* | **M9** | PKZILLA-1__KR9 | *(R)* | Yes |
| C71 | α-OH | *(S)* | **M9** | PKZILLA-1__KR9 /FLX2/3 | *n.d.* | |
| C68 | β-OH | *(S)* | **M11** | PKZILLA-1__KR11 | *(S)* | Yes |
| C67 | α-OH | *(S)* | **M11** | PKZILLA-1__KR11/FLX4 | *n.d.* | |
| C60 | β-OH | *(S)* | **M16** | PKZILLA-1__KR16 | *(S)* | Yes |
| C56 | "β"-Cl | *(R)* | *n.d.* [**M19**] | N/A | *n.d.* | |
| C52 | β-OH | *(R)* | **M21**/M22 | PKZILLA-1__KR21 | *(R)* | Yes |
| C48 | β-OH | *(R)* | **M24**/M25 | PKZILLA-1__KR23 | *(R)* | Yes |
| C39 | "α"-$CH_3$ | *(R)* | **M30** | PKZILLA-2__MT1/ER5 | *n.d.* | |
| C32 | β-OH | *(S)* | **M36**/M37 | PKZILLA-2__KR3 | *(R)* | No |
| C20 | β-OH | *(S)* | **M47** | PKZILLA-2__KR11* | *(S)* | Yes |
| C14 | "β"-amine | *(R)* | **M50** | PKZILLA-2__AMT1… | *n.d.* | |

**Table S12:** Comparison of stereochemical assignments from structural elucidations and retrobiosynthetic inferences with bioinformatically predicted stereochemical assignments. *=Stereochemistry determined from retrobiosynthetic inferences of reported prymnesin stereochemistry (Fig. 3) onto the proposed pre-prymnesin biosynthetic precursor (PPBP), with (*R*) or (*S*) stereochemistry calculated on a standardized polyketide chain elongation thioester intermediate (fig. S20). Due to an inversion of Cahn–Ingold–Prelog priority when γ,δ bond is an alkene vs saturated, or if the γ carbon has a hydroxyl, γ,δ carbons were drawn as always saturated and without hydroxyls for the purposes of stereochemical assignment. **=See previous Table. *n.d.* = not determined . N/A = not applicable.

| Double bonded carbons in PA | Empirically reported (*E*)-ene or (*Z*)-ene isomer | Responsible module | Mapped responsible domain(s) for isomeric outcome on PPBP | Bioinformatically predicted isomeric outcome** | Predicted isomer matches empirical? |
|---|---|---|---|---|---|
| C21-C22 | *(E)-* | **M49** | *n.d.* | *n.d.* | |
| C23-C24 | *(E)-* | **M49** | *n.d.* | *n.d.* | |
| C12-C11 | *(E)-* | **M51** | PKZILLA-2__KR14/DH10 | (*E*)- | Yes |
| C10-C9 | *(E)-* | **M52** | PKZILLA-2__KR15/DH11 | (*E*)- | Yes |
| C8-C7 | (alkyne) | **M53** | PKZILLA-2__KR16/DH12 | (*E*)- | *n.d.* |

**Table S13:** Comparison of (*E*)-ene vs (*Z*)-ene isomer observations on prymnesin from structural elucidations with bioinformatically predicted isomeric outcomes on PPBP. **=See Table S11. *n.d.* = not determined, due to lack of bioinformatic heuristic for prediction of isomeric outcome for vinylogous dehydration. N/A = not applicable. PA=prymnesin aglycone (see Fig. 3), PPBP=pre-prymnesin biosynthetic precursor (see Fig. 3).

| | Title of Dataset or Summary for Github repository | Category | DOI or URL |
|---|---|---|---|
| 1 | Version v1.1 of the reference genome for Prymnesium parvum 12B1 (12B1_scaffolds_v1.1.fasta) | Genomics | https://doi.org/10.5281/zenodo.10023322 |
| 2 | Version 1.1 of the Prymnesium parvum 12B1 gene annotation (12B1_v1.1.gff3) | Genomics | https://doi.org/10.5281/zenodo.10023330 |
| 3 | Hotspots of PKS coding evidence within the Prymnesium parvum 12B1 v1.1 genome assembly | Genomics | https://doi.org/10.5281/zenodo.10309063 |
| 4 | Kallisto v0.50.0 expression quantification of day and night phase rRNA depletion RNA-Seq data from *P. parvum* strain 12B1 | Transcriptomics | https://doi.org/10.5281/zenodo.10023426 |
| 5 | Detailed mass spectrometry instrument method text description, used to produce proteomic data files for PKZILLA manuscript | Proteomics | https://doi.org/10.5281/zenodo.10023360 |
| 6 | SearchGUI proteomic search analysis for PKZILLAs in P. parvum 12B1 | Proteomics | https://doi.org/10.5281/zenodo.10023441 |
| 7 | Scripts and analysis files for categorization of PKZILLA matching proteomic peptides into protein-unique, protein-multimatch & exon-unique, exon-multimatch categories. | Proteomics | https://doi.org/10.5281/zenodo.10028959 |
| 8 | A Nextflow pipeline that wraps InterProScan , including making it parallelized & doing some plotting | Supporting code | https://github.com/photocyte/interproscan_parallel |
| 9 | InterProScan domain annotation on non-PKZILLA polypeptides | PKS analysis | https://doi.org/10.5281/zenodo.10011739 |
| 10 | InterProScan domain annotation of PKZILLA polypeptides | PKS analysis | https://doi.org/10.5281/zenodo.10023460 |
| 11 | PKZILLA candidate domains of unknown function (cDUFs) analysis | PKS analysis | https://doi.org/10.5281/zenodo.10028178 |
| 12 | Prymnesium parvum 12B1 polyketide synthase (PKS) domain and module analysis | PKS analysis | https://doi.org/10.5281/zenodo.10028239 |
| 13 | Tabular analysis of polyketide synthase domain active site residues in the PKZILLAs | PKS analysis | https://doi.org/10.5281/zenodo.10028517 |
| 14 | Bioinformatic assignment of PKZILLA ketoreductase (KR) stereochemical outcome based on application of a literature trans-AT KR diagnostic residue heuristic | PKS analysis | https://doi.org/10.5281/zenodo.10569208 |
| 15 | Domain phylogenetics for the PKZILLAs | PKS analysis | https://doi.org/10.5281/zenodo.10152638 |
| 16 | A wrapper around some simple Python scripts, to query the InterPro API with respect to fetching domains from arbitrary taxonomic groups, ranging from all life to specific strains | Supporting code | https://github.com/photocyte/InterPro_API_domain_toolkit |
| 17 | PKZILLA domain phylogenies with outgroup representatives from Uniprot | PKS analysis | https://doi.org/10.5281/zenodo.10247217 |
| 18 | Non-length-normalized graphical plots of the PKZILLA domains, modules, cDUFs, and exon extents | PKS analysis | https://doi.org/10.5281/zenodo.10093617 |
| 19 | Scripts, data, and analyses for extending flavoprotein (FLX) domains C-terminally to form extended FLX domains (eFLX) & performing structural predictions & alignments using AlphaFold2, RoseTTAFold, FoldSeek, and DALI | PKS analysis | https://doi.org/10.5281/zenodo.10120640 |

**Table S14:** Extended analyses, datasets, and code available on Zenodo or Github.

#### References:


1. G. M. Hallegraeff, D. M. Anderson, K. Davidson, F. Gianella, P. Hansen, *Fish-Killing Marine Algal Blooms: Causative Organisms, Ichthyotoxic Mechanisms, Impacts and Mitigation.* (UNESCO, Paris, France, 2023; https://dx.doi.org/10.25607/OBP-1964)*IOC Manuals and Guides*.

2. J. Sobieraj, D. Metelski, Insights into Toxic *Prymnesium parvum* Blooms as a Cause of the Ecological Disaster on the Odra River. *Toxins* **15**, 403 (2023).

3. D. M. Anderson, E. Fensin, C. J. Gobler, A. E. Hoeglund, K. A. Hubbard, D. M. Kulis, J. H. Landsberg, K. A. Lefebvre, P. Provoost, M. L. Richlen, J. L. Smith, A. R. Solow, V. L. Trainer, Marine harmful algal blooms (HABs) in the United States: History, current status and future trends. *Harmful Algae* **102**, 101975 (2021).

4. K. C. Nicolaou, M. O. Frederick, R. J. Aversa, The Continuing Saga of the Marine Polyether Biotoxins. *Angew. Chem. Int. Ed.* **47**, 7182–7225 (2008).

5. T. Igarashi, M. Satake, T. Yasumoto, Structures and Partial Stereochemical Assignments for Prymnesin-1 and Prymnesin-2: Potent Hemolytic and Ichthyotoxic Glycosides Isolated from the Red Tide Alga *Prymnesium parvum*. *J. Am. Chem. Soc.* **121**, 8499–8511 (1999).

6. R. E. Moore, G. Bartolini, Structure of palytoxin. *J. Am. Chem. Soc.* **103**, 2491–2494 (1981).

7. M. Murata, H. Naoki, T. Iwashita, S. Matsunaga, M. Sasaki, A. Yokoyama, T. Yasumoto, Structure of maitotoxin. *J. Am. Chem. Soc.* **115**, 2060–2062 (1993).

8. K. Nakanishi, The chemistry of brevetoxins: A review. *Toxicon* **23**, 473–479 (1985).

9. I. Vilotijevic, T. F. Jamison, Epoxide-Opening Cascades Promoted by Water. *Science* **317**, 1189–1192 (2007).

10. A. Nivina, K. P. Yuet, J. Hsu, C. Khosla, Evolution and Diversity of Assembly-Line Polyketide Synthases. *Chem. Rev.* **119**, 12524–12547 (2019).

11. K. Anestis, G. S. Kohli, S. Wohlrab, E. Varga, T. O. Larsen, P. J. Hansen, U. John, Polyketide synthase genes and molecular trade-offs in the ichthyotoxic species *Prymnesium parvum*. *Sci. Total Environ.* **795**, 148878 (2021).

12. F. M. V. Dolah, J. S. Morey, S. Milne, A. Ung, P. E. Anderson, M. Chinain, Transcriptomic analysis of polyketide synthases in a highly ciguatoxic dinoflagellate, *Gambierdiscus polynesiensis* and low toxicity *Gambierdiscus pacificus*, from French Polynesia. *PLOS ONE* **15**, e0231400 (2020).

13. N. Heimerl, E. Hommel, M. Westermann, D. Meichsner, M. Lohr, C. Hertweck, A. R. Grossman, M. Mittag, S. Sasso, A giant type I polyketide synthase participates in zygospore maturation in *Chlamydomonas reinhardtii*. *Plant J.* **95**, 268–281 (2018).

14. M. H. Medema, T. de Rond, B. S. Moore, Mining genomes to illuminate the specialized chemistry of life. *Nat. Rev. Genet.* **22**, 553–571 (2021).

15. M.-L. Bang, T. Centner, F. Fornoff, A. J. Geach, M. Gotthardt, M. McNabb, C. C. Witt, D. Labeit, C. C. Gregorio, H. Granzier, S. Labeit, The Complete Gene Sequence of Titin, Expression of an Unusual ≈700-kDa Titin Isoform, and Its Interaction With Obscurin Identify a Novel Z-Line to I-Band Linking System. *Circ. Res.* **89**, 1065–1072 (2001).

16. M. Sasaki, N. Takeda, H. Fuwa, R. Watanabe, M. Satake, Y. Oshima, Synthesis of the JK/LM-ring model of prymnesins, potent hemolytic and ichthyotoxic polycyclic ethers isolated from the red tide alga

*Prymnesium parvum*: confirmation of the relative configuration of the K/L-ring juncture. *Tetrahedron Lett.* **47**, 5687–5691 (2006).

17. T. Hashimoto, J. Hashimoto, I. Kozone, K. Amagai, T. Kawahara, S. Takahashi, H. Ikeda, K. Shin-ya, Biosynthesis of Quinolidomicin, the Largest Known Macrolide of Terrestrial Origin: Identification and Heterologous Expression of a Biosynthetic Gene Cluster over 200 kb. *Org. Lett.* **20**, 7996–7999 (2018).

18. S. Donadio, M. Staver, J. McAlpine, S. Swanson, L. Katz, Modular organization of genes required for complex polyketide biosynthesis. *Science* **252**, 675–679 (1991).

19. S. B. Binzer, D. K. Svenssen, N. Daugbjerg, C. Alves-de-Souza, E. Pinto, P. J. Hansen, T. O. Larsen, E. Varga, A-, B- and C-type prymnesins are clade specific compounds and chemotaxonomic markers in *Prymnesium parvum*. *Harmful Algae* **81**, 10–17 (2019).

20. J. H. Wisecaver, R. P. Auber, A. L. Pendleton, N. F. Watervoort, T. R. Fallon, O. L. Riedling, S. R. Manning, B. S. Moore, W. W. Driscoll, Extreme genome diversity and cryptic speciation in a harmful algal-bloom-forming eukaryote. *Curr. Biol.* **33**, 2246-2259.e8 (2023).

21. J. H. Wisecaver, J. D. Hackett, Dinoflagellate Genome Evolution. *Annu. Rev. Microbiol.* **65**, 369–387 (2011).

22. H.-H. Hong, H.-G. Lee, J. Jo, H. M. Kim, S.-M. Kim, J. Y. Park, C. B. Jeon, H.-S. Kang, M. G. Park, C. Park, K. Y. Kim, H.-H. Hong, H.-G. Lee, J. Jo, H. M. Kim, S.-M. Kim, J. Y. Park, C. B. Jeon, H.-S. Kang, M. G. Park, C. Park, K. Y. Kim, The exceptionally large genome of the harmful red tide dinoflagellate *Cochlodinium polykrikoides* Margalef (Dinophyceae): determination by flow cytometry. *Algae* **31**, 373–378 (2016).

23. National Center for Biotechnology Information (NCBI), Sequence Read Archive (SRA), *Prymnesium parvum* strain 12B Illumina reads - SRA PRJNA201451, SRR1685644. https://www.ncbi.nlm.nih.gov/sra/?term=SRR1685644.

24. Y. Dou, Y. Liu, X. Yi, L. K. Olsen, H. Zhu, Q. Gao, H. Zhou, B. Zhang, SEPepQuant enhances the detection of possible isoform regulations in shotgun proteomics. *Nat. Commun.* **14**, 5809 (2023).

25. P. Jones, D. Binns, H.-Y. Chang, M. Fraser, W. Li, C. McAnulla, H. McWilliam, J. Maslen, A. Mitchell, G. Nuka, S. Pesseat, A. F. Quinn, A. Sangrador-Vegas, M. Scheremetjew, S.-Y. Yong, R. Lopez, S. Hunter, InterProScan 5: genome-scale protein function classification. *Bioinformatics* **30**, 1236–1240 (2014).

26. F. Hemmerling, K. E. Lebe, J. Wunderlich, F. Hahn, An Unusual Fatty Acyl:Adenylate Ligase (FAAL)–Acyl Carrier Protein (ACP) Didomain in Ambruticin Biosynthesis. *ChemBioChem* **19**, 1006–1011 (2018).

27. F. Hemmerling, R. A. Meoded, A. E. Fraley, H. A. Minas, C. L. Dieterich, M. Rust, R. Ueoka, K. Jensen, E. J. N. Helfrich, C. Bergande, M. Biedermann, N. Magnus, B. Piechulla, J. Piel, Modular Halogenation, α-Hydroxylation, and Acylation by a Remarkably Versatile Polyketide Synthase. *Angew. Chem. Int. Ed.* **61**, e202116614 (2022).

28. A. J. Winter, R. N. Khanizeman, A. M. C. Barker-Mountford, A. J. Devine, L. Wang, Z. Song, J. A. Davies, P. R. Race, C. Williams, T. J. Simpson, C. L. Willis, M. P. Crump, Structure and Function of the α-Hydroxylation Bimodule of the Mupirocin Polyketide Synthase. *Angew. Chem. Int. Ed.* **62**, e202312514 (2023).

29. E. J. N. Helfrich, J. Piel, Biosynthesis of polyketides by trans-AT polyketide synthases. *Nat. Prod. Rep.* **33**, 231–316 (2016).

30. D. T. Wagner, J. Zeng, C. B. Bailey, D. C. Gay, F. Yuan, H. R. Manion, A. T. Keatinge-Clay, Structural and Functional Trends in Dehydrating Bimodules from trans-Acyltransferase Polyketide Synthases. *Structure* **25**, 1045-1055.e2 (2017).

31. J. Masschelein, P. K. Sydor, C. Hobson, R. Howe, C. Jones, D. M. Roberts, Z. Ling Yap, J. Parkhill, E. Mahenthiralingam, G. L. Challis, A dual transacylation mechanism for polyketide synthase chain release in enacyloxin antibiotic biosynthesis. *Nat. Chem.* **11**, 906–912 (2019).

32. E. J. N. Helfrich, R. Ueoka, A. Dolev, M. Rust, R. A. Meoded, A. Bhushan, G. Califano, R. Costa, M. Gugger, C. Steinbeck, P. Moreno, J. Piel, Automated structure prediction of trans-acyltransferase polyketide synthase products. *Nat. Chem. Biol.* **15**, 813–821 (2019).

33. F. Taft, M. Brünjes, T. Knobloch, H. G. Floss, A. Kirschning, Timing of the Δ10,12-Δ11,13 Double Bond Migration During Ansamitocin Biosynthesis in *Actinosynnema pretiosum*. *J. Am. Chem. Soc.* **131**, 3812–3813 (2009).

34. G. Hibi, T. Shiraishi, T. Umemura, K. Nemoto, Y. Ogura, M. Nishiyama, T. Kuzuyama, Discovery of type II polyketide synthase-like enzymes for the biosynthesis of cispentacin. *Nat. Commun.* **14**, 8065 (2023).

35. L. Gu, B. Wang, A. Kulkarni, J. J. Gehret, K. R. Lloyd, L. Gerwick, W. H. Gerwick, P. Wipf, K. Håkansson, J. L. Smith, D. H. Sherman, Polyketide Decarboxylative Chain Termination Preceded by O-Sulfonation in Curacin A Biosynthesis. *J. Am. Chem. Soc.* **131**, 16033–16035 (2009).

36. Y. Jiang, A. Kim, C. Olive, J. C. Lewis, Selective C-H Halogenation of Alkenes and Alkynes Using Flavin-Dependent Halogenases. ChemRxiv [Preprint] (2023). https://doi.org/10.26434/chemrxiv-2023-23r4l.

37. T. Hua, D. Wu, W. Ding, J. Wang, N. Shaw, Z.-J. Liu, Studies of Human 2,4-Dienoyl CoA Reductase Shed New Light on Peroxisomal β-Oxidation of Unsaturated Fatty Acids. *J. Biol. Chem.* **287**, 28956–28965 (2012).

38. Yong-Yeng Lin, Martin Risk, Sammy M. Ray, Donna Van Engen, Jon Clardy, Jerzy Golik, John C. James, Koji Nakanishi, Isolation and structure of brevetoxin B from the “red tide” dinoflagellate *Ptychodiscus brevis* (*Gymnodinium breve*). *J. Am. Chem. Soc.* **103**, 6773–6775 (1981).

39. Z.-P. Jiang, S.-H. Sun, Y. Yu, A. Mándi, J.-Y. Luo, M.-H. Yang, T. Kurtán, W.-H. Chen, L. Shen, J. Wu, Discovery of benthol A and its challenging stereochemical assignment: opening up a new window for skeletal diversity of super-carbon-chain compounds. *Chem. Sci.* **12**, 10197–10206 (2021).

40. J. Cortes, P. Schöffski, B. A. Littlefield, Multiple modes of action of eribulin mesylate: Emerging data and clinical implications. *Cancer Treat. Rev.* **70**, 190–198 (2018).

41. F. M. V. Dolah, G. S. Kohli, J. S. Morey, S. A. Murray, Both modular and single-domain Type I polyketide synthases are expressed in the brevetoxin-producing dinoflagellate, Karenia brevis (Dinophyceae). *J. Phycol.* **53**, 1325–1339 (2017).

42. J. Jian, Z. Wu, A. Silva-Núñez, X. Li, X. Zheng, B. Luo, Y. Liu, X. Fang, C. T. Workman, T. O. Larsen, P. J. Hansen, E. C. Sonnenschein, Long-read genome sequencing provides novel insights into the harmful algal bloom species *Prymnesium parvum*. *Sci. Total Environ.*, 168042 (2023).

43. R. M. Van Wagoner, M. Satake, J. L. C. Wright, Polyketide biosynthesis in dinoflagellates: what makes it different? *Nat. Prod. Rep.*, 37 (2014).

44. R. Teufel, A. Miyanaga, Q. Michaudel, F. Stull, G. Louie, J. P. Noel, P. S. Baran, B. Palfey, B. S. Moore, Flavin-mediated dual oxidation controls an enzymatic Favorskii-type rearrangement. *Nature* **503**, 552–556 (2013).

45. S. Guo, Y. Sang, C. Zheng, X.-S. Xue, Z. Tang, W. Liu, Enzymatic α-Ketothioester Decarbonylation Occurs in the Assembly Line of Barbamide for Skeleton Editing. *J. Am. Chem. Soc.* **145**, 5017–5028 (2023).

46. J. K. Brunson, S. M. K. McKinnie, J. R. Chekan, J. P. McCrow, Z. D. Miles, E. M. Bertrand, V. A. Bielinski, H. Luhavaya, M. Oborník, G. J. Smith, D. A. Hutchins, A. E. Allen, B. S. Moore, Biosynthesis of the neurotoxin domoic acid in a bloom-forming diatom. *Science* **361**, 1356–1358 (2018).

47. Y. Perez-Riverol, J. Bai, C. Bandla, D. García-Seisdedos, S. Hewapathirana, S. Kamatchinathan, D. J. Kundu, A. Prakash, A. Frericks-Zipper, M. Eisenacher, M. Walzer, S. Wang, A. Brazma, J. A. Vizcaíno, The PRIDE database resources in 2022: a hub for mass spectrometry-based proteomics evidences. *Nucleic Acids Res.* **50**, D543–D552 (2022).

48. W. W. Driscoll, N. J. Espinosa, O. T. Eldakar, J. D. Hackett, ALLELOPATHY AS AN EMERGENT, EXPLOITABLE PUBLIC GOOD IN THE BLOOM-FORMING MICROALGA *PRYMNESIUM PARVUM*. *Evolution* **67**, 1582–1590 (2013).

49. R. R. L. Guillard, P. E. Hargraves, *Stichochrysis immobilis* is a diatom, not a chrysophyte. *Phycologia* **32**, 234–236 (1993).

50. F. J. Sedlazeck, P. Rescheneder, M. Smolka, H. Fang, M. Nattestad, A. von Haeseler, M. C. Schatz, Accurate detection of complex structural variations using single-molecule sequencing. *Nat. Methods* **15**, 461–468 (2018).

51. J. T. Robinson, H. Thorvaldsdóttir, W. Winckler, M. Guttman, E. S. Lander, G. Getz, J. P. Mesirov, Integrative genomics viewer. *Nat. Biotechnol.* **29**, 24–26 (2011).

52. A. R. Quinlan, I. M. Hall, BEDTools: a flexible suite of utilities for comparing genomic features. *Bioinformatics* **26**, 841–842 (2010).

53. R. Pracana, A. Priyam, I. Levantis, R. A. Nichols, Y. Wurm, The fire ant social chromosome supergene variant Sb shows low diversity but high divergence from SB. *Mol. Ecol.* **26**, 2864–2879 (2017).

54. W. J. Kent, BLAT—The BLAST-Like Alignment Tool. *Genome Res.* **12**, 656–664 (2002).

55. O. Tange, GNU Parallel 20230522 ('Charles'), Zenodo (2023); https://doi.org/10.5281/zenodo.7958356.

56. G. Gremme, S. Steinbiss, S. Kurtz, GenomeTools: A Comprehensive Software Library for Efficient Processing of Structured Genome Annotations. *IEEE/ACM Trans. Comput. Biol. Bioinform.* **10**, 645–656 (2013).

57. P. Di Tommaso, M. Chatzou, E. W. Floden, P. P. Barja, E. Palumbo, C. Notredame, Nextflow enables reproducible computational workflows. *Nat. Biotechnol.* **35**, 316–319 (2017).

58. T. Lassmann, O. Frings, E. L. L. Sonnhammer, Kalign2: high-performance multiple alignment of protein and nucleotide sequences allowing external features. *Nucleic Acids Res.* **37**, 858–865 (2009).

59. N. P. Brown, C. Leroy, C. Sander, MView: a web-compatible database search or multiple alignment viewer. *Bioinformatics* **14**, 380–381 (1998).

60. Zymo Research, RNA Clean & Concentrator -25 Protocol. https://files.zymoresearch.com/protocols/_r1017_r1018_rna_clean_concentrator-25.pdf.

61. A. M. Bolger, M. Lohse, B. Usadel, Trimmomatic: a flexible trimmer for Illumina sequence data. *Bioinformatics*, 1–7 (2014).

62. D. Kim, J. M. Paggi, C. Park, C. Bennett, S. L. Salzberg, Graph-based genome alignment and genotyping with HISAT2 and HISAT-genotype. *Nat. Biotechnol.* **37**, 907–915 (2019).

63. M. Pertea, G. M. Pertea, C. M. Antonescu, T.-C. Chang, J. T. Mendell, S. L. Salzberg, StringTie enables improved reconstruction of a transcriptome from RNA-seq reads. *Nat. Biotechnol.* **33**, 290–295 (2015).

64. C. Camacho, G. Coulouris, V. Avagyan, N. Ma, J. Papadopoulos, K. Bealer, T. L. Madden, BLAST+: architecture and applications. *BMC Bioinformatics* **10**, 421 (2009).

65. J. M. Palmer, J. Stajich, Funannotate v1.8.1: Eukaryotic genome annotation, Zenodo (2020); https://doi.org/10.5281/zenodo.4054262.

66. G. S. C. Slater, E. Birney, Automated generation of heuristics for biological sequence comparison. *BMC Bioinformatics* **6**, 31 (2005).

67. R. Craig, R. C. Beavis, TANDEM: matching proteins with tandem mass spectra. *Bioinformatics* **20**, 1466–1467 (2004).

68. J. K. Eng, T. A. Jahan, M. R. Hoopmann, Comet: An open-source MS/MS sequence database search tool. *PROTEOMICS* **13**, 22–24 (2013).

69. H. Barsnes, M. Vaudel, SearchGUI: A Highly Adaptable Common Interface for Proteomics Search and de Novo Engines. *J. Proteome Res.* **17**, 2552–2555 (2018).

70. J. E. Elias, S. P. Gygi, "Target-Decoy Search Strategy for Mass Spectrometry-Based Proteomics" in *Proteome Bioinformatics*, S. J. Hubbard, A. R. Jones, Eds. (Humana Press, Totowa, NJ, 2010; https://doi.org/10.1007/978-1-60761-444-9_5)*Methods in Molecular Biology*$^{TM}$, pp. 55–71.

71. W. Shen, S. Le, Y. Li, F. Hu, SeqKit: A Cross-Platform and Ultrafast Toolkit for FASTA/Q File Manipulation. *PLOS ONE* **11**, e0163962 (2016).

72. M. Vaudel, J. M. Burkhart, R. P. Zahedi, E. Oveland, F. S. Berven, A. Sickmann, L. Martens, H. Barsnes, PeptideShaker enables reanalysis of MS-derived proteomics data sets. *Nat. Biotechnol.* **33**, 22–24 (2015).

73. D. W. Powell, C. M. Weaver, J. L. Jennings, K. J. McAfee, Y. He, P. A. Weil, A. J. Link, Cluster Analysis of Mass Spectrometry Data Reveals a Novel Component of SAGA. *Mol. Cell. Biol.* **24**, 7249–7259 (2004).

74. G. M. Kurtzer, cclerget, M. Bauer, I. Kaneshiro, D. Trudgian, D. Godlove, hpcng/singularity: Singularity, Zenodo (2021); https://doi.org/10.5281/zenodo.4667718.

75. G. M. Kurtzer, V. Sochat, M. W. Bauer, Singularity: Scientific containers for mobility of compute. *PLOS ONE* **12**, e0177459 (2017).

76. T. R. Fallon, photocyte/interproscan_parallel: v1.0.0, Zenodo (2023); https://doi.org/10.5281/zenodo.8432691.

77. V. Zulkower, S. Rosser, DNA Features Viewer: a sequence annotation formatting and plotting library for Python. *Bioinformatics* **36**, 4350–4352 (2020).

78. M. C. Kim, J. M. Winter, R. Cullum, A. J. Smith, W. Fenical, Expanding the Utility of Bioinformatic Data for the Full Stereostructural Assignments of Marinolides A and B, 24- and 26-Membered Macrolactones Produced by a Chemically Exceptional Marine-Derived Bacterium. *Mar. Drugs* **21**, 367 (2023).

79. K. D. Bauman, V. V. Shende, P. Y.-T. Chen, D. B. B. Trivella, T. A. M. Gulder, S. Vellalath, D. Romo, B. S. Moore, Enzymatic assembly of the salinosporamide γ-lactam-β-lactone anticancer warhead. *Nat. Chem. Biol.*, 1–9 (2022).

80. D. Khare, W. A. Hale, A. Tripathi, L. Gu, D. H. Sherman, W. H. Gerwick, K. Håkansson, J. L. Smith, Structural Basis for Cyclopropanation by a Unique Enoyl-Acyl Carrier Protein Reductase. *Structure* **23**, 2213–2223 (2015).

81. J. Jumper, R. Evans, A. Pritzel, T. Green, M. Figurnov, O. Ronneberger, K. Tunyasuvunakool, R. Bates, A. Žídek, A. Potapenko, A. Bridgland, C. Meyer, S. A. A. Kohl, A. J. Ballard, A. Cowie, B. Romera-Paredes, S. Nikolov, R. Jain, J. Adler, T. Back, S. Petersen, D. Reiman, E. Clancy, M. Zielinski, M. Steinegger, M. Pacholska, T. Berghammer, S. Bodenstein, D. Silver, O. Vinyals, A. W. Senior, K. Kavukcuoglu, P. Kohli, D. Hassabis, Highly accurate protein structure prediction with AlphaFold. *Nature*, 1–11 (2021).

82. M. Mirdita, K. Schütze, Y. Moriwaki, L. Heo, S. Ovchinnikov, M. Steinegger, ColabFold: making protein folding accessible to all. *Nat. Methods* **19**, 679–682 (2022).

83. M. van Kempen, S. S. Kim, C. Tumescheit, M. Mirdita, J. Lee, C. L. M. Gilchrist, J. Söding, M. Steinegger, Fast and accurate protein structure search with Foldseek. *Nat. Biotechnol.*, 1–4 (2023).

84. F. M. Ferroni, C. Tolmie, M. S. Smit, D. J. Opperman, Structural and Catalytic Characterization of a Fungal Baeyer-Villiger Monooxygenase. *PLOS ONE* **11**, e0160186 (2016).

85. C. R. Nicoll, G. Bailleul, F. Fiorentini, M. L. Mascotti, M. W. Fraaije, A. Mattevi, Ancestral-sequence reconstruction unveils the structural basis of function in mammalian FMOs. *Nat. Struct. Mol. Biol.* **27**, 14–24 (2020).

86. R. Al-Dhelaan, P. S. Russo, S. E. Padden, A. Amaya, D. W. Dong, Y.-O. You, Condensation-Incompetent Ketosynthase Inhibits trans-Acyltransferase Activity. *ACS Chem. Biol.* **14**, 304–312 (2019).

87. J. Masschelein, P. K. Sydor, C. Hobson, R. Howe, C. Jones, D. M. Roberts, Z. L. Yap, J. Parkhill, E. Mahenthiralingam, G. L. Challis, A dual transacylation mechanism for polyketide synthase chain release in enacyloxin antibiotic biosynthesis. *Nat. Chem.* **11**, 906–912 (2019).

88. S. Smith, S.-C. Tsai, The type I fatty acid and polyketide synthases: a tale of two megasynthases. *Nat. Prod. Rep.* **24**, 1041–1072 (2007).

89. J. Zheng, A. T. Keatinge-Clay, The status of type I polyketide synthase ketoreductases. *MedChemComm* **4**, 34–40 (2013).

90. T. M. McCullough, A. Dhar, D. L. Akey, J. R. Konwerski, D. H. Sherman, J. L. Smith, Structure of a modular polyketide synthase reducing region. *Structure* **31**, 1109-1120.e3 (2023).

91. C. Pockrandt, M. Alzamel, C. S. Iliopoulos, K. Reinert, GenMap: ultra-fast computation of genome mappability. *Bioinformatics* **36**, 3687–3692 (2020).

92. A. M. Kozlov, D. Darriba, T. Flouri, B. Morel, A. Stamatakis, RAxML-NG: a fast, scalable and user-friendly tool for maximum likelihood phylogenetic inference. *Bioinformatics* **35**, 4453–4455 (2019).

93. J. Huerta-Cepas, F. Serra, P. Bork, ETE 3: Reconstruction, Analysis, and Visualization of Phylogenomic Data. *Mol. Biol. Evol.* **33**, 1635–1638 (2016).

94. Z. Yin, J. S. Dickschat, Cis double bond formation in polyketide biosynthesis. *Nat. Prod. Rep.* **38**, 1445–1468 (2021).
